# Discovery and Optimization of LAG-3-Targeted Small Molecules via DNA-Encoded Chemical Library (DEL) Screening for Cancer Immunotherapy

**DOI:** 10.1101/2025.08.06.668839

**Authors:** Somaya A. Abdel-Rahman, Laura Calvo-Barreiro, Nelson García Vázquez, Hossam Nada, Moustafa T. Gabr

## Abstract

Lymphocyte activation gene-3 protein (LAG-3) is an immune checkpoint receptor that promotes T cell exhaustion and immune evasion in cancer. While antibody-based LAG-3 inhibitors have reached the clinic, small molecule modulators remain unexplored. Here, we report compound **11**, the most potent small molecule LAG-3 inhibitor to date. Identified via a 4.2-billion compound DNA-encoded chemical library (DEL) screen, compound **11** binds LAG-3 with submicromolar affinity and disrupts the LAG-3/MHCII interaction. Molecular modeling suggests direct antagonism at the LAG-3/MHCII interface with potential allosteric effects. In functional assays, compound **11** enhances IFN-γ secretion and promotes tumor cell killing in co-cultures of PBMCs and cancer cells. Importantly, compound **11** also exhibits favorable pharmacokinetics. These findings support the development of small molecule LAG-3 inhibitors as immunotherapeutic agents and provide a foundation for further optimization.

**Table of Contents artwork:** 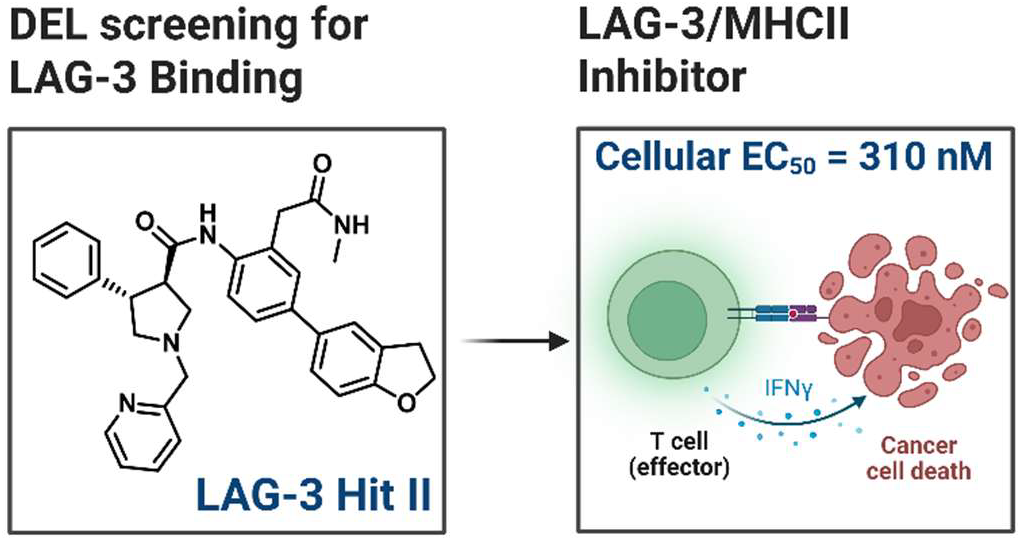

## INTRODUCTION

The development of immune checkpoint blockade (ICB) therapies has led to major breakthroughs in cancer treatment, yielding substantial clinical benefits and extended survival for certain patient populations.^1-3^ This progress has been primarily driven by monoclonal antibodies (mAbs) that inhibit immune checkpoints, particularly programmed cell death protein 1 (PD-1) and cytotoxic T-lymphocyte-associated protein 4 (CTLA-4).^4,5^ Despite the clinical success of ICB, a substantial subset of cancer patients exhibit primary resistance or develop acquired resistance over time, limiting the overall efficacy of these therapies.^6-9^ To address these challenges, research has increasingly focused on targeting additional immune checkpoints—such as lymphocyte activation gene 3 protein (LAG-3), T-cell immunoglobulin mucin-3 (TIM-3), and V-domain immunoglobulin suppressor of T-cell activation (VISTA)—to expand the therapeutic benefits of ICB to a broader range of cancer patients.^6-9^

LAG-3 is an inhibitory receptor highly expressed on exhausted T cells and represents a promising target for immunotherapy.^10-15^ LAG-3 operates in conjunction with other immune checkpoints, including PD-1 and CTLA-4, to augment the suppressive activity of regulatory T cells, thereby facilitating immune tolerance through antigen-presenting cells (APCs).^16^ LAG-3^+^ T cells interact with ligands, including major histocompatibility complex class II (MHCII) and fibrinogen-like protein 1 (FGL1), leading to inhibition of T-cell activation and cytokine production by indirectly interfering with T-cell receptor (TCR) signaling.^10-13^ Interestingly, the engagement of LAG-3 with MHCII—its primary ligand—initiates intracellular signaling in dendritic cells, activating phospholipase C γ2, p72syk, PI3K/AKT, p42/44, and p38 protein kinases.^17^ Furthermore, FGL1 was recently identified as a LAG-3 ligand that suppresses antigen-specific T-cell responses.^18,19^

LAG-3 expression is notably elevated in tumor-infiltrating lymphocytes (TILs) across various malignancies.^20-23^ While numerous studies have demonstrated that dual inhibition of PD-1 and LAG-3 on CD8^+^ and CD4^+^ TILs significantly enhances antitumor activity in preclinical models of ovarian cancer, colon adenocarcinoma, and melanoma.^24-26^ A key clinical study, RELATIVITY-047, reported that combined blockade of LAG-3 and PD-1 using mAbs resulted in a median progression-free survival of 10.1 months in metastatic melanoma patients, compared to 4.6 months with PD-1 inhibition alone.^27^ As a result, the combination therapy—relatlimab (anti-LAG-3) and nivolumab (anti-PD-1)—was granted approval by the US Food and Drug Administration (FDA) in 2022 for the treatment of metastatic melanoma.^27^

Despite more than 100 ongoing clinical trials evaluating LAG-3-targeting therapies, current strategies rely primarily on mAbs with no available small molecule inhibitors.^28^ While highly specific and potent, mAbs present several challenges, including limited tumor penetration, high manufacturing costs, and the risk of triggering immunogenic responses.^29-31^ Additionally, their extended half-life can lead to prolonged on-target immune-related adverse events (irAEs).^32,33^ In contrast, small molecules offer advantages such as oral bioavailability, improved tumor penetration, and greater flexibility in optimizing pharmacokinetics (PK).^34,35^ These properties enable more adaptable dosing regimens that may help mitigate irAEs associated with mAb-based therapies. Furthermore, small molecules can be more easily engineered for intracellular targets, broadening their potential applications in immuno-oncology. Consequently, the development of small molecule LAG-3 inhibitors could provide a viable therapeutic alternative, expanding combination treatment options and enhancing translational potential in solid tumor immunotherapy.

Our lab has been at the forefront of discovering small-molecule immune checkpoint inhibitors, including first-in-class inhibitors of LAG-3, TIM-3, VISTA, and ICOS (inducible costimulator of T cells).^36-44^ We have employed random, focused, and virtual screening of commercially available small-molecule libraries.^36-44^ Specifically for LAG-3, we identified first-in-class small-molecule inhibitors through high-throughput screening (HTS) and pharmacophore-based screening.^36-38^ However, the top hit compounds from our previous work exhibited moderate LAG-3 binding affinity in the micromolar range and/or failed to inhibit the LAG-3/MHCII interaction.^36-38^ These findings underscore the need for alternative screening platforms with access to a broader chemical space.

DNA-encoded chemical library (DEL) technology is a powerful platform for identifying small molecules that reversibly bind target proteins, offering access to a significantly larger chemical space than traditional HTS campaigns.^45-47^ It employs DNA-tagged compound libraries, enabling efficient screening of vast chemical spaces.^45-47^ A DEL is constructed by attaching an initial chemical building block to an oligonucleotide linker, followed by iterative cycles of DNA barcode extension and chemical conjugation.^45,46^ For instance, incorporating 1,000 unique building blocks per cycle over three iterations can theoretically yield a billion distinct compounds. This high-throughput approach enhances screening efficiency and cost-effectiveness by generating and analyzing extensive libraries in a single experiment. DEL compounds are screened in pooled reactions, requiring minimal target protein.^45,46^ High-affinity binders selectively interact with immobilized proteins on beads, while weaker or non-binding compounds are washed away. Strong binders are then eluted and identified through DNA sequencing, akin to barcode scanning, enabling rapid hit discovery.^45,46^

In this study, we employed DEL screening to identify novel LAG-3-targeted hit compounds (Figure 1). Through structural optimization of the LAG-3 hit, we developed the most potent small molecule LAG-3 inhibitor reported to date, capable of disrupting the LAG-3/MHCII interaction in co-culture assays at submicromolar concentrations (Figure 1). In functional assays, our optimized LAG-3 inhibitor enhanced IFN-γ secretion and promoted tumor cell killing in co-culture assays, confirming its immunomodulatory activity. Our computational analysis of the LAG-3 binding site for the lead compound provides valuable insights for future optimization, while assessment of the PK profile ensures the feasibility of in vivo evaluation.

**Figure 1.**
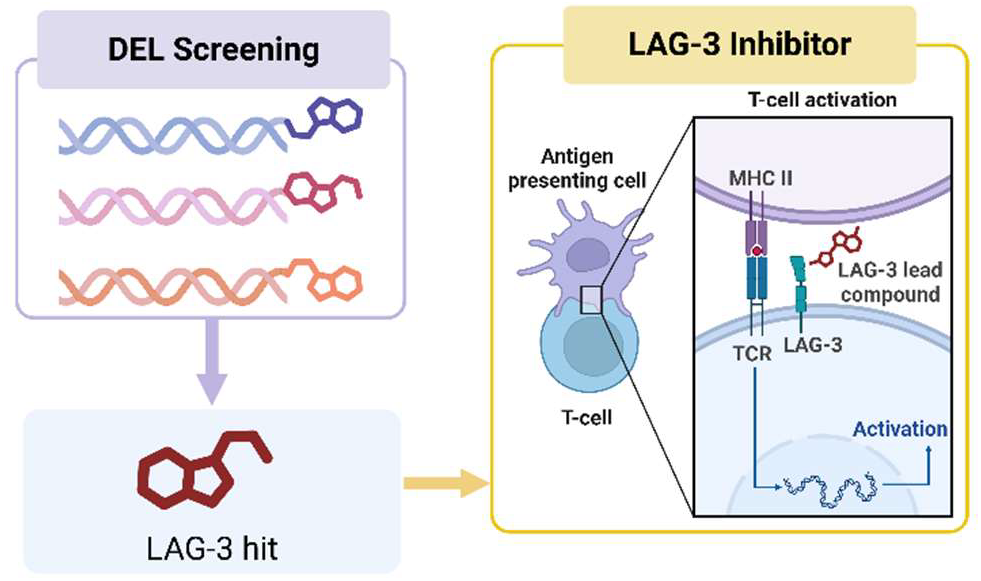
DEL-screening based development of LAG-3 Inhibitors. DEL screening identified a LAG-3 binder, optimized into an inhibitor that enhances T-cell activation.

## RESULTS AND DISCUSSION

### DEL Screening

We performed affinity selection screening (Figure 2A) of 4.2 billion compounds from the DELopen kit [DNA-Encoded Libraries for Academic Users, WuXi AppTec], for LAG-3 by capturing LAG-3 on Ni-NTA magnetic beads for binding affinity screening. After incubation with the DELopen library, beads conjugated to LAG-3 or control empty beads were magnetically pulled down, washed, and heated to release the bound library molecules. We selected the top compounds after removing NTC (non-target control) signals (C4) from each target condition (C1, C2, and C3: human LAG-3 protein), applying an enrichment score (Figure 2B) threshold above 100 for each target condition, and considering only those consistently present across all biological replicates (ABC area, Figure 2A). Structural patterns emerged across multiple libraries, consistently exhibiting strong signal performance across all biological replicates (C1, C2, and C3). The final selection of the two top compounds (**LAG-3 Hit I** and **LAG-3 Hit II**, Figure 2C) was based on the aforementioned criteria, prioritizing structural diversity, the most represented chemotypes in the data analysis, and compounds that passed the PAINS filter to exclude promiscuous binders.^48^

**Figure 2.**
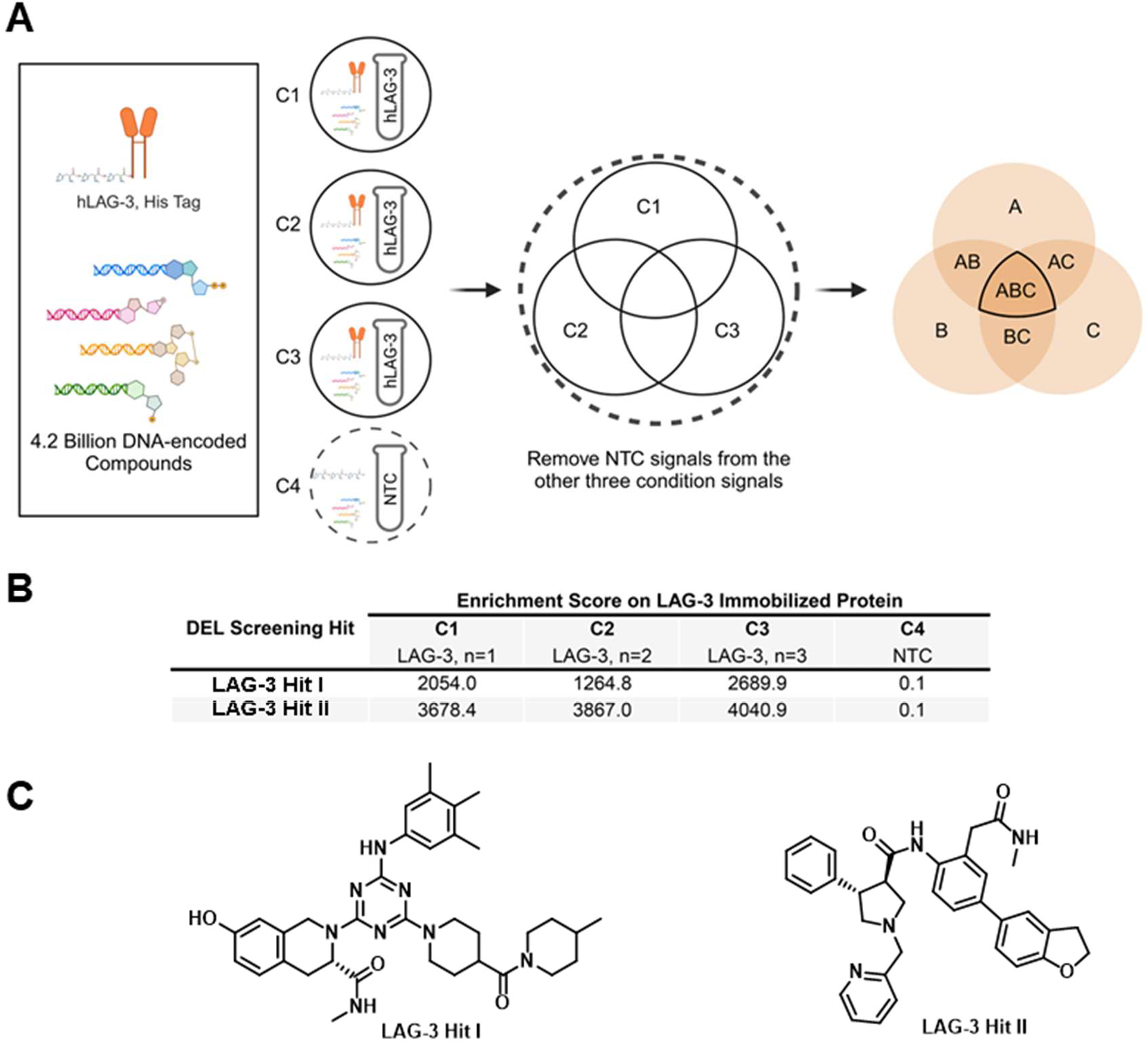
Selection of the top two small molecule LAG-3 binders discovered through DEL screening. **A**. A total of 27 libraries, comprising 4.2 billion DNA-encoded compounds, were incubated with human LAG-3 protein immobilized on an affinity matrix in triplicate (C1, C2, C3) or with the affinity matrix alone as a non-target control (NTC, C4). Following affinity selection, bound small molecules were sequenced to decode their DNA tags. After removing NTC signals (C4) and applying selection criteria, compounds consistently detected across all replicates (ABC area) were identified as potential human LAG-3 binders. **B**. Enrichment scores of the top two small molecules (**LAG-3 Hit I** and **LAG-3 Hit II**) selected for binding to human LAG-3 protein following affinity selection. **C**. Chemical Structures of **LAG-3 Hit I** and **LAG-3 Hit II** compounds.

After procuring **LAG-3 Hit I** and **LAG-3 Hit II** from WuXi AppTec (see Figures S1-S10 for NMR, LCMS, mass spectra, HPLC purity, and chiral supercritical fluid chromatography (SFC)), we initially evaluated their binding affinity to LAG-3 using microscale thermophoresis (MST). To validate our MST protocol, we measured the estimated dissociation rate constant (*K*_*D*_) value for FGL1 binding to LAG-3, yielding an estimated value of 5.76 nM. Screening **LAG-3 Hit I** and **LAG-3 Hit II** using this validated protocol revealed that **LAG-3 Hit I** exhibited minimal affinity for LAG-3 (Figure S11), **LAG-3 Hit II** bound LAG-3 with a *K*_*D*_ of 4.32 ± 1.02 μM (Figure S12). Additionally, we developed a surface plasmon resonance (SPR) assay for LAG-3 using FGL1 protein and relatlimab (anti-LAG-3 mAb) as positive controls. These experiments confirmed that the immobilized LAG-3 protein remained stable and active on the Sensor Chip CAP, which we subsequently used for smallmolecule screening. Using this approach, we further confirmed the binding of **LAG-3 Hit II** to LAG-3, determining a *K*_*D*_ value of 2.97 ± 1.69 µM (Figure 3). Notably, the binding affinity of **LAG-3 Hit II** measured by SPR was consistent with our MST results, highlighting the robustness of our assay methodologies and supporting **LAG-3 Hit II** as a valid LAG-3 binder. To demonstrate cellular target engagement, we employed a Cellular Thermal Shift Assay (CETSA) using Raji-hLAG-3 cells, following a protocol previously developed by our group for cellular target engagement of small molecule LAG-3 binders.^38^ As shown in Figure S14, treatment with **LAG-3 Hit II** resulted in a reproducible thermal shift of approximately 5.1 °C, indicative of specific intracellular binding to full-length LAG-3 under physiologically relevant conditions. This result complements our biophysical assays (SPR and MST), providing a robust, multi-tiered validation of LAG-3 engagement by **LAG-3 Hit II**.

**Figure 3.**
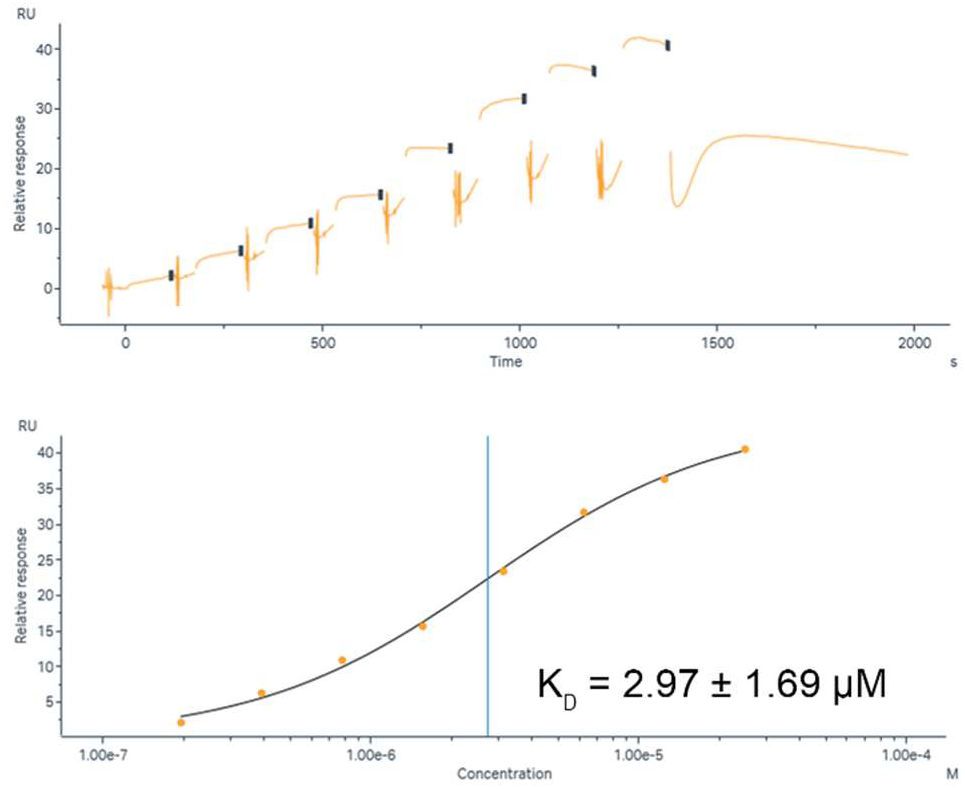
SPR binding curve of LAG-3 Hit II to LAG-3. Serial dilutions of **LAG-3 Hit II** (25 µM to 195.31 nM, 2-fold dilution, 8 steps) were injected onto a Sensor Chip CAP with immobilized human LAG-3 protein using single-cycle kinetics. Graphs show one representative experiment out of the three independent experiments that were performed.

### Optimization of the LAG-3 Hit

In our preliminary optimization of **LAG-3 Hit II**, we identified a key core fragment of **LAG-3 Hit II** (pyrrolidine-3-carboxamide core, compound **A**, Scheme 1). To explore its preliminary structure-activity relationship (SAR), we coupled the pyrrolidine-3-carboxylic acid derivative (compound **A**) with various aromatic amines (**B**) using HBTU (Hexafluorophosphate Benzotriazole Tetramethyl Uronium) in DMF and in the presence of triethylamine (Scheme 1). The chemical structures of the synthesized compounds (compound **1-11**), as well as their LAG-3 binding affinity using MST, are shown in Table 1. The structural characterization of the synthesized (compound **1-11**) is presented in Figures S15-45. The MST binding curves of compound **1-11** are depicted in Figures S46-S54.

**Scheme 1.**
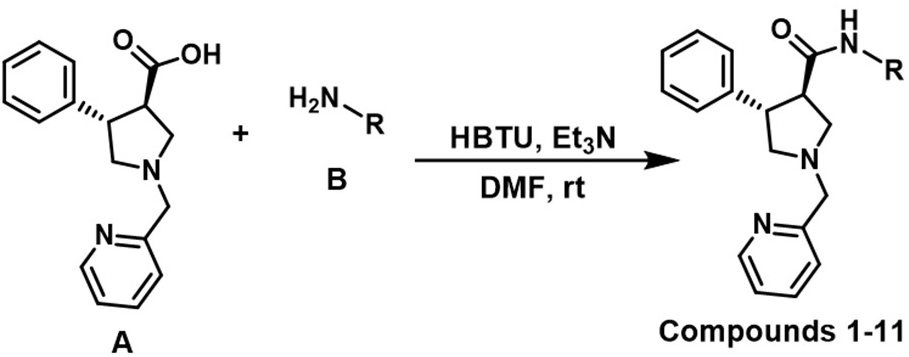
Synthesis of compound **1-11** via coupling of compound **A** with various aromatic primary amines (**B**).

**Table 1.**
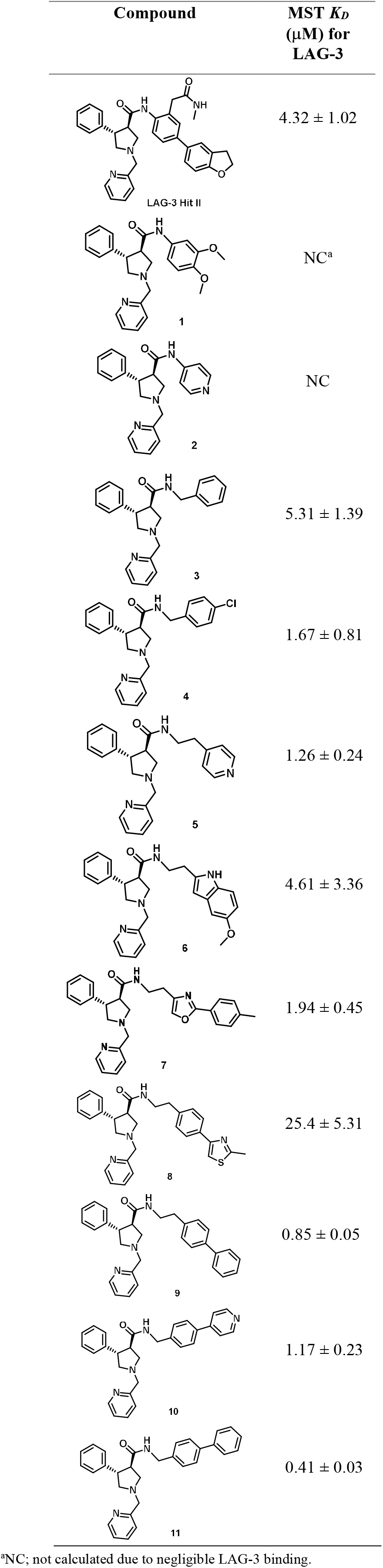
Structural analogs of LAG-3 Hit II (compounds 1-11) and their LAG-3 binding affinity using MST.

As shown in Table 1, structural simplification by removing the 2,3-dihydrobenzofuran-5-yl moiety from **LAG-3 Hit II** and coupling with substituted anilines or aminopyridines resulted in negligible LAG-3 binding affinity (compounds **1** and **2**). Aiming to enhance LAG-3 binding, we investigated the introduction of various linkers between the pyrrolidine-3-carboxamide core (**A**) and the aromatic moiety of the primary amine (**B**). Simultaneously, we explored preliminary structural modifications of the aromatic amine (**B**) to determine the impact of electronic and steric properties on binding. Using benzylamine as the aromatic amine yielded compound **3**, which exhibited a slight reduction in the LAG-3 binding affinity compared to **LAG-3 Hit II** (Table 1). However, replacing benzylamine with 4-chlorobenzylamine (compound **4**) resulted in an improvement in LAG-3 binding affinity, suggesting that the chlorine substituent enhances interactions within the binding pocket (Table 1).

Building on these findings, we explored the effect of incorporating an ethylene bridge between the pyrrolidine-3-carboxamide core (**A**) and the aromatic amine (**B**). Most of the resulting derivatives (compounds **5, 6, 7**, and **9**) exhibited *K*_*D*_ values below 5 µM, except for compound **8**, which had a *K*_*D*_ value of 25.4 ± 5.31 (Table 1). This indicates that the ethylene bridge generally contributed to maintaining or improving LAG-3 binding affinity. Among these derivatives, compound **9**, which incorporated a 1,1’-biphenyl fragment, demonstrated the highest potency, achieving a *K*_*D*_ of 0.85 ± 0.05 µM. This suggests that the biphenyl system enhances ligand-receptor interactions, potentially through increased hydrophobic contacts. Based on the promising LAG-3 binding affinity of compound **9**, we synthesized additional derivatives featuring a methylene linker and substituted aromatic fragments, including 4-phenylpyridyl (compound **10**) and 1,1’-biphenyl (compound **11**). Among these, compound **11** emerged as the most potent LAG-3 binder, with a *K*_*D*_ value of 0.41 ± 0.03 µM, representing a >10-fold improvement in affinity compared to **LAG-3 Hit II** (Table 1). These results underscore the critical role of both the linker and the nature of the aromatic moiety in optimizing LAG-3 binding.

### Preliminary Evaluation of Compound 11

Among our optimized compounds, compound **11** emerged as the most potent LAG-3 binder. In our pursuit to identify the functional activity of compound **11**, we further assessed its efficacy as a LAG3/MHCII inhibitor using a cell-based bioassay. This approach aimed to validate the compound’s ability to block the LAG-3/MHCII interaction in a biological context, providing insight into its potential as a therapeutic candidate for immune modulation. We employed a bioluminescent cell-based assay developed by Promega, designed specifically to evaluate the ability of agents to disrupt the LAG-3/MHCII interaction. This assay uses two distinct cell lines: an MHCII-positive human cell line and NanoLuc^®^ (NL) luciferase reporter/Jurkat LAG-3 cells. When co-cultured, the LAG-3 protein on the Jurkat cells inhibits TCR pathway activation, resulting in reduced luminescence. The introduction of a LAG-3/MHCII inhibitor disrupts LAG-3’s binding to MHCII, thereby restoring TCR activation and generating a luminescent signal. This signal is quantifiable in a dose-dependent manner by the addition of Bio-Glo^™^ Reagent and measurement with a luminometer. As demonstrated in Figure 4, compound **11** exhibited a dose-dependent increase in luminescence signal in the Promega LAG-3/MHCII blockade assay, confirming its ability to inhibit the interaction between LAG-3 and MHCII on the cell surface. The EC_50_ value of compound **11** in this assay was determined to be 0.31 ± 0.09 μM (Figure 4). Remarkably, compound **11** is the first small molecule LAG-3 inhibitor to demonstrate disruption of the LAG-3/MHC II interaction in a cellular assay at submicromolar concentrations.

**Figure 4.**
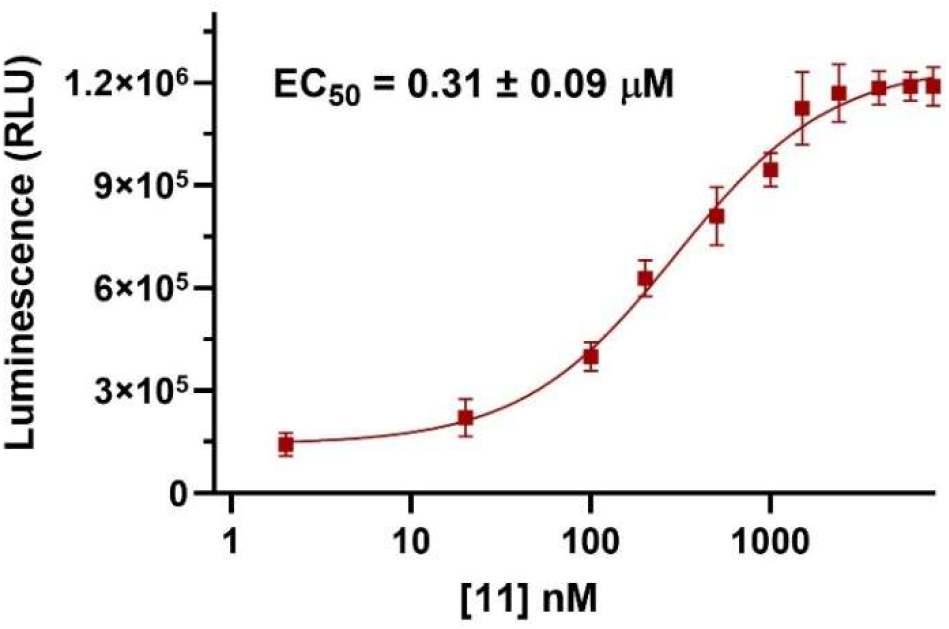
Dose-dependent response of compound **11** in the LAG-3/MHCII blockade assay from Promega. The luminescence was measured upon co-culturing LAG-3 effectors and MHCII APC cells in the presence of increasing concentrations of compound **11**. Error bars represent standard deviation (n = 3).

To evaluate the therapeutic potential of compound **11** as a LAG-3/MHCII inhibitor, we adapted a model used in the study of acute myeloid leukemia (AML) immunosuppression. In this model,^49^ peripheral blood mononuclear cells (PBMCs) from healthy donors were co-cultured THP1 AML cell line. To evaluate the impact of the LAG-3/MHCII inhibitor on immune responses, we focused on assessing IFN-γ secretion and the killing activity of THP-1 AML cells. The level of IFN-γ was measured as an indicator of T cell activation. Additionally, cell viability was assessed through 7-AAD/CFSE assays to determine the inhibitor’s effect on immune cell-mediated killing of THP-1 cells, offering insights into its potential for modulating AML-induced immunosuppression. The addition of compound **11** (10 μM) to PBMCs co-cultured with THP-1 AML cells resulted in a significant increase in IFN-γ secretion, mirroring the effect observed with relatlimab (Figure 5A). Additionally, compound **11** (10 μM) enhanced AML cell killing at 24 h, indicating that LAG-3 inhibition effectively potentiates immune responses against AML (Figure 5B). These findings suggest that targeting LAG-3 via compound **11** could be a promising strategy for enhancing anti-AML immunity and may provide therapeutic benefits when combined with other treatments.

**Figure 5.**
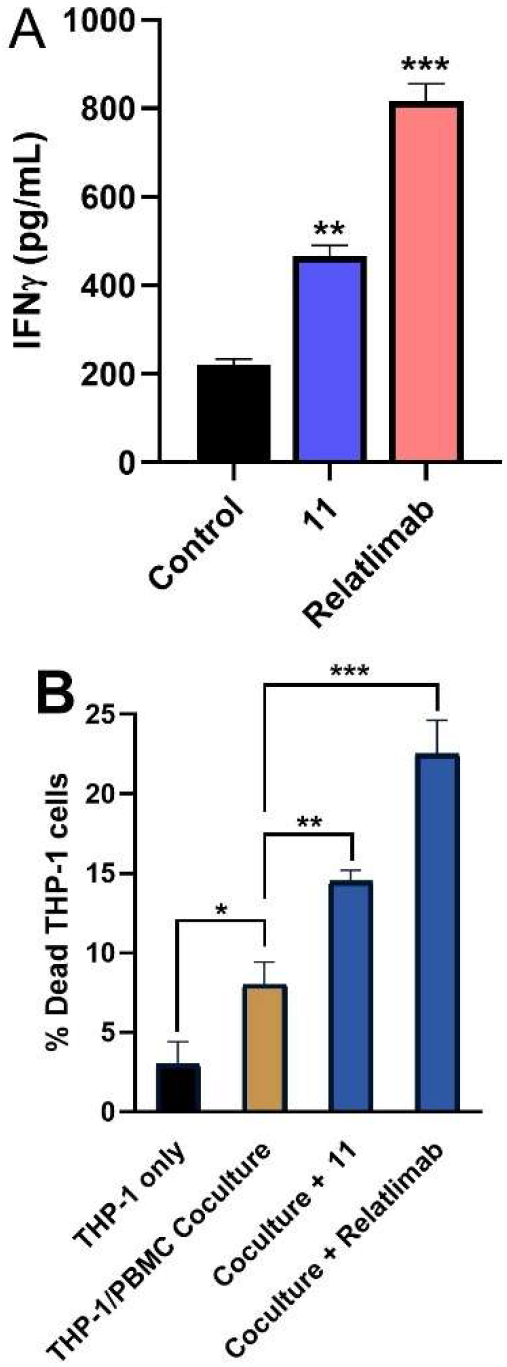
**A**. Production of IFNγ from PBMCs upon co-culturing with THP-1 cells in the absence and presence of relatilmab (100 µg/ml) and compound **11** (10 μM), ^**^ *p* < 0.01, ^***^ *p* < 0.001 in comparison to control. **B**. The % of dead THP-1 cells as assessed by 7-AAD/CFSE assay in the co-culture assay of PBMCs and THP-1 cells in the absence and presence of relatilmab (100 µg/ml) and compound **11** (10 μM), ^*^ *p* < 0.05, ^**^ *p* < 0.01, and ^***^ *p* < 0.001. Error bars represent standard deviation (n = 3).

Although compound **11** significantly enhanced IFN-γ secretion and promoted tumor cell killing in PBMC/THP-1 co-cultures, the magnitude of cytotoxicity was modest—similar to what was observed with the anti-LAG-3 antibody relatlimab. This is consistent with previous studies demonstrating that LAG-3 blockade alone is often insufficient to elicit strong effector responses and may require co-targeting of additional checkpoints, such as PD-1 or TIM-3, for optimal therapeutic benefit.^11-13^ These findings suggest that compound **11** may be most effective in combination immunotherapy regimens and underscore the importance of further evaluation in more physiologically relevant models. While the current study focuses on the in vitro optimization and characterization of compound **11**, future follow-up studies will include in vivo pharmacokinetic profiling and anti-tumor efficacy evaluation to further define its translational potential.

To validate the reproducibility and generalizability of compound **11**’s immune-enhancing effects, we extended the PBMC co-culture assay to two additional tumor cell lines. First, we used Kasumi-1, an independent AML cell line, to replicate the results obtained with THP-1. Treatment with compound **11** significantly increased IFN-γ secretion and promoted Kasumi-1 cell killing, confirming its immune-activating function within hematologic malignancies (Figure S55). Second, to evaluate the effects in a solid tumor context, we cocultured A549 lung carcinoma cells with PBMCs in the presence of compound **11**. A marked increase in IFN-γ production and cancer cell cytotoxicity was observed, consistent with LAG-3 checkpoint blockade (Figure S56). These results demonstrate that compound **11** activates immune effector responses across both hematological and epithelial cancer models, supporting its translational potential.

### Computational Analysis of the Binding Mode of Compound 11 to LAG-3

Given that compound **11** is the most potent small molecule LAG-3 inhibitor reported to date, we sought to investigate its binding mode to gain molecular insights into its interaction with LAG-3. Understanding these interactions could provide a structural basis for further optimization and the design of next-generation LAG-3 inhibitors with enhanced affinity and specificity. A recent study reported the crystal structure of murine LAG-3 interactions with the MHC class II.^50^ The study reported the presence of a cluster of residues in LAG-3 which are responsible for attracting MHC II. As such a comparative analysis was carried out to compare human and mouse LAG-3 protein sequences (Figures S57-S60) which revealed a complex interplay of conservation and divergence across species. Hydrophobicity profiles demonstrate overall structural similarity between human and mouse LAG-3, with conserved hydrophobic and hydrophilic regions suggesting similar folding patterns. However, domain-specific sequence similarity reveals significant divergence in the Ig-like domain, potentially reflecting species-specific functional adaptations. Conversely, the transmembrane region remains highly conserved, indicating its critical role in membrane anchoring. This is further supported by the fluctuating similarity scores along the protein alignment, highlighting specific regions of higher and lower conservation.

A heatmap of amino acid frequencies (Figure S59) reinforces these findings, pinpointing conserved residues likely crucial for protein function and variable regions potentially contributing to species-specific differences. Collectively, these analyses suggested that while the core structural and functional elements of LAG-3 are maintained across species and that the hot spot residues occupy homologous structural regions despite differing linear positions. Building upon these findings, computational modeling was employed to investigate the potential proteinprotein complex between human LAG-3 and HLA-DRA, the alpha chain of the human MHC class II molecule HLA-DR. The modelling study yielded the LAG-3/HLA-DRA complex illustrated in Figures 6A and 6B.

**Figure 6.**
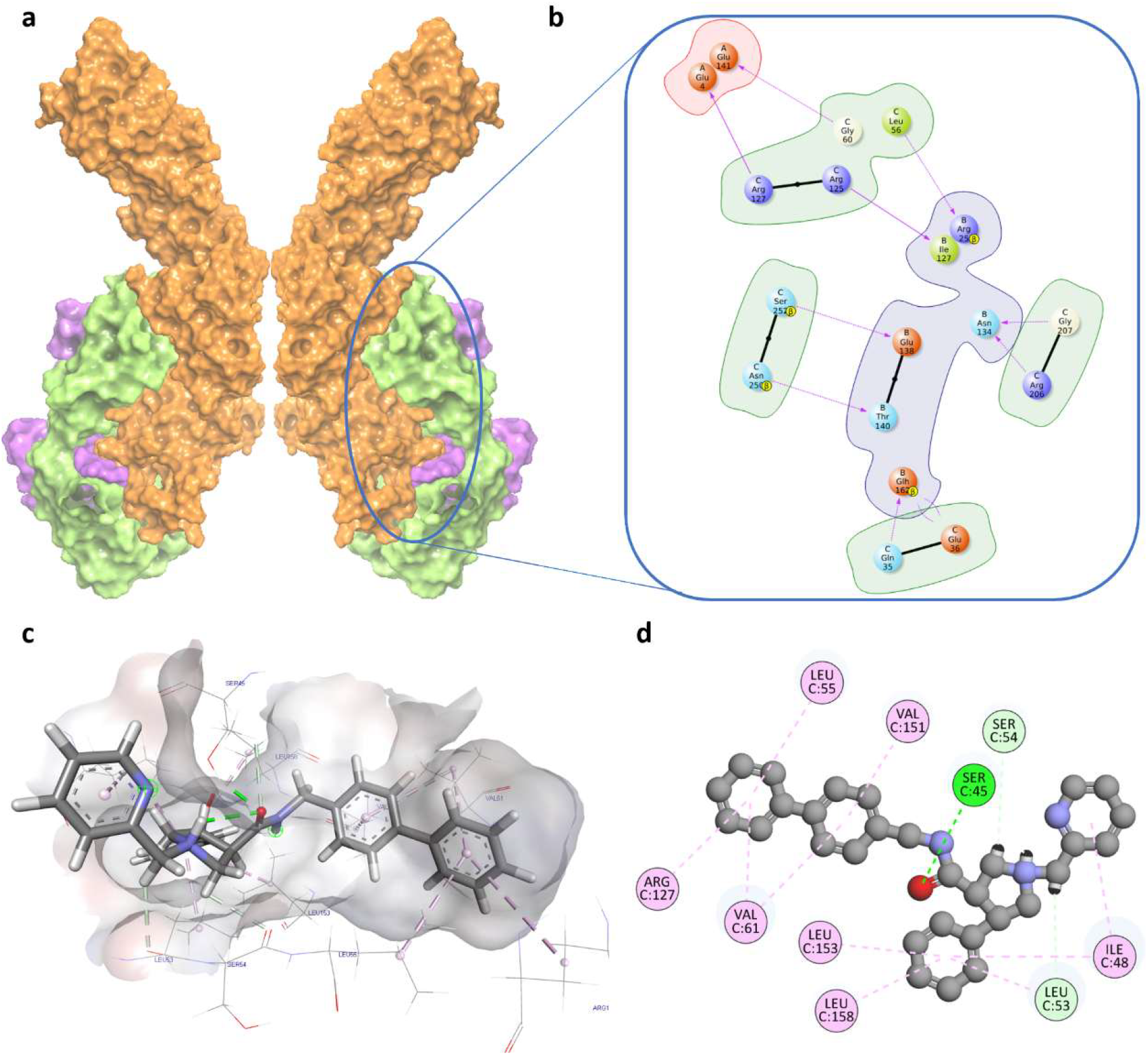
Molecular basis for LAG-3 and HLA-DRA and their inhibition using small molecules. A. Homology model of human LAG-3-HLA-DRA Complex: A 3D representation of the complex formed between LAG-3 (orange) and HLA-DRA (chain B: green, chain A: magenta). **B**. LAG-3-HLA-DRA Interaction Diagram: A schematic illustrating key molecular interactions between LAG-3 and HLA-DRA. **C**. Compound **11** Bound to LAG-3: A 3D structure showing the small molecule inhibitor compound **11** bound to the binding site on LAG-3. **D**. Compound **11**-LAG-3 Interaction Diagram: A 2D representation of the molecular interactions between compound **11** and LAG-3, highlighting key residues and binding forces.

The predicted protein-protein interaction interface revealed two contact points between the interacting proteins with LAG-3 Arg125, adopting a comparable binding orientation to that observed in the mouse LAG-3/MHC class II HLA-DRA subunit complex, thereby reinforcing the reliability of the generated model. The identified hot spot residues, including Gln35, Arg206, Arg125, Arg127, Asn250, and Ser252, formed a cluster of charged polar amino acids within two D1 loops. These residues were positioned opposite a charge-complementary surface of the MHC class II HLA-DRA subunit which contained key residues such as Glu4, Arg25, Ile127, Asn134, Glu138, Thr140, and Glu141.

Next, compound **11** was systematically docked into the identified hotspot residues of the LAG-3 at the site of LAG-3/MHC interface (Figures 6C and 6D) using the induced fit docking (IFD) protocol within Maestro Schrödinger Suite (v2021.2). This approach was selected to account for the ligand flexibility and conformational adjustments in the protein binding pocket. The molecular docking investigation revealed that compound **11** binds to LAG-3 at a deep groove. The ligand/protein interface involved a hydrogen bond with Ser45 and π–π interactions between the ligand and Ile48, Leu55, Val61, Arg127, Val151, Leu153, and Leu158 amino acid residues of the binding site. Two additional carbon-hydrogen interactions between compound **11** and Leu53 and Ser54 residue of LAG-3 helped to further stabilize the observed binding interaction.

Molecular dynamics (MD) simulations were performed on both the unbound LAG-3 protein and the LAG3/compound **11** complex to validate the molecular docking predictions. Contrary to expectations, the ligand-bound complex demonstrated higher Root Mean Square Deviation (RMSD) and Root Mean Square Fluctuation (RMSF) values compared to the unbound protein (Figures S61-S62). This observation suggests that compound **11** induces significant dynamic changes within the LAG-3 structure. Moreover, while LAG-3 inherently possesses a degree of flexibility, which is reflected by the high RMSD of the unbound LAG-3 (Figure S61), the observed increase in RMSD and RMSF upon the formation of the LAG-3/LIG complex indicates that the ligand does not simply restrict LAG3 to a single conformation. Instead, it appears to promote a dynamic equilibrium between multiple conformational states. This is further supported by the RMSF analysis, which revealed region-specific effects where the ligand induced increased flexibility in certain segments of LAG-3, while simultaneously stabilizing others. Together, these observations suggest a complex interplay between ligand binding and protein dynamics and the resulting eventual observed inhibitory effect.

The stability of the compound **11** complex during the MD simulation was assessed through interaction analysis, which demonstrated the persistent formation of at least one hydrogen bond throughout the simulation. This sustained interaction likely contributes to establishing the stable energy minima observed in the Gibbs free energy diagram (Figure S63). Analysis of the radius of gyration (Rg, Figure S64) revealed that the ligand does not significantly enhance the overall compactness or folding of LAG-3, which suggests that the ligand primarily influences the protein’s dynamic behavior rather than inducing a substantial folding transition. In summary, the MD simulations indicate that compound **11** interacts with LAG-3 in a dynamic manner, inducing both local stabilization and increased flexibility. This suggests a complex binding mode, potentially involving allosteric modulation, rather than a simple rigidification of the protein.

### Preliminary PK evaluation of Compound 11

To evaluate the in vivo potential and drug-like properties of the lead compound, compound **11**, we assessed its physicochemical characteristics and early ADME properties. The distribution coefficient at physiological pH (Log D_7.4_) was measured, along with the calculation of its topological polar surface area (TPSA) (Table 2). Both parameters fell within the established optimal ranges for drug-like compounds, with Log D_7.4_ values between −0.1 and 3.0 and TPSA values below 140 Å^2^ (Table 2). Kinetic solubility in 1% DMSO/PBS was >100 µM, and solubility in fasted state simulated intestinal fluid (FaSSIF) was similarly favorable (120 µM), supporting oral absorption. Compound **11** also exhibited high passive permeability in a PAMPA assay (*P*_*app*_ = 3.5 × 10^−6^ cm/s), consistent with drug-like permeability.

**Table 2.** In vitro PK profile of compound 11.

| PK parameter | Value |
| --- | --- |
| $\text{Log } D_{7.4}^a$ | 2.79 |
| TPSA | $121 \text{ \AA}^2$ |
| Kinetic Solubility<br>1% DMSO/PBS<br>[$\mu\text{M}$]<sup>b</sup> | $>100$ |
| Solubility in FaSSIF<br>[$\mu\text{M}$] | 120 |
| PAMPA $P_{app}$ [cm/s] | $3.5 \times 10^{-6}$ |
| Stability in simulated<br>gastric fluid (pH 1.2,<br>2h) | $>90\%$ remaining |
| Stability in simulated intestinal fluid (pH 6.8, 2h) | >85% remaining |
| t<sub>1/2</sub> [min] mouse liver microsomes<sup>c</sup> / CL<sub>int</sub><sup>d</sup> | 68.2 ± 7.5 / 28.3 ± 3.5 |
| t<sub>1/2</sub> [min] human liver microsomes<sup>c</sup> / CL<sub>int</sub><sup>d</sup> | 96.4 ± 5.3 / 17.1 ± 1.8 |
| t<sub>1/2</sub> [min] mouse plasma | >150 |
| t<sub>1/2</sub> [min] human plasma | >150 |
| t<sub>1/2</sub> [min] PBS | >500 |
| PPB human [%]<sup>c</sup> | 99.2 ± 0.83 |
| IC<sub>50</sub> against WI-38 [μM] | >100 |
| IC<sub>50</sub> against HS 27 [μM] | >100 |
| IC<sub>50</sub> against hERG [μM] | >100 |
| CYP3A4 inhibition (% at 10 μM) | 9.5% |
| CYP2D6 inhibition (% at 10 μM) | 8.1% |
| CYP2C9 inhibition (% at 10 μM) | 7.8% |
| CYP2C19 inhibition (% at 10 μM) | 8.3% |

<sup>a</sup>LogD (pH = 7.4) the distribution coefficient at physiological pH 7.4.
<sup>b</sup>Kinetic solubility measured in 1% DMSO/PBS (pH 7.4) by UV-vis spectrophotometry (n=3). <sup>c</sup>Values are mean ± SD (n=3). <sup>d</sup>Intrinsic clearance expressed in μL/(min × mg) protein. Values are mean ± SD (n=3).

Given the importance of metabolic stability in drug development, the metabolic stability of compound **11** was evaluated using both mouse and human liver microsomes. Notably, compound **11** exhibited relatively high stability in mouse liver microsomes, as reflected by a half-life (t_1/2_) of 68.2 minutes and an intrinsic hepatic clearance (Cl_int_) of 28.3 µL/mg/min (Table 2). Similarly, the stability of compound **11** in human microsomes was established with a t_1/2_ of 96.4 minutes and a Cl_int_ of 17.1 µL/mg/min. Furthermore, compound **11** demonstrated high stability (Table 2) in both mouse and human plasma, as well as in phosphate-buffered saline (PBS). Simulated gastric and intestinal fluid stability studies further confirmed the compound’s chemical stability across physiologically relevant pH conditions (Table 2). Plasma protein binding was high (PPB = 99.2%, Table 2). To further evaluate the translational potential of compound **11**, we performed in vitro–in vivo extrapolation (IVIVE) modeling using the well-stirred liver model. Based on the compound’s intrinsic clearance in human liver microsomes, high PPB, and physicochemical properties (LogD_7.4_ = 2.79), compound **11** is predicted to have low hepatic clearance and a moderate volume of distribution (~1.5 L/kg). These values suggest a prolonged systemic half-life, which is favorable for infrequent dosing. While additional in vivo studies are warranted, the IVIVE data support the compound’s developability and guide future pharmacokinetic optimization.

The safety of compound **11** was validated through testing on normal cell lines (WI-38 and HS 27 normal cells). Finally, the minimal inhibitory profile of compound **11** in the hERG assay confirmed the favorable safety profile (Table 2). Additionally, inhibition of the four major CYP450 isoforms was minimal, with <10% inhibition at 10 µM against CYP3A4, CYP2D6, CYP2C9, and CYP2C19. Overall, these findings indicate that compound **11** possesses promising drug-like properties, including favorable physicochemical characteristics, metabolic stability, and a satisfactory safety profile, making it a viable candidate for further in vivo evaluation and development

## CONCLUSIONS

This study introduces compound **11** as a potent small molecule LAG-3 inhibitor, derived through the optimization of a LAG-3 hit (**LAG-3 Hit II**). Importantly, compound **11** restores immune function in vitro, enhancing IFN-γ secretion and promoting tumor cell killing in co-culture assays—key indicators of T cell reinvigoration. Furthermore, PK profiling revealed favorable properties that support its potential for in vivo administration and further preclinical development.

Computational modeling and MD simulations revealed that compound **11** binds a conserved groove at the LAG-3/MHCII interface and induces dynamic conformational changes within the protein. These findings suggest a complex binding mode involving both direct antagonism and potential allosteric effects. Together, these results provide both mechanistic and translational foundations for the continued development of LAG-3-targeting small molecules. Future work will focus on elucidating in vivo efficacy, exploring synergistic combinations with PD-1/PD-L1 inhibitors, and structurally refining the compound **11** scaffold for enhanced specificity. This study establishes a platform for next-generation immunotherapies targeting LAG-3 and expands the drug discovery landscape for immune checkpoint modulation.

## EXPERIMENTAL

### DEL-Based Small Molecule Screening for LAG-3

A DNA-encoded chemical library comprising 4.2 billion compounds, provided by WuXi AppTec, was screened against the LAG-3 protein. Prior to the affinity selection process that would allow us to isolate those small molecule compounds that bound to our target protein, we performed a protein capture experiment to confirm that proper amounts of LAG-3 extracellular domain (ECD) protein (Cat. # 16498-H08H, Sino Biological) can be stably immobilized to the affinity matrix over the experimental protocol. Based on this initial experiment, we selected 1x PBS, 0.05% Tween-20, 0.1 mg/mL sssDNA, and 10 mM imidazole as the selection buffer; 1x PBS and 0.05% Tween-20 as the elution buffer; 5 µg of LAG-3 protein for each round of selection; and HisPur^™^ Ni-NTA Magnetic Beads (Cat. # 88831, Thermo Fisher, Waltham, MA, USA) as the affinity matrix. The product formulation of the PBS is: 0.144 g/L KH_2_PO_4_, 9.00 g/L NaCl, 0.795 g/L Na_2_HPO_4_ (anhydrous), pH 7.4.

Later, we performed the affinity selection protocol, including both the protein immobilization step and the rounds of selection, as described by the manufacturer. Each DELopen kit provides four identical copies of the 27 different libraries that comprise 4.2 billion compounds. Three out the four copies were used as biological replicates of LAG-3 protein while the last copy was used as a non-target control (NTC; HisPur^™^ Ni-NTA Magnetic Beads with no LAG-3 protein) to assess the amount of unspecific binders to the selected matrix.

The heated eluted samples obtained after the third round of selection were chosen as the post-selection samples based on the self-QC step recommended by the manufacturer. Briefly, this real-time, semi-quantitative step allows the researcher to quickly confirm that the amount of selected molecules in each experimental condition is less than 5×10^8^ copies (standard control) and that no cross-contamination is present. These post-selection samples were sent to WuXi AppTec who confirmed that all three post-selection samples (biological replicates) had met the sequencing quality standards and would go through the next generation sequencing (NGS) step to decode the DNA tags linked to each individual small molecule. The Enrichment Score value is calculated and reported by WuXi AppTec for every molecule that binds to the target protein. The Enrichment Score is statistically calculated from the copy number of that compound, considering the library size, NGS depth, and other normalization factors (not disclosed by WuXi). The higher the enrichment score (higher than 1,000) and the higher the copy number (greater than 10), the higher the possibility of the selected compound being a potential binder. After analyzing the data, two compounds (**LAG-3 Hit I** and **LAG-3 Hit II**) were identified as the top binders. WuXi AppTec synthesized these two tag-free compounds, while the authors conducted their validation using various biophysical screening techniques.

### MST Screening for LAG-3 Binding

The MST assay for LAG-3 binding was performed using our previously reported protocol.^38^

### SPR Screening for LAG-3 Binding

The binding analysis was performed on a Biacore^™^ 8K instrument (Cytiva, Marlborough, MA, USA) at 25°C using 1x PBS-P+ buffer (Cat. # 28995084, Cytiva) supplemented with 2% DMSO as the running buffer. Biotinylated Human LAG-3 ECD Protein, His, Avitag^™^ (Cat. # LA3-H82E9, Acro Biosystems) was captured on a Series S Sensor Chip CAP at a flow rate of 10 μL/min for 60 seconds yielding an immobilization level of approximately 4,000 RU using the Biotin CAPture Kit (Cytiva). The solutions provided are a Biotin CAPture Reagent, which is a modified streptavidin, for capturing the ligand on to the sensor chip CAP and two regeneration stock solutions for the regeneration of the surface after each run. Single-cycle kinetics was used to determine the kinetic parameters of each small molecule. Serial dilutions of candidate small molecules, ranging from mid-micromolar to low-nanomolar concentrations, were prepared in running buffer and injected over the chip at a flow rate of 30 μL/min for 120 seconds (association phase). The dissociation phase was set at 10 minutes. A solvent correction step was included into the experimental method to adjust for variations in the bulk refractive index of the samples. Data was obtained using the Biacore 8K Control Software (Cytiva) and analyzed by non-linear curve fitting using a steady-state affinity analysis applying the software Biacore^™^ Insight Evaluation Software (Cytiva).

### CETSA for LAG-3 Engagement

This CETSA protocol was adapted from our previously published method for evaluating LAG-3 thermal stabilization by small molecules and optimized for use in Raji-hLAG-3 cells expressing full-length LAG-3.^38^

### Chemistry

Commercially available chemicals and solvents were used without prior purification. The synthesized compounds were analyzed using ^1^H and ^13^C NMR, as well as highresolution mass spectrometry (HRMS). ^1^H and ^13^C NMR spectra were acquired on a Bruker Avance III 500HD spectrometer (500 or 125 MHz, Billerica, MA, USA) using at room temperature. Chemical shifts are reported in parts per million (ppm) relative to tetramethylsilane (δ 0.00, s), with coupling constants given in hertz (Hz). Signal multiplicities are denoted as s (singlet), d (doublet), t (triplet), q (quartet), and m (multiplet). MS spectra were obtained using an SQ Detector 2 mass spectrometer (Waters, USA), while HRMS spectra were recorded on a Waters LCT Premier XE mass spectrometer (USA).

### General Procedure for the Synthesis of Compounds 1-11

To a stirred solution of compound **A** (1.0 equiv) in dry N,N-dimethylformamide (DMF) (0.1 M) was added HBTU (1.1 equiv) and triethylamine (3.0 equiv) at room temperature under a nitrogen atmosphere. The reaction mixture was stirred for 10 minutes to allow for activation of the carboxylic acid, after which the aromatic amine (compound **B**, 1.1 equiv) was added. The reaction mixture was stirred at room temperature for 12 hours, and progress was monitored by TLC. Subsequently, the reaction was quenched with water and extracted with ethyl acetate (3×). The combined organic layers were washed with brine, dried over anhydrous Na_2_SO_4_, filtered, and concentrated under reduced pressure. The crude product was purified by flash column chromatography (silica gel, gradient elution with ethyl acetate/hexanes) to afford the desired amide derivative.

#### Compound 1

Yield 82%. (3*R*,4*S*)-*N*-(3,4-dimethoxyphenyl)-4-phenyl-1-(pyridin-2-ylmethyl)pyrrolidine-3-carboxamide. ^1^H NMR (500 MHz, CDCl_3_) δ ppm: 2.75 (t, *J* = 8.6 Hz, 1H), 2.88 (t, *J* = 8.4 Hz, 1H), 2.97– 3.07 (m, 1H), 3.28 (dd, *J* = 9.6, 3.5 Hz, 1H), 3.48 (t, *J* = 8.8 Hz, 1H), 3.64 – 3.76 (m, 1H), 3.83 (d, *J* = 13.7 Hz, 1H), 3.86 (s, 3H), 3.88 (s, 3H), 4.08 (d, *J* = 13.6 Hz, 1H), 6.80 (d, *J* = 8.7 Hz, 1H), 6.95 (dd, *J* = 8.6, 2.5 Hz, 1H), 7.16 – 7.29 (m, 2H), 7.33 (d, *J* = 4.4 Hz, 4H), 7.41 (d, *J* = 7.8 Hz, 1H), 7.47 (d, *J* = 2.4 Hz, 1H), 7.70 (td, *J* = 7.6, 1.8 Hz, 1H), 8.59 (d, *J* = 5.2 Hz, 1H), 8.87 (s, 1H). ^13^C NMR (126 MHz, CDCl_3_) δ ppm: 48.64, 54.66, 55.89, 56.14, 57.32, 60.58, 61.78, 104.53, 111.32, 122.89, 122.41, 126.87, 127.43, 128.79, 132.42, 136.67, 143.37, 145.37, 148.96, 149.52, 158.20, 172.96. MS m/z: (ES+), [M + H]^+^, 418.30. HRMS (ESI): calculated for C_25_H_28_N_3_O_3_ (M + H)^+^, 418.2131; found, 418.2125.

#### Compound 2

Yield 73%. (3*R*,4*S*)-4-phenyl-1-(pyridin-2-ylmethyl)-*N*-(pyridin-4-yl)pyrrolidine-3-carboxamide. ^1^H NMR (500 MHz, CDCl_3_) δ ppm: 2.70 (t, *J* = 8.2 Hz, 1H), 2.72 – 2.79 (m, 1H), 3.00 – 3.11 (m, 1H), 3.34 (d, *J* = 9.9 Hz, 1H), 3.59 – 3.72 (m, 2H), 3.82 (d, *J* = 13.7 Hz, 1H), 4.18 (d, *J* = 13.7 Hz, 1H), 7.23 – 7.40 (m, 9H), 7.65 (d, *J* = 6.3 Hz, 2H), 7.73 (td, *J* = 7.6, 1.8 Hz, 1H), 8.51 (d, *J* = 6.4 Hz, 2H), 8.63 (d, *J* = 5.0 Hz, 1H), 9.98 (s, 1H). ^13^C NMR (126 MHz, CDCl_3_) δ ppm: 48.46, 54.65, 56.47, 59.76, 61.64, 113.73, 122.65, 122.85, 127.06, 127.35, 128.83, 136.75, 142.84, 145.96, 149.79, 150.50, 157.66, 174.46. MS m/z: (ES+), [M + H]^+^, 359.24. HRMS (ESI): calculated for C_22_H_23_N_4_O (M + H)^+^, 359.1882; found, 359.1872.

#### Compound 3

Yield 71%. (3*R*,4*S*)-*N*-benzyl-4-phenyl-1-(pyridin-2-ylmethyl)pyrrolidine-3-carboxamide. ^1^H NMR (500 MHz, CDCl_3_) δ ppm: 2.93 (t, *J* = 8.9 Hz, 1H), 3.07 (dq, *J* = 20.3, 7.4 Hz, 2H), 3.29 – 3.21 (m, 1H), 3.39 (t, *J* = 9.2 Hz, 1H), 3.66 (q, *J* = 7.9 Hz, 1H), 3.89 (d, *J* = 13.6 Hz, 1H), 4.07 (d, *J* = 13.7 Hz, 1H), 4.32 (dd, *J* = 15.0, 5.3 Hz, 1H), 4.48 (td, *J* = 15.9, 6.0 Hz, 1H), 6.59 (s, 1H), 7.65 (t, *J* = 7.7 Hz, 1H), 8.49 (s, 1H). ^13^C NMR (126 MHz, CDCl3) δ ppm: 43.40, 48.60, 53.62, 57.33, 60.70, 61.16, 122.56, 123.27, 127.00, 127.34, 127.52, 127.61, 128.62, 128.82, 136.85, 138.33, 142.38, 149.23, 157.16, 173.32. HRMS (ESI): calculated for C_24_H_26_N_3_O (M + H)^+^, 372.2076; found, 372.2072.

#### Compound 4

Yield 64%. (3*R*,4*S*)-*N*-(4-chlorobenzyl)-4-phenyl-1-(pyridin-2-ylmethyl)pyrrolidine-3-carboxamide. ^1^H NMR (500 MHz, CDCl_3_) δ ppm: 3.05 (t, *J* = 9.3 Hz, 1H), 3.14 (t, *J* = 6.9 Hz, 1H), 3.23 (s, 1H), 3.30 – 3.37 (m, 1H), 3.50 (t, *J* = 9.3 Hz, 1H), 3.67 (q, *J* = 8.5 Hz, 1H), 3.97 (d, *J* = 13.6 Hz, 1H), 4.16 (d, *J* = 13.6 Hz, 1H), 4.24 (dd, *J* = 15.0, 5.3 Hz, 1H), 4.48 (dd, *J* = 15.0, 6.6 Hz, 1H), 6.63 (t, *J* = 5.5 Hz, 1H), 7.06 (d, *J* = 8.4 Hz, 2H), 7.21 – 7.35 (m, 11H), 7.42 (d, *J* = 7.9 Hz, 1H), 7.57 – 7.77 (m, 2H), 8.50 (d, *J* = 5.5 Hz, 1H). ^13^C NMR (126 MHz, CDCl_3_) δ ppm: 42.66, 48.58, 53.31, 57.28, 60.53, 60.99, 122.88, 123.45, 127.31, 127.55, 127.88, 128.72, 128.89, 128.94, 133.11, 136.82, 136.94, 137.40, 149.48. MS m/z: (ES+), [M + H]^+^, 406.21, [M + 2H]^+^, 407.44. HRMS (ESI): calculated for C_24_H_25_N_3_OCl (M + H)^+^, 406.1686; found, 406.1683.

#### Compound 5

Yield 81%. (3*R*,4*S*)-*N*-phenethyl-4-phenyl-1-(pyridin-2-ylmethyl)pyrrolidine-3-carboxamide. ^1^H NMR (500 MHz, CDCl_3_) δ ppm: 2.73 (t, *J* = 8.5 Hz, 1H), 2.79 (t, *J* = 6.9 Hz, 2H), 2.90 (dd, *J* = 13.4, 6.6 Hz, 3H), 3.10 (d, *J* = 4.4 Hz, 1H), 3.25 (t, *J* = 8.9 Hz, 1H), 3.39 – 3.62 (m, 4H), 3.80 (d, *J* = 13.7 Hz, 1H), 3.95 (d, *J* = 13.7 Hz, 1H), 6.47 (t, *J* = 6.0 Hz, 1H), 7.03 (d, *J* = 6.1 Hz, 2H), 7.12 – 7.37 (m, 8H), 7.67 (td, *J* = 7.7, 1.9 Hz, 1H), 8.43 (d, *J* = 6.1 Hz, 2H), 8.54 (d, *J* = 6.7 Hz, 1H). ^13^C NMR (CDCl_3_, d ppm): 35.04, 39.45, 48.66, 53.62, 57.60, 61.04, 61.83, 122.28, 122.87, 124.12, 126.85, 127.37, 128.76, 136.62, 143.22, 148.06, 149.30, 149.83, 158.27, 174.32. MS m/z: (ES+), [M + H]^+^, 387.25. HRMS (ESI): calculated for C_25_H_27_N_3_O (M + 2H)^+2^, 387.2222; found, 387.2224.

#### Compound 6

Yield 59%. (3*R*,4*S*)-*N*-(2-(5-methoxy-1H-in-dol-2-yl)ethyl)-4-phenyl-1-(pyridin-2-ylmethyl)pyrrolidine-3-carboxamide. ^1^H NMR (500 MHz, CDCl_3_) δ ppm: 3.01 – 2.81 (m, 4H), 3.11 – 3.25 (m, 2H), 3.35 (t, *J* = 9.2 Hz, 1H), 3.45 – 3.67 (m, 3H), 3.86 (s, 3H), 3.93 (d, *J* = 13.4 Hz, 1H), 4.04 (d, *J* = 13.4 Hz, 1H), 6.06 (t, *J* = 6.0 Hz, 1H), 6.80 (s, 1H), 6.87 (dd, *J* = 8.9, 2.4 Hz, 1H), 7.01 (d, *J* = 2.6 Hz, 1H), 7.20 – 7.30 (m, 8H), 7.39 (d, *J* = 7.8 Hz, 1H), 7.66 (td, *J* = 7.6, 1.8 Hz, 1H), 7.97 (s, 1H), 8.57 (d, *J* = 3.4 Hz, 1H). ^13^C NMR (126 MHz, CDCl_3_) δ ppm: 25.22, 39.22, 48.34, 53.41, 55.95, 57.36, 60.59, 60.83, 100.53, 111.96, 112.37, 112.50, 122.77, 122.83, 123.58, 127.03, 127.52, 127.66, 128.82, 131.51, 136.96, 141.92, 149.22, 154.05, 156.44, 172.73. HRMS (ESI): calculated for C_28_H_30_N_4_O_3_ (M + H)^+^, 455.2447; found, 455.2432.

#### Compound 7

Yield 76%. (3*R*,4*S*)-4-phenyl-1-(pyridin-2-ylmethyl)-*N*-(2-(2-(p-tolyl)oxazol-4-yl)ethyl)pyrrolidne-3-carboxamide. ^1^H NMR (500 MHz, CDCl_3_) δ 2.41 (s, 3H), 2.73 (t, *J* = 6.5 Hz, 2H), 2.86 – 2.79 (m, 1H), 3.01 – 2.97 (m, 2H), 3.05 (t, *J* = 8.7 Hz, 1H), 3.21 (dd, *J* = 9.6, 5.6 Hz, 1H), 3.29 (t, *J* = 8.9 Hz, 1H), 3.47 – 3.83 (m, 3H), 3.88 (d, *J* = 13.7 Hz, 1H), 4.00 (d, *J* = 13.7 Hz, 1H), 6.61 (t, *J* = 5.8 Hz, 1H), 7.20 – 7.12 (m, 2H), 7.13 – 7.31 (m, 2H), 7.36 (s, 1H), 7.41 (d, *J* = 7.8 Hz, 1H), 7.63 (td, *J* = 7.7, 1.8 Hz, 1H) 7.84 (d, *J* = 8.2 Hz, 2H), 8.53 (d, *J* = 2.6 Hz, 1H). ^13^C NMR (126 MHz, CDCl_3_) δ 21.53, 26.05, 38.25, 48.56, 53.50, 57.69, 61.10, 61.71, 122.33, 122.94, 124.71, 126.25, 126.88, 127.39, 128.78, 129.48, 134.47, 136.70, 139.27, 140.65, 142.72, 149.22, 157.76, 161.89, 173.74. HRMS (ESI): calculated for C_29_H_31_N_4_O_2_ (M + H)^+^, 467.2447; found, 467.2457.

#### Compound 8

Yield 88%. (3*R*,4*S*)-N-(4-(2-methylthiazol-4-yl)phenethyl)-4-phenyl-1-(pyridin-2-ylmethyl)pyrrolidine-3-carboxamide. ^1^H NMR (500 MHz, CDCl_3_) δ ppm: 2.98 – 2.73 (m, 9H), 3.14 (dd, *J* = 9.5, 5.2 Hz, 1H), 3.27 (t, *J* = 8.9 Hz, 1H), 3.62 – 3.41 (m, 4H), 3.84 (d, *J* = 13.6 Hz, 1H), 3.95 (d, *J* = 13.7 Hz, 1H), 6.16 (t, *J* = 6.0 Hz, 1H), 7.31 – 7.14 (m, 10H), 7.36 (d, *J* = 7.8 Hz, 1H), 7.65 (td, *J* = 7.7, 1.9 Hz, 1H), 7.80 – 7.74 (m, 2H), 8.54 (d, *J* = 4.9 Hz, 1H). ^13^C NMR (126 MHz, CDCl_3_) δ ppm: 19.35, 35.44, 40.34, 48.62, 53.69, 57.71, 61.11, 61.72, 111.91, 122.25, 122.91, 126.52, 126.81, 127.43, 128.75, 129.10, 132.93, 136.64, 138.74, 143.16, 149.24, 154.91, 158.12, 165.82, 173.99. HRMS (ESI): calculated for C_29_H_30_N_4_OSNa (M + Na)^+^, 505.2038; found, 505.2060.

#### Compound 9

Yield 53%. (3*R*,4*S*)-*N*-(2-([1,1’-biphenyl]-4-yl)ethyl)-4-phenyl-1-(pyridin-2-ylmethyl)pyrrolidine-3-carboxamide. ^1^H NMR (500 MHz, Acetone) δ ppm: 2.69 – 2.92 (m, 3H), 3.32 – 3.58 (m, 4H), 3.82 – 4.26 (m, 5H), 4.91– 5.03 (m, 2H), 7.16 (d, *J* = 8.2 Hz, 2H), 7.32 – 7.56 (m, 13H), 7.61 (dd, *J* = 8.3, 1.3 Hz, 2H), 7.77 (d, *J* = 7.8 Hz, 1H), 8.01 (td, *J* = 7.7, 1.8 Hz, 1H), 8.70 (d, *J* = 5.6 Hz, 1H). HRMS (ESI): calculated for C_31_H_32_N_3_O (M + H)^+^, 462.2530; found, 462.2545.

#### Compound 10

Yield 71%. (3*R*,4*S*)-4-phenyl-1-(pyridin-2-ylmethyl)-*N*-(4-(pyridine-4-yl)benzyl)pyrrolidine-3-carboxamide. ^1^H NMR (500 MHz, CDCl_3_) δ ppm: 2.83 – 2.76 (m, 1H), 3.03 – 2.88 (m, 2H), 3.21 (dd, *J* = 9.2, 4.4 Hz, 1H), 3.35 (t, *J* = 8.9 Hz, 1H), 3.64 (td, *J* = 7.9, 6.0 Hz, 1H), 3.80 (d, *J* = 13.7 Hz, 1H), 4.01 (d, *J* = 13.7 Hz, 1H), 4.41 (dd, *J* = 15.0, 5.4 Hz, 1H), 4.58 (dd, *J* = 15.0, 6.4 Hz, 1H), 6.81 (t, *J* = 6.0 Hz, 1H), 7.14 (ddd, *J* = 7.6, 4.9, 1.3 Hz, 1H), 7.28 – 7.21 (m, 2H), 7.53 – 7.46 (m, 3H), 7.65 – 7.57 (m, 3H), 8.49 – 8.43 (m, 1H), 8.70 – 8.63 (m, 3H). ^13^C NMR (126 MHz, CDCl^3^) δ ppm: 42.92, 48.78, 53.82, 57.51, 60.99, 61.86, 121.50, 122.18, 122.83, 126.85, 127.21, 127.47, 128.33, 128.77, 136.56, 137.11, 139.79, 143.35, 147.89, 149.31, 150.31, 158.36, 174.35. HRMS (ESI): calculated for C_29_H_28_N_4_ONa (M + Na)^+^, 471.2161; found, 471.2177.

#### Compound 11

Yield 80%. (3*R*,4*S*)-*N*-([1,1’-biphenyl]-4-ylmethyl)-4-phenyl-1-(pyridin-2-ylmethyl)pyrrolidine-3-carboxamide. ^1^H NMR (500 MHz, CDCl_3_) δ ppm: 2.91 – 2.81 (m, 2H), 3.09 – 3.00 (m, 2H), 3.31 – 3.24 (m, 1H), 3.40 (t, *J* = 8.9 Hz, 1H), 3.67 (q, *J* = 8.2 Hz, 1H), 3.89 (d, *J* = 13.7 Hz, 1H), 4.06 (d, *J* = 13.9 Hz, 1H), 4.36 (dd, *J* = 14.9, 5.3 Hz, 1H), 4.55 (dd, *J* = 14.9, 6.3 Hz, 1H), 6.68 (s, 1H), 7.20 – 7.12 (m, 1H), 7.25 (dd, *J* = 8.6, 4.2 Hz, 3H), 7.32 (d, *J* = 4.4 Hz, 4H), 7.40 – 7.33 (m, 2H), 7.46 (t, *J* = 7.6 Hz, 2H), 7.53 (d, *J* = 8.2 Hz, 2H), 7.64 – 7.57 (m, 3H), 8.46 (d, *J* = 6.3 Hz, 1H). ^13^C NMR (126 MHz, CDCl_3_) δ ppm: 43.10, 48.75, 53.53, 57.52, 60.86, 61.73, 122.40, 122.93, 127.00, 127.05, 127.36, 127.50, 128.05, 128.83, 136.73, 137.43, 140.32, 140.74, 142.57, 149.31, 157.54, 173.92. HRMS (ESI): calculated for C_30_H_30_N_3_O (M + H)^+^, 448.2389; found, 448.2391.

### HPLC Purity

The purity of **LAG-3 Hit II** was confirmed to be >95% using both LCMS (Figure S7) and HPLC (Figure 9). The purity of compound **11** was confirmed to be >95% using HPLC analysis (Figure S64) with a reversed-phase column Phenomenex Gemini, C18 (250 mm × 4.60 mm, 5 μm) on an HPLC Agilent system. The mobile phase used was an acetonitrile-H_2_O gradient and a 1 mL/min flow rate. UV absorption at 210 and 254 nm was used to monitor the method.

### LAG-3/MHCII Blockade Cellular Assay

The cellular LAG-3/MHCII blockade assay was performed using the LAG-3/MHCII blockade kit from Promega (Catalog # JA1111). The assay was performed according to the manufacturer’s recommended protocol by co-culturing both MHCII-positive human cell line and NanoLuc^®^ (NL) luciferase reporter/Jurkat LAG-3 cells in the presence of increasing concentrations of compound **11** (n=3).

### PBMC/Cancer Cells Coculture Assay

To evaluate the efficacy of LAG-3 inhibitors, PBMCs from healthy donors were cocultured with THP-1, Kausumi-1, or A549 cells at a 1:2 ratio. The cells were treated with compound **11** (10 μM, n=3) or relatlimab (100 μg/ml, n=3). IFN-γ levels were measured using an IFN-γ ELISA kit (from R&D Systems). The killing activity of the treated PBMCs was assessed at 24 h using a CFSE/7-ADD assay kit (from Cell Signaling Technology) to measure AML cell viability. Results were compared to a control group treated with anti-LAG-3 antibody (relatlimab; 100 µg/ml) for comparison.

### Computational Study

Comparative analysis of human and mouse proteins was carried out using the in-house script, which aligned the amino acid sequence of human LAG-3 with the mouse LAG-3 sequence for similarities and patterns. The threedimensional structure of the human LAG-3/HLA-DRA complex was generated by docking the extracellular region of LAG-3 (PDB ID: 7TZG) with HLA-DRA PDB ID: 4MD4). Among the 30 different possible conformations (data not shown), only the chosen protein-protein complex exhibited a complex similar in shape and binding interface location as that of the reported mouse LAG3/MHC II complex. The generated model was subjected to energy minimization and structural refinement using Maestro Schrödinger Suite with the OPLS3e force field.

The predicted LAG-3/HLA-DRA complex was analyzed to identify contact points between the two proteins. The binding orientation of Arg125 in human LAG-3 was compared to that observed in the mouse complex to assess the reliability of the model. Putative hot spot residues within the LAG-3 interface were identified based on their spatial proximity to HLA-DRA and their physicochemical properties (e.g., charged polar residues).

Next, compound **11** was docked into the identified hot spot residues of LAG-3 at the LAG-3/HLA-DRA interface using the Induced Fit Docking (IFD) protocol implemented in Maestro Schrödinger Suite (v2021.2). This approach was selected to account for the flexibility of both the ligand and the protein binding site, allowing for conformational adjustments during the docking process. The ligand structure was prepared using LigPrep module within Maestro. The protein structure was prepared using the protein preparation wizard within Maestro. The IFD protocol involved an initial Glide docking step, followed by Prime refinement of the protein-ligand complex, and a final Glide redocking step. The resulting docked poses were ranked based on their Glide score and analyzed for their interaction patterns. The docked poses of compound **11** within the LAG-3 binding site were analyzed and visualized using the Discovery Studio Visualizer software.

The molecular dynamics investigation was carried out according to the methodology of our previously published work.^51^

### PK Studies

The preliminary evaluation of PK parameters for compound **11** was performed as previously reported by us.^38^

## ASSOCIATED CONTENT

### Supporting Information

The Supporting Information is available free of charge on the ACS Publications website.

NMR spectra, mass spectrometry charts, HPLC traces, binding

curves, and computational studies (PDF)

Molecular formula strings (CSV)

LAG-3 PPI homolog (PDB)

LAG-3/compound **11** complex (PDB)

## AUTHOR INFORMATION

### Author Contributions

The manuscript was written through contributions of all authors. All authors have given approval to the final version of the manuscript.

## ABBREVIATIONS

7-AAD: 7-Aminoactinomycin D
ADME: Absorption, Distribution, Metabolism, Excretion
AML: Acute Myeloid Leukemia
APCs: Antigen-Presenting Cells
CETSA: Cellular Thermal Shift Assay
CFSE: Carboxyfluorescein Succinimidyl Ester
CRBN: Cereblon
CTLA-4: Cytotoxic T-Lymphocyte-Associated Protein 4
DEL: DNA-Encoded Library
ECD: Extracellular domain
FaS-SIF: fasted state simulated intestinal fluid
FGL1: Fibrinogen-like Protein 1
hERG: Human Ether-a-go-go-Related Gene
HPLC: High-Performance Liquid Chromatography
HRMS: High-Resolution Mass Spectrometry
HTS: High-Throughput Screening
ICB: Immune Checkpoint Blockade
IFD: Induced Fit Docking
IFN-γ: Interferon Gamma
irAEs: Immune-Related Adverse Events
LAG-3: Lymphocyte Activation Gene 3 Protein
LCMS: Liquid Chromatography-Mass Spectrometry
mAbs: Monoclonal Antibodies
MD: Molecular Dynamics
MHCII: Major Histocompatibility Complex Class II
MST: Microscale Thermophoresis
NMR: Nuclear Magnetic Resonance
NSCLC: Non-Small Cell Lung Cancer
PAINS: Pan-Assay Interference Compounds
PBMCs: Peripheral Blood Mononuclear Cells
PD-1: Programmed Cell Death Protein 1
PK: Pharmacokinetic
PPB: Plasma Protein Binding
Rg: Radius of Gyration
RMSD: Root Mean Square Deviation
RMSF: Root Mean Square Fluctuation
SAR: Structure-Activity Relationship
SFC: Supercritical Fluid Chromatography
SPR: Surface Plasmon Resonance
TCR: T-cell Receptor
TILs: Tumor-Infiltrating Lymphocytes
TIM-3: T-cell Immunoglobulin Mucin-3
VISTA: V-domain Immunoglobulin Suppressor of T-cell Activation

## Notes

The authors declare no competing financial interests.

## ACKNOWLEDGMENTS

We gratefully acknowledge financial support from the ELSA U. Pardee Foundation (Award ID: 2022-215).

## Notes

### Competing Interest Statement

The authors have declared no competing interest.

