## Supporting Information for "Discovery and Optimization of LAG-3-Targeted Small Molecules via DNA-Encoded Chemical Library (DEL) Screening for Cancer Immunotherapy"

*Electronic Supplementary Information*

| **Contents** |  |
| --- | --- |
| ^1^H NMR spectrum of **LAG-3** **Hit I** | S3 |
| LCMS report of **LAG-3** **Hit I** | S4 |
| Mass spectrum of **LAG-3** **Hit I** | S5 |
| HPLC purity of **LAG-3** **Hit I** | S6 |
| Chiral SFC report of **LAG-3** **Hit I** | S7 |
| ^1^H NMR spectrum of **LAG-3** **Hit II** | S8 |
| LCMS report of **LAG-3** **Hit II** | S9 |
| Mass spectrum of **LAG-3** **Hit II** | S10 |
| HPLC purity of **LAG-3** **Hit II** | S11 |
| Chiral SFC report of **LAG-3** **Hit II** | S13 |
| MST binding of **LAG-3** **Hit II** (increasing concentrations, n=4) to LAG-3 | S14 |
| MST binding of **LAG-3** **Hit II** (increasing concentrations, n=4) to LAG-3 | S14 |
| Three independent dose-response curves for the binding interaction between **LAG-3 Hit II** and LAG-3 (top). The corresponding MST time traces are shown on the bottom. | S15 |
| Cellular thermal shift assay (CETSA) for the binding of **LAG-3 Hit II** (25 μM) to LAG-3 in cell lysate of LAG-3 expressing Raji cells. Error bars represent standard deviation (n=5). | S16 |
| Spectral data for compounds **1-11** | S17 |
| MST binding curves for compounds **1-11** | S48 |
| **A.** Production of IFNγ from PBMCs upon co-culturing with Kasumi-1 cells in the absence and presence of relatilmab (100 µg/ml) and compound **11** (10 μM), *** *p* < 0.001 in comparison to control. **B.** The % of dead Kasumi-1 cells as assessed by 7-AAD/CFSE assay in the co-culture assay of PBMCs and Kasumi-1 in the absence and presence of relatilmab (100 µg/ml) and compound **11** (10 μM), * *p* < 0.05, and *** *p* < 0.001. Error bars represent standard deviation (n = 3). | S52 |
| **A.** Production of IFNγ from PBMCs upon co-culturing with A549 cells in the absence and presence of relatilmab (100 µg/ml) and compound **11** (10 μM), *** *p* < 0.001 in comparison to control. **B.** The % of dead A549 cells as assessed by 7-AAD/CFSE assay in the co-culture assay of PBMCs and A549 in the absence and presence of relatilmab (100 µg/ml) and compound **11** (10 μM), * *p* < 0.05, and *** *p* < 0.001. Error bars represent standard deviation (n = 3). | S53 |
| Sequence conservation diagram comparing mouse and human LAG-3 | S54 |
| Sequence similarity diagram comparing mouse and human LAG-3 | S55 |
| Heatmap of amino acids at each position obtained by aligning mouse and human LAG-3 sequences | S56 |
| Hydrophobicity profile comparing mouse and human LAG-3 sequences | S57 |
| RMSD calculations for the unbound LAG-3 and LAG-3/ compound **11** complex | S58 |
| RMSF calculations for the unbound LAG-3 and LAG-3/ compound **11** complex | S59 |
| Gibbs free energy diagram of LAG-3/ compound **11** complex | S59 |
| HPLC trace of compound **11** | S60 |

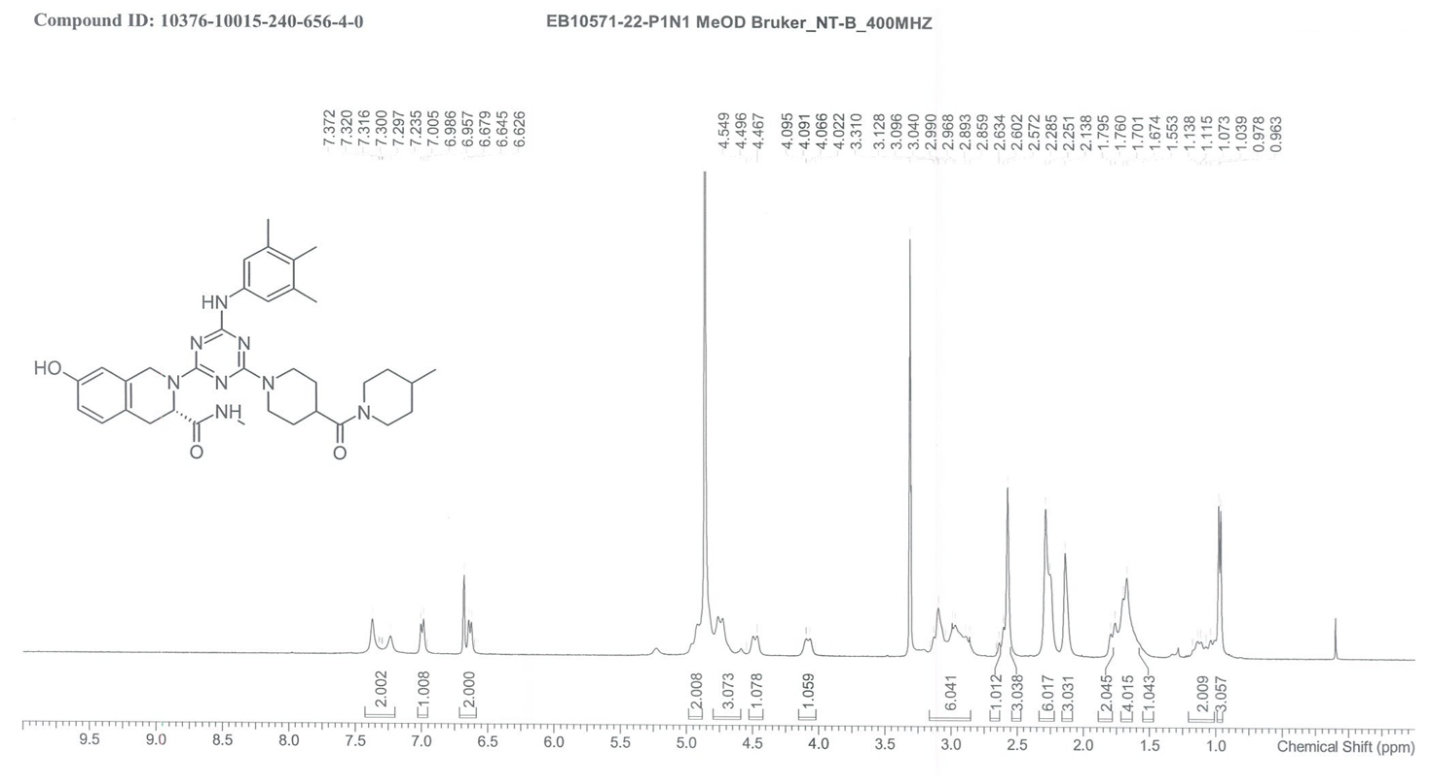

**Figure S1**. ^1^H NMR spectrum of **LAG-3** **Hit I**.

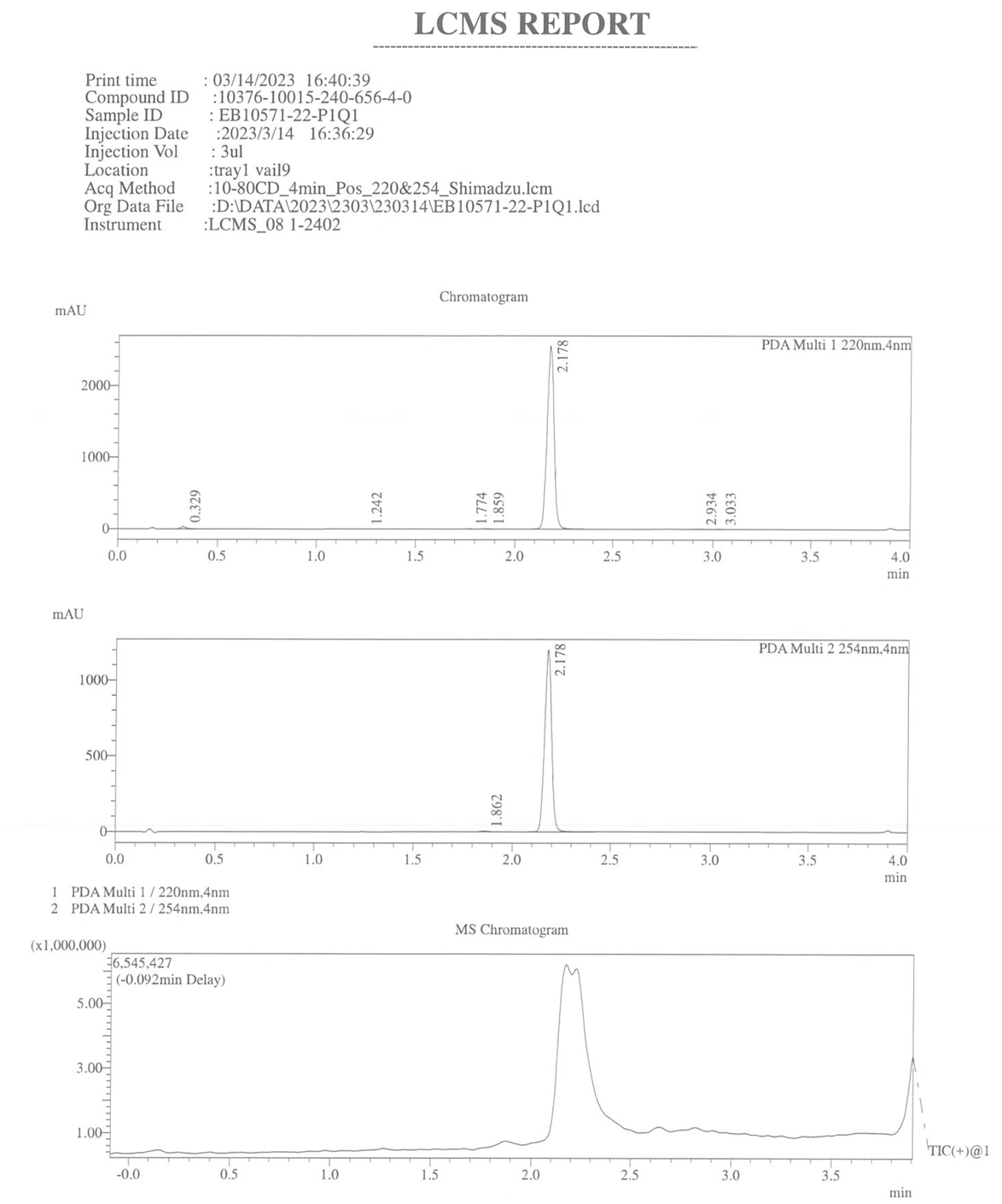

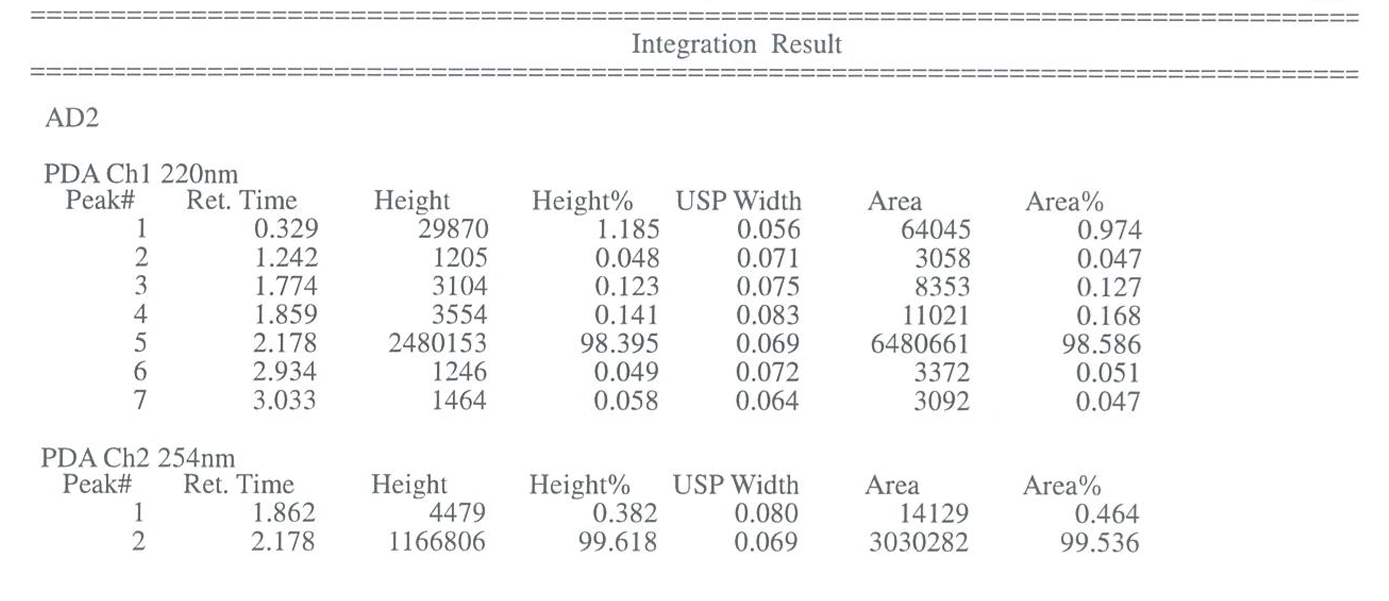

**Figure S2**. LCMS report of **LAG-3** **Hit I**.

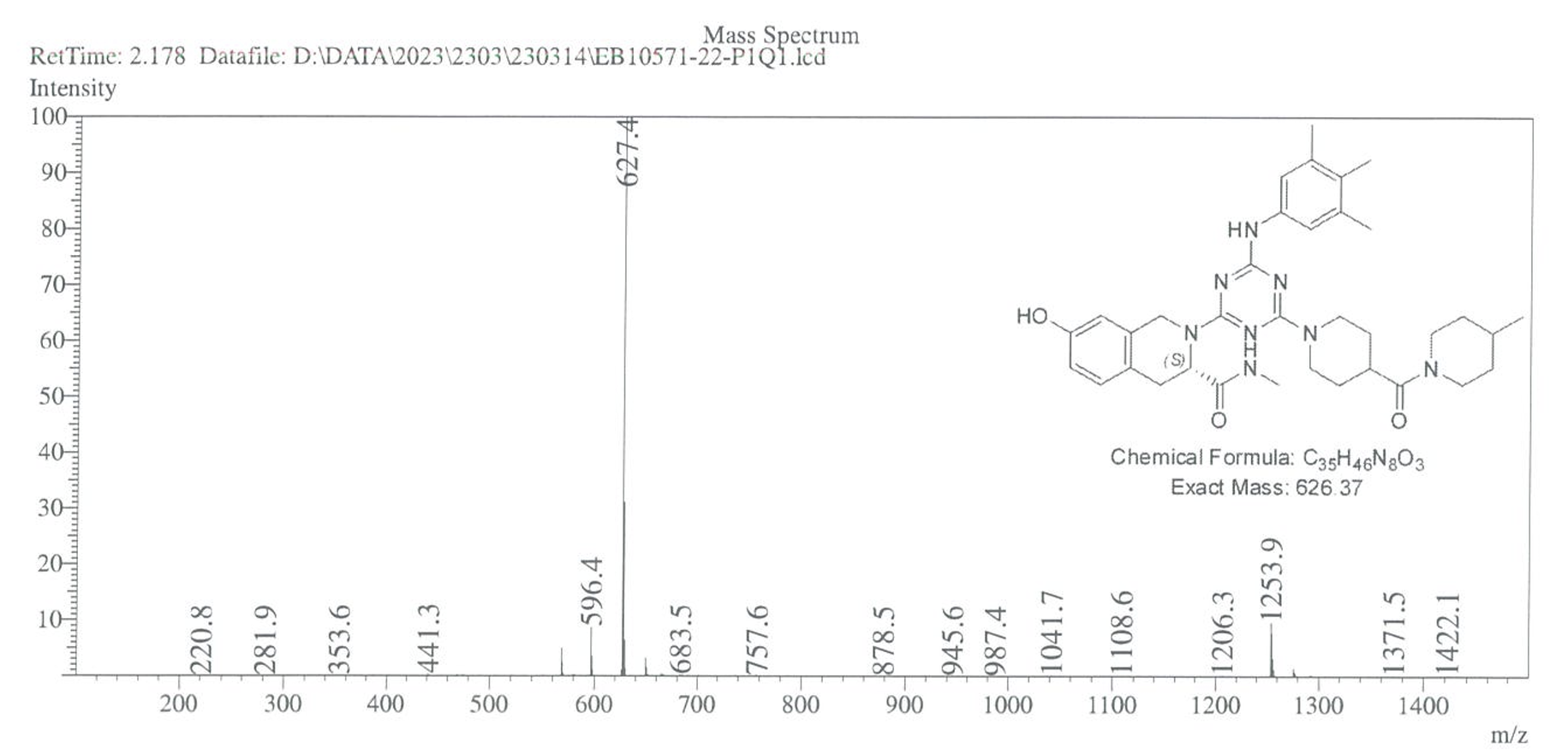

**Figure S3**. Mass spectrum of **LAG-3** **Hit I**.

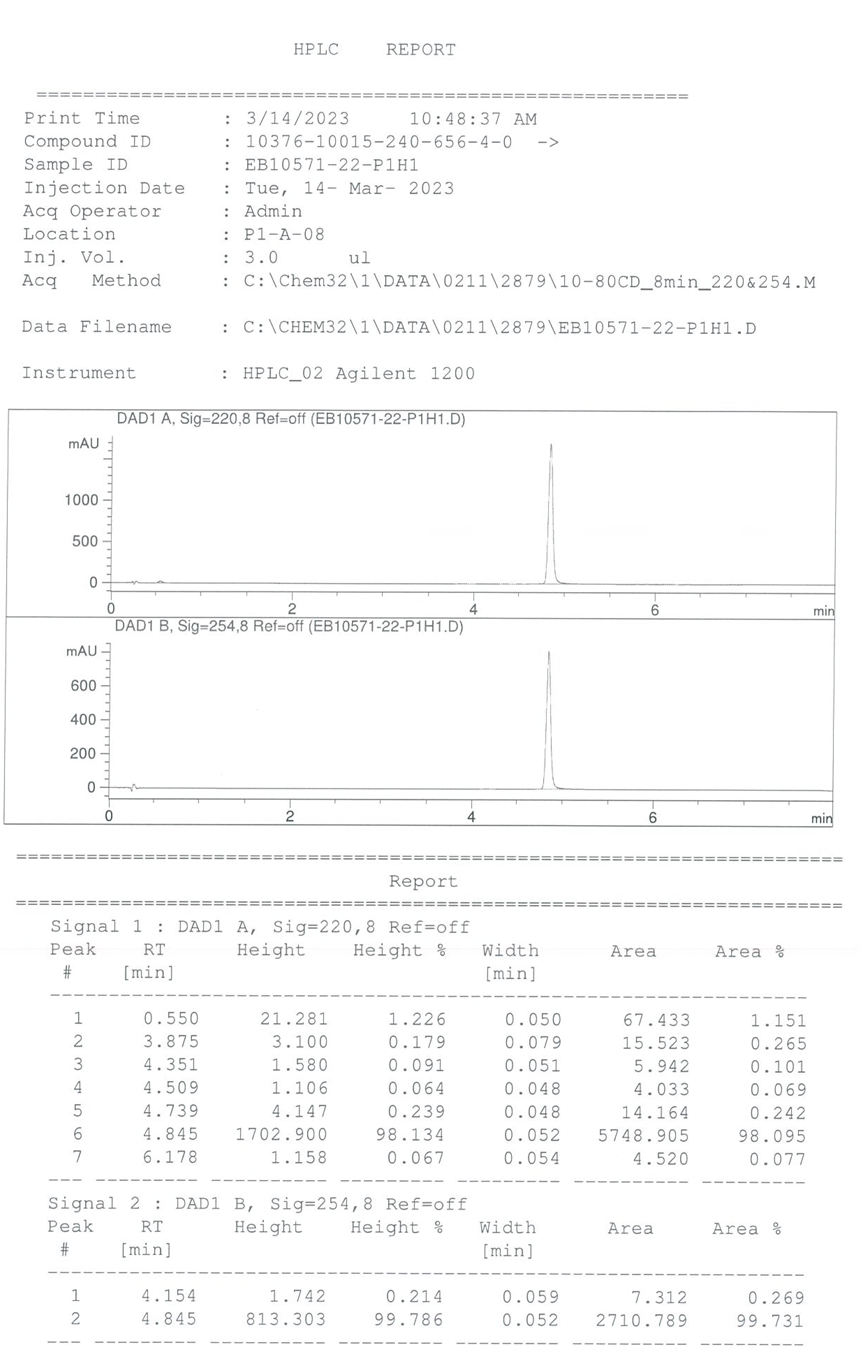

**Figure S4**. HPLC purity of **LAG-3** **Hit I**.

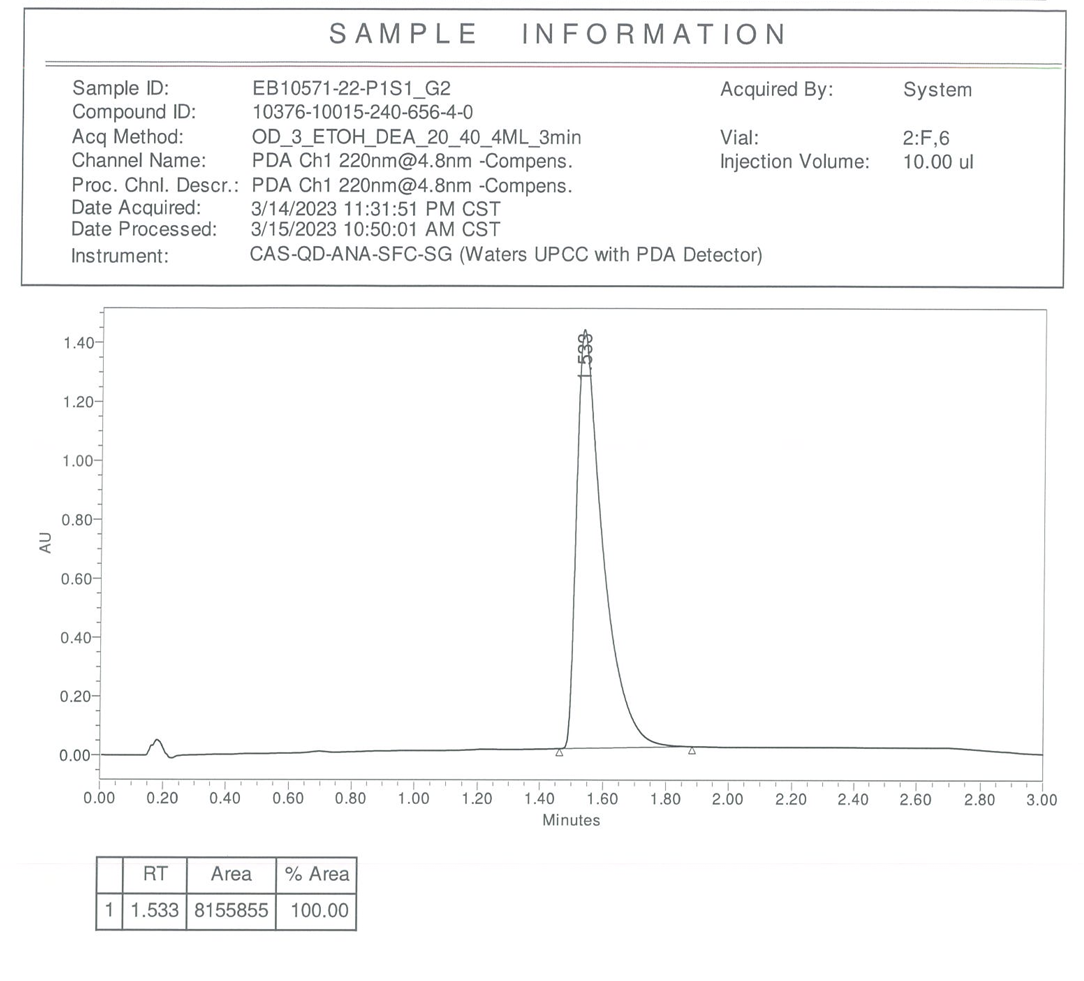

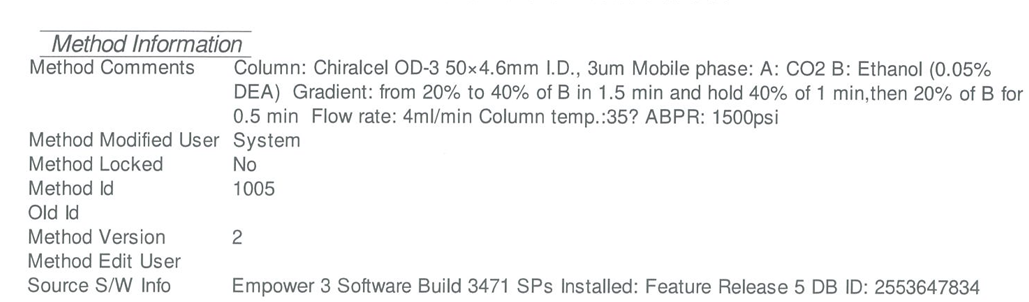

**Figure S5**. Chiral SFC report of **LAG-3** **Hit I**.

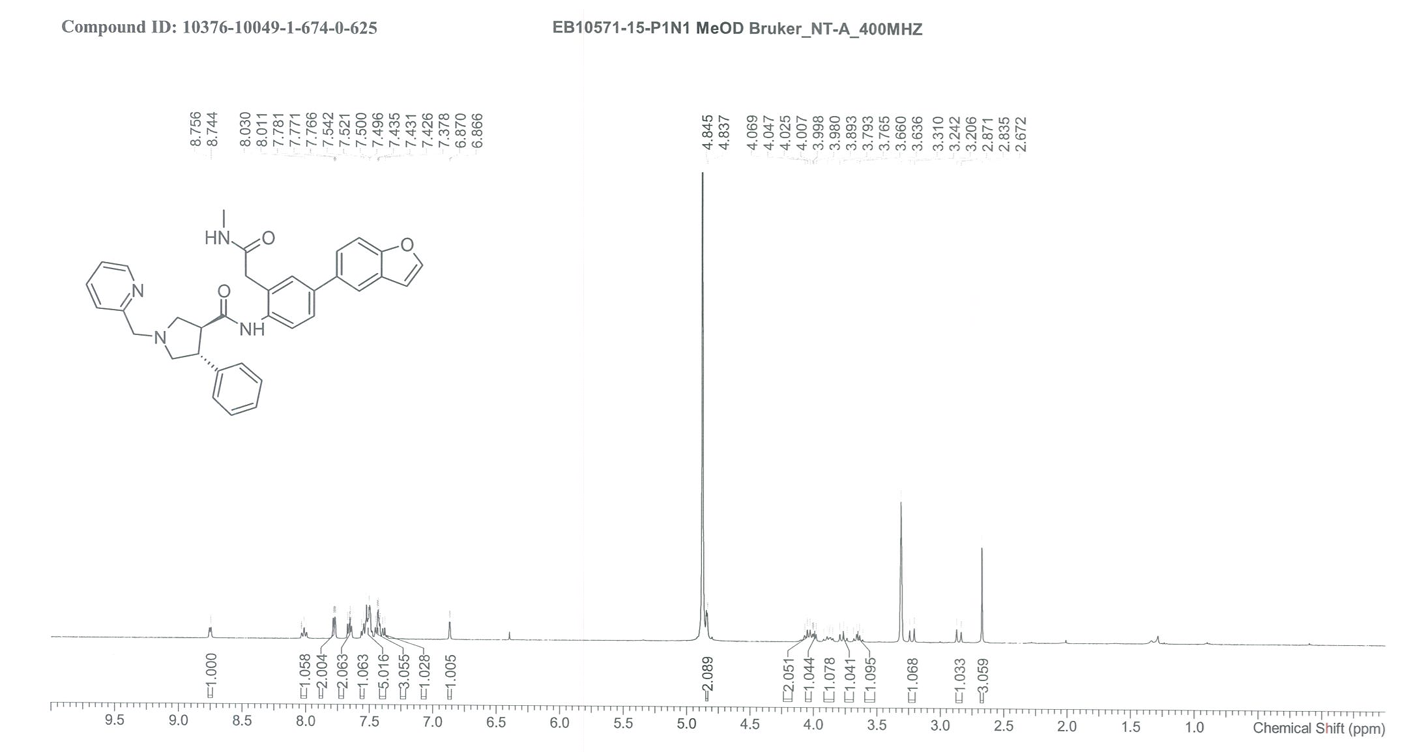

**Figure S6**. ^1^H NMR spectrum of **LAG-3** **Hit II**.

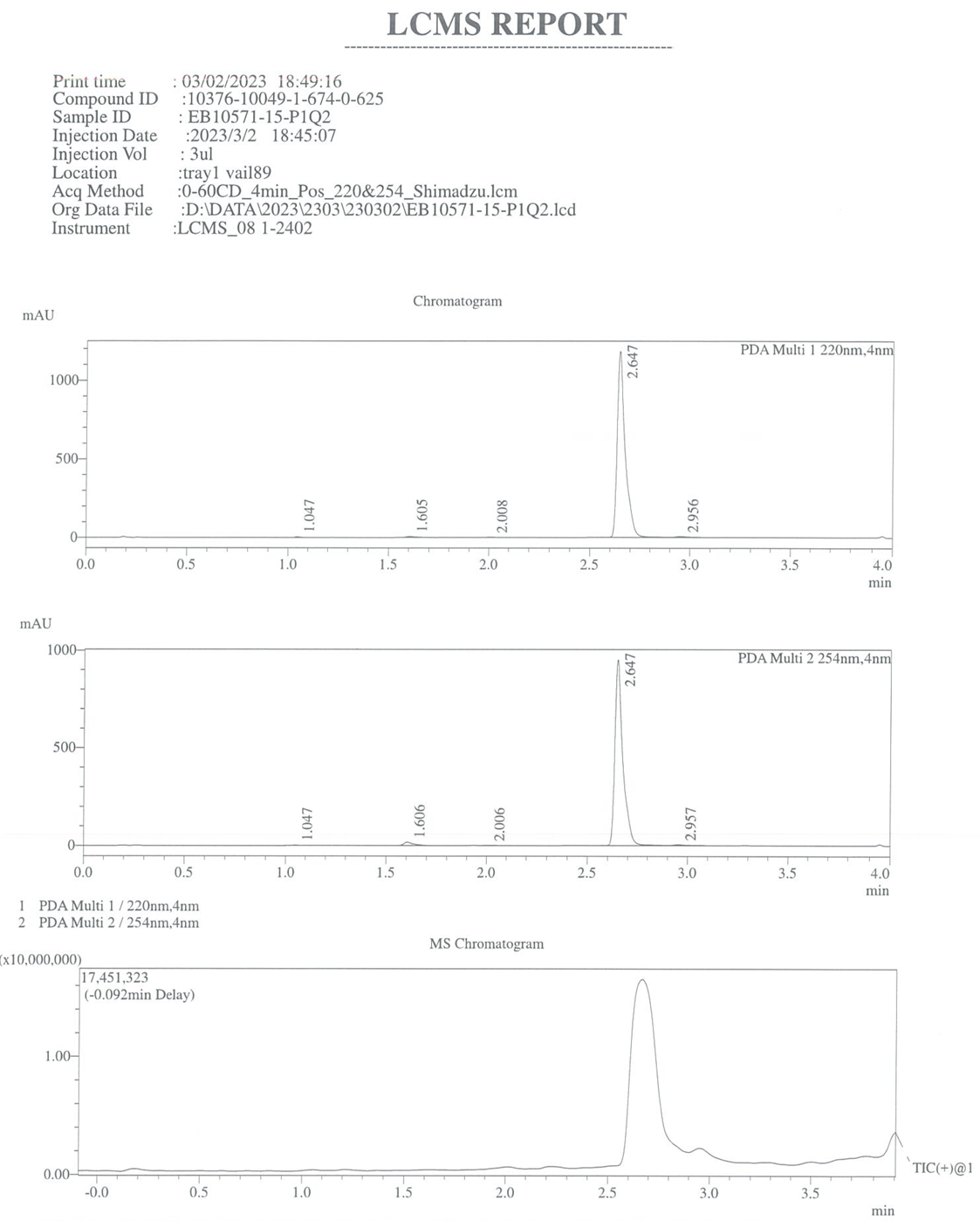

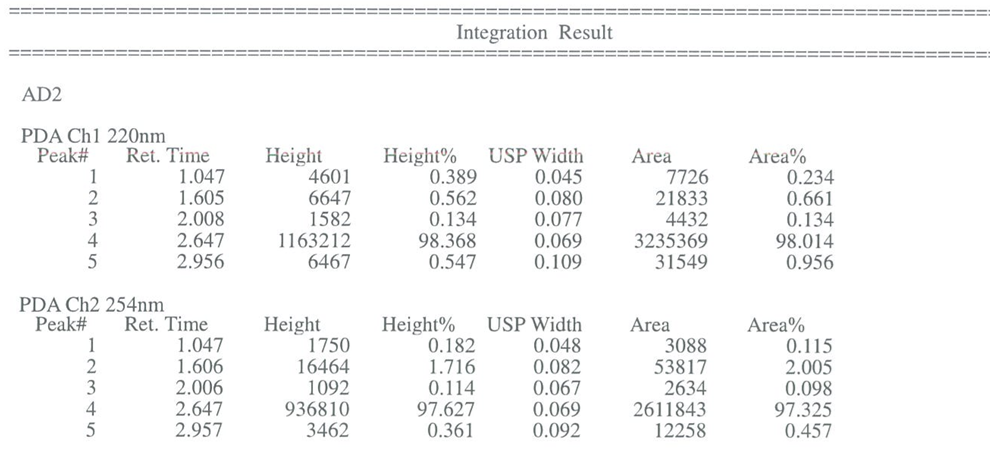

**Figure S7**. LCMS report of compound **LAG-3** **Hit II**.

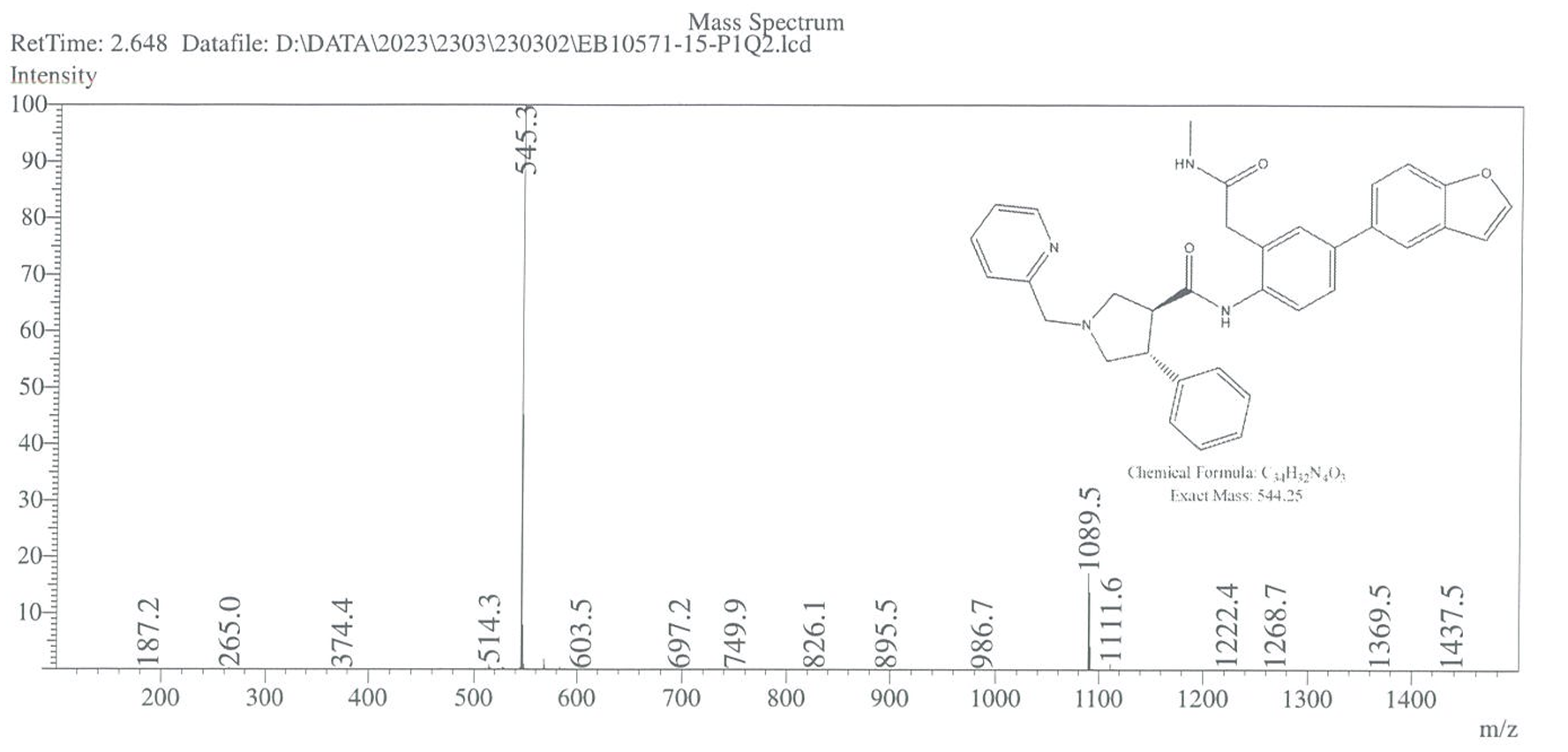

**Figure S8**. Mass spectrum of **LAG-3** **Hit II**.

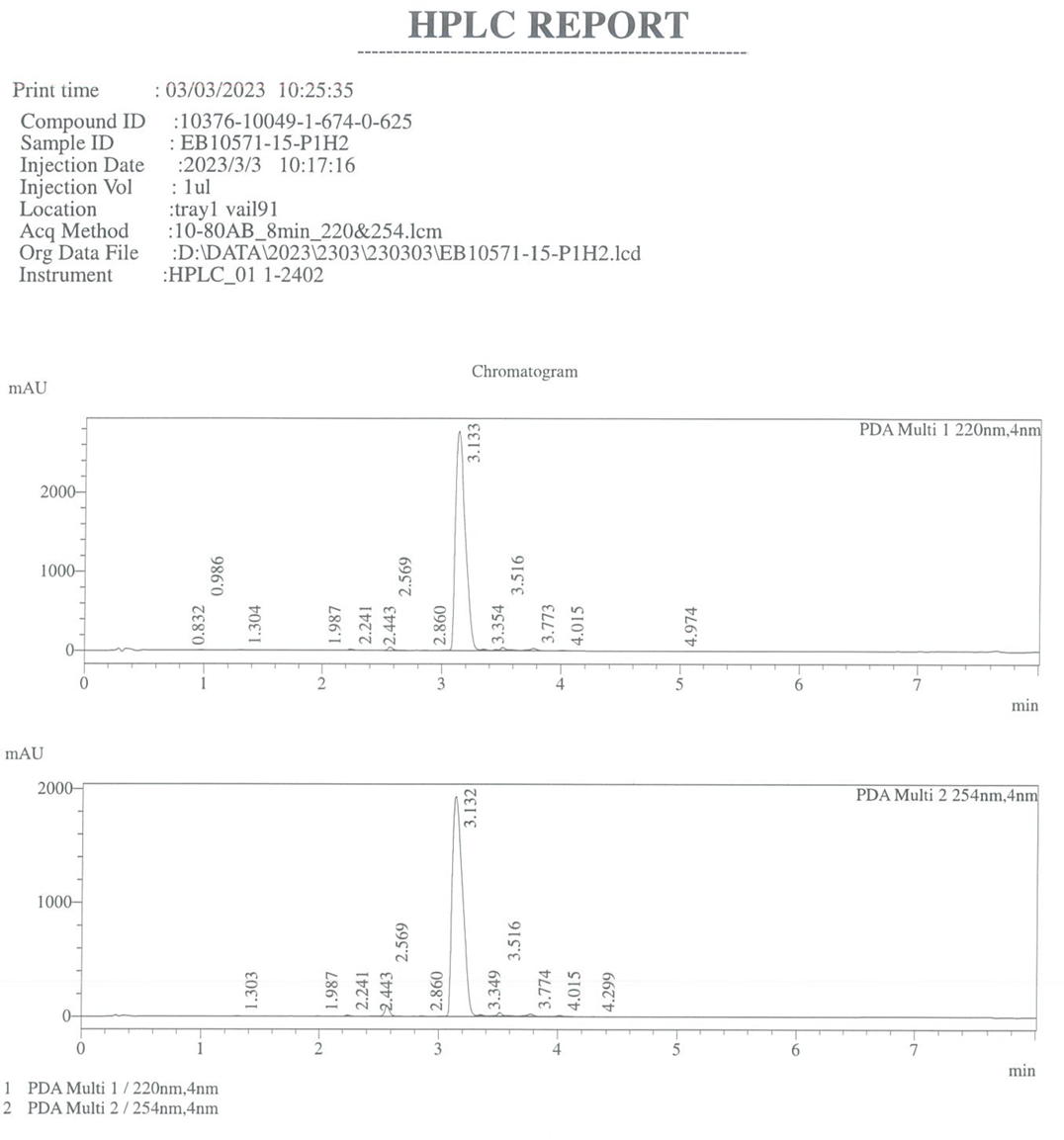

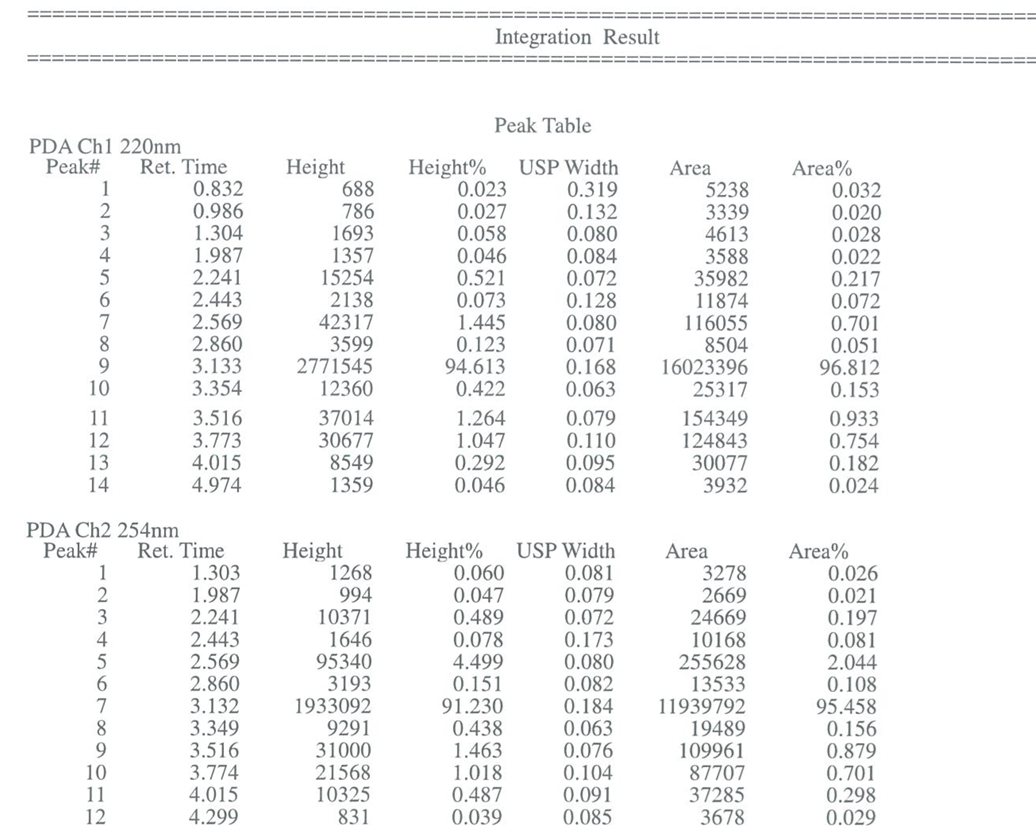

**Figure S9**. HPLC purity of **LAG-3** **Hit II**.

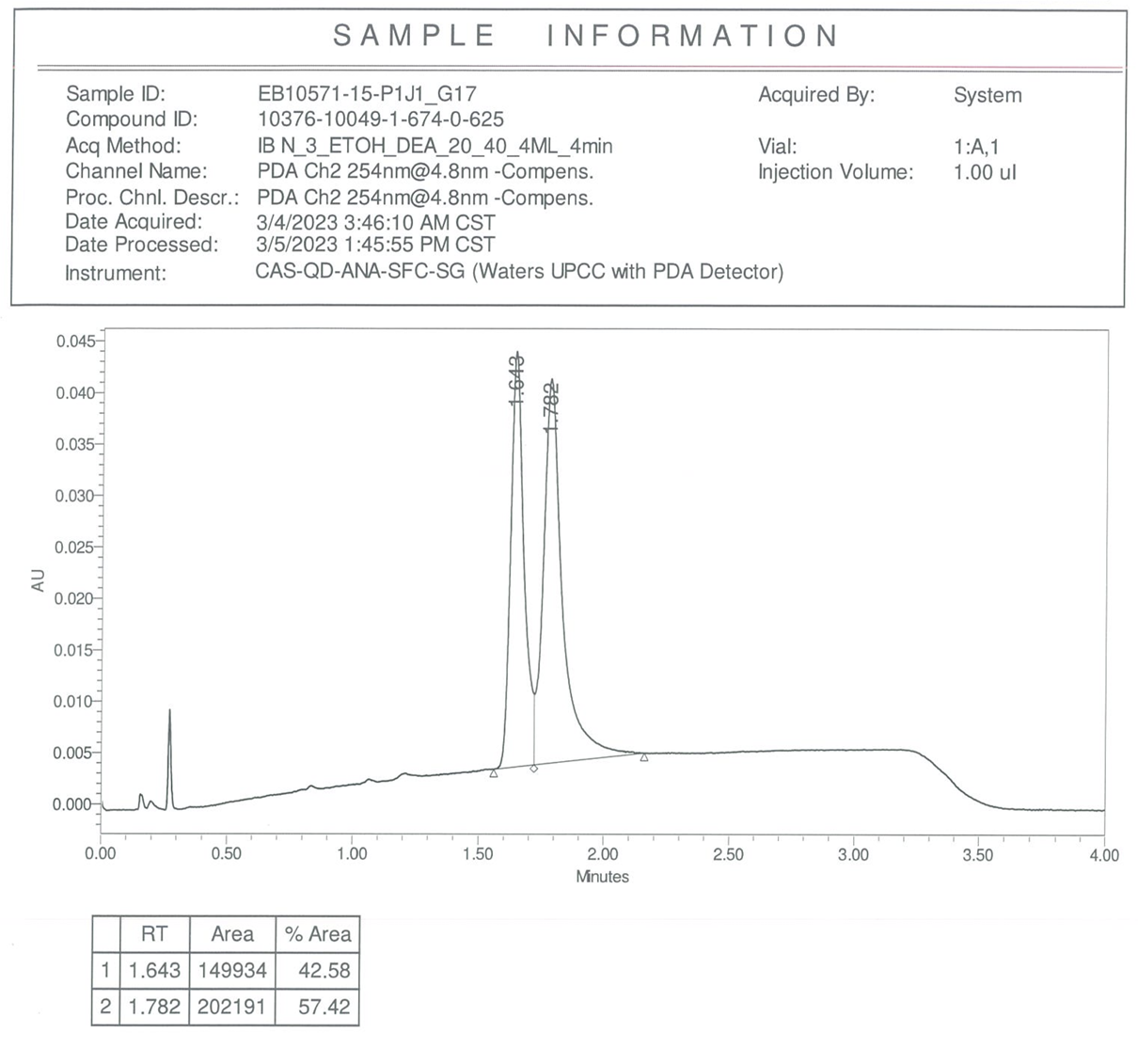

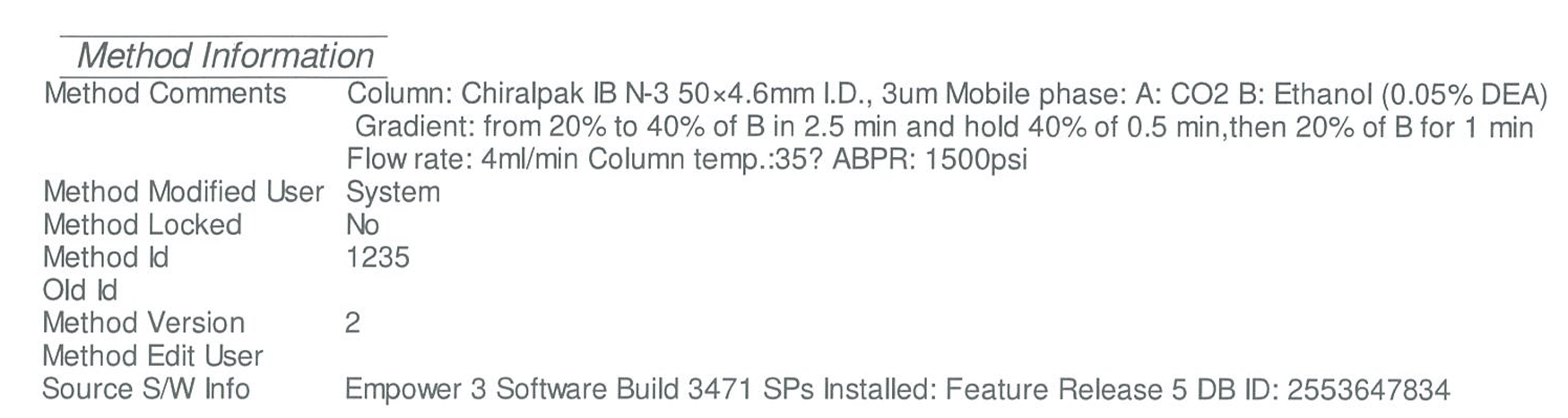

**Figure S10**. Chiral SFC report of **LAG-3** **Hit II**.

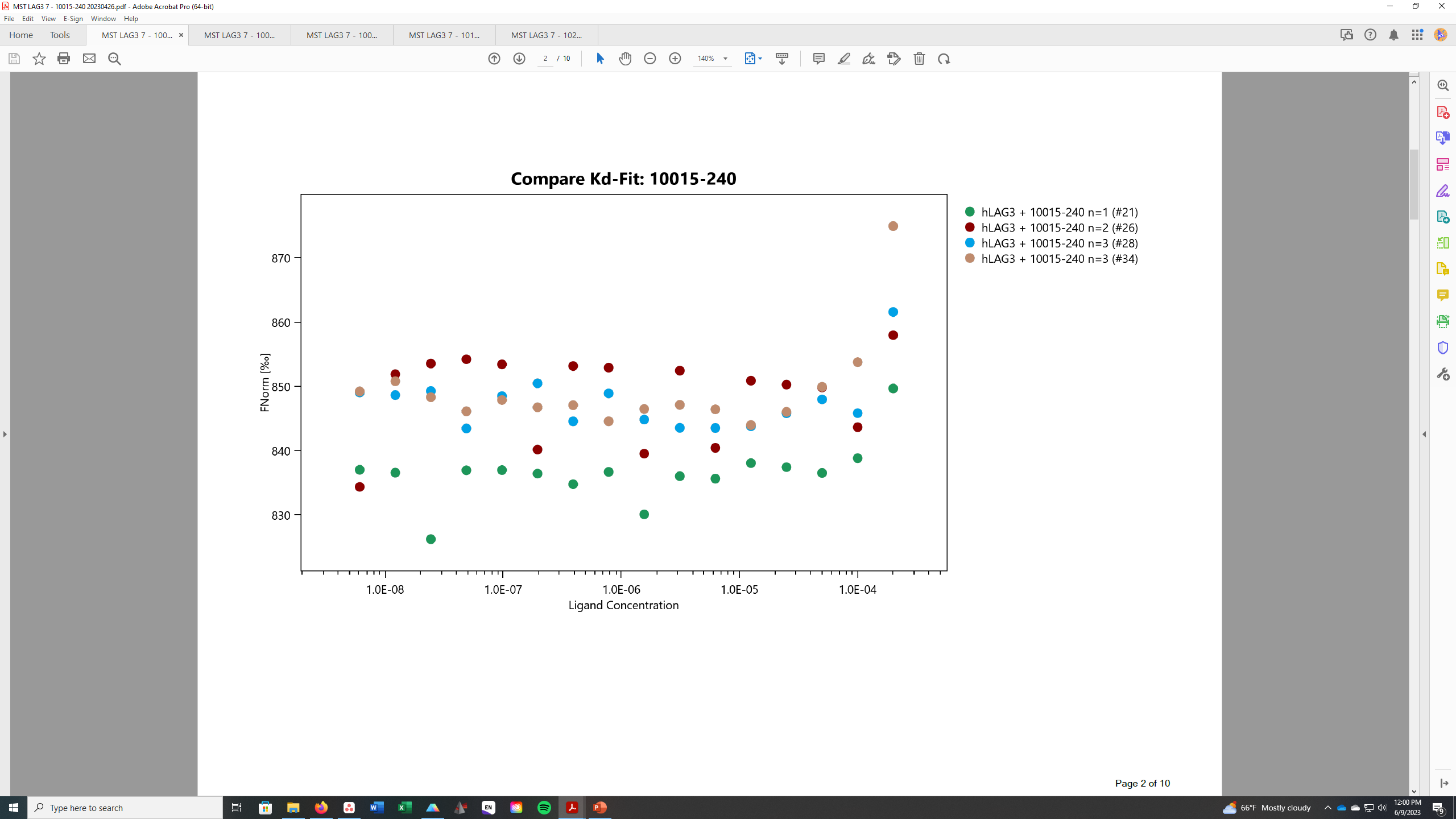

**Figure S11**. MST binding of **LAG-3** **Hit I** (increasing concentrations, n=4) to LAG-3.

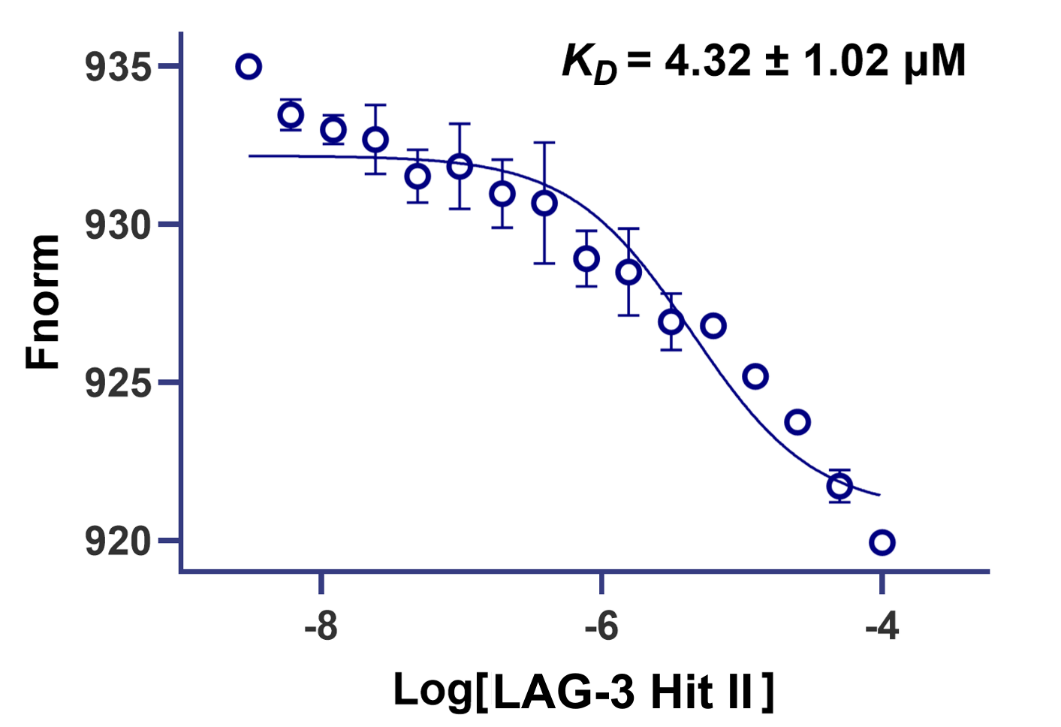

**Figure S12**. MST binding of **LAG-3** **Hit II** (increasing concentrations, n=3) to LAG-3. Binding was analyzed using nonlinear regression.

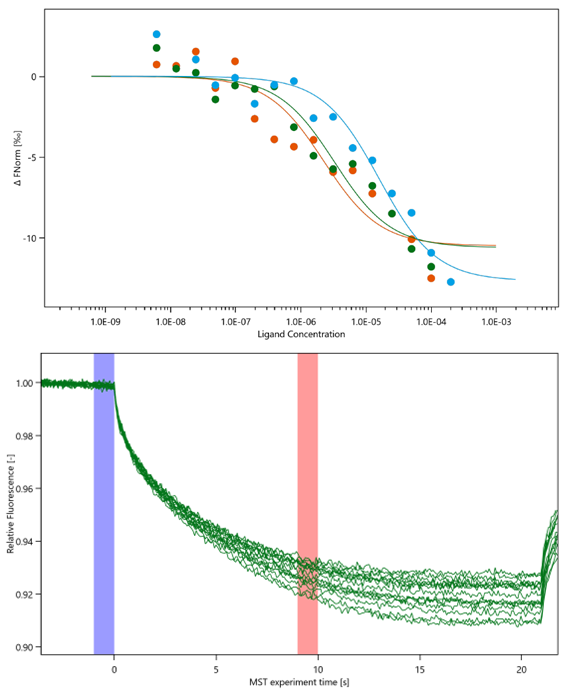

**Figure S13**. Three independent dose-response curves for the binding interaction between **LAG-3 Hit II** and LAG-3 (top). The corresponding MST time traces are shown on the bottom.

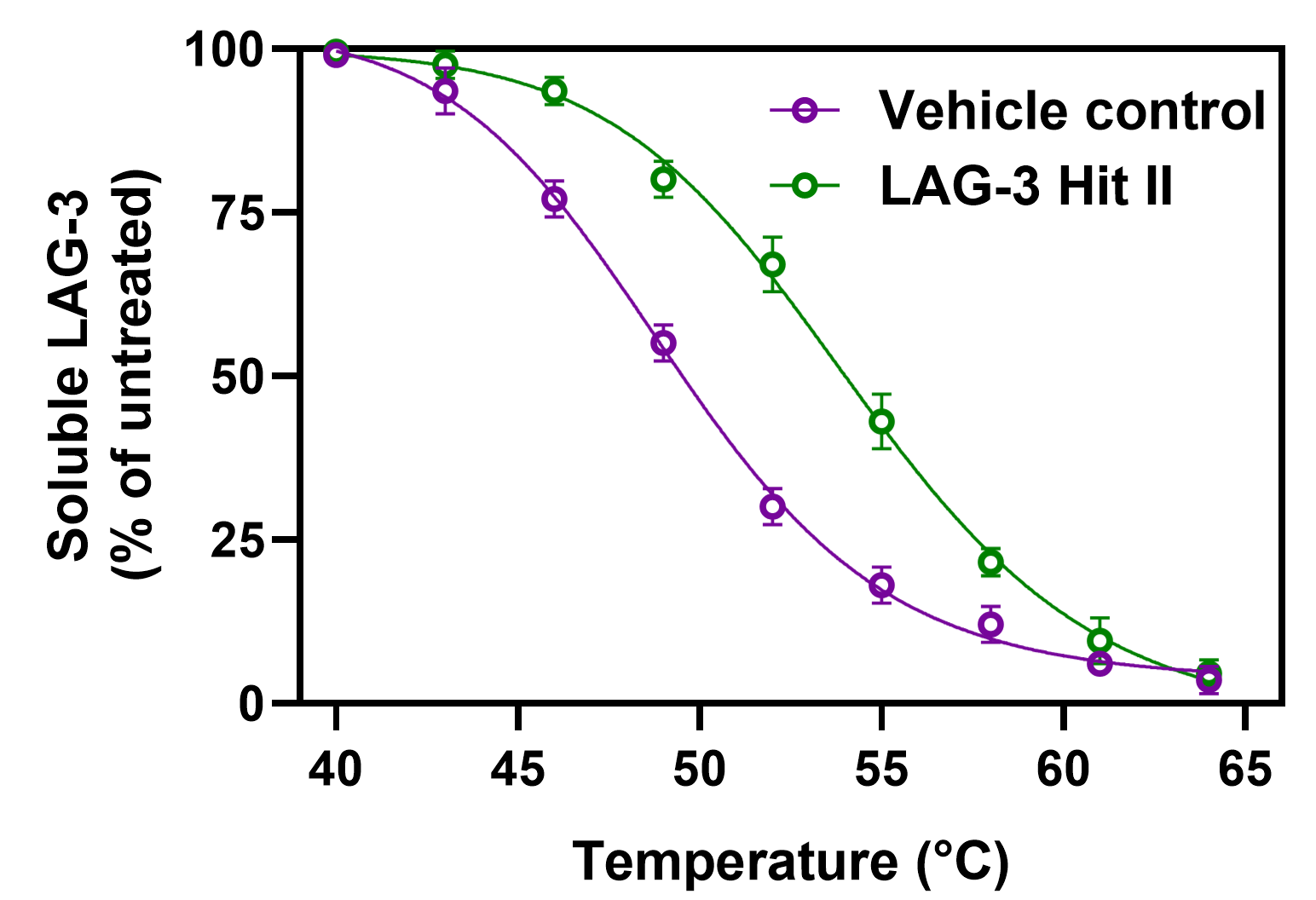

**Figure S14**. Cellular thermal shift assay (CETSA) for the binding of **LAG-3 Hit II** (25 μM) to LAG-3 in cell lysate of LAG-3 expressing Raji cells. Error bars represent standard deviation (n=5).

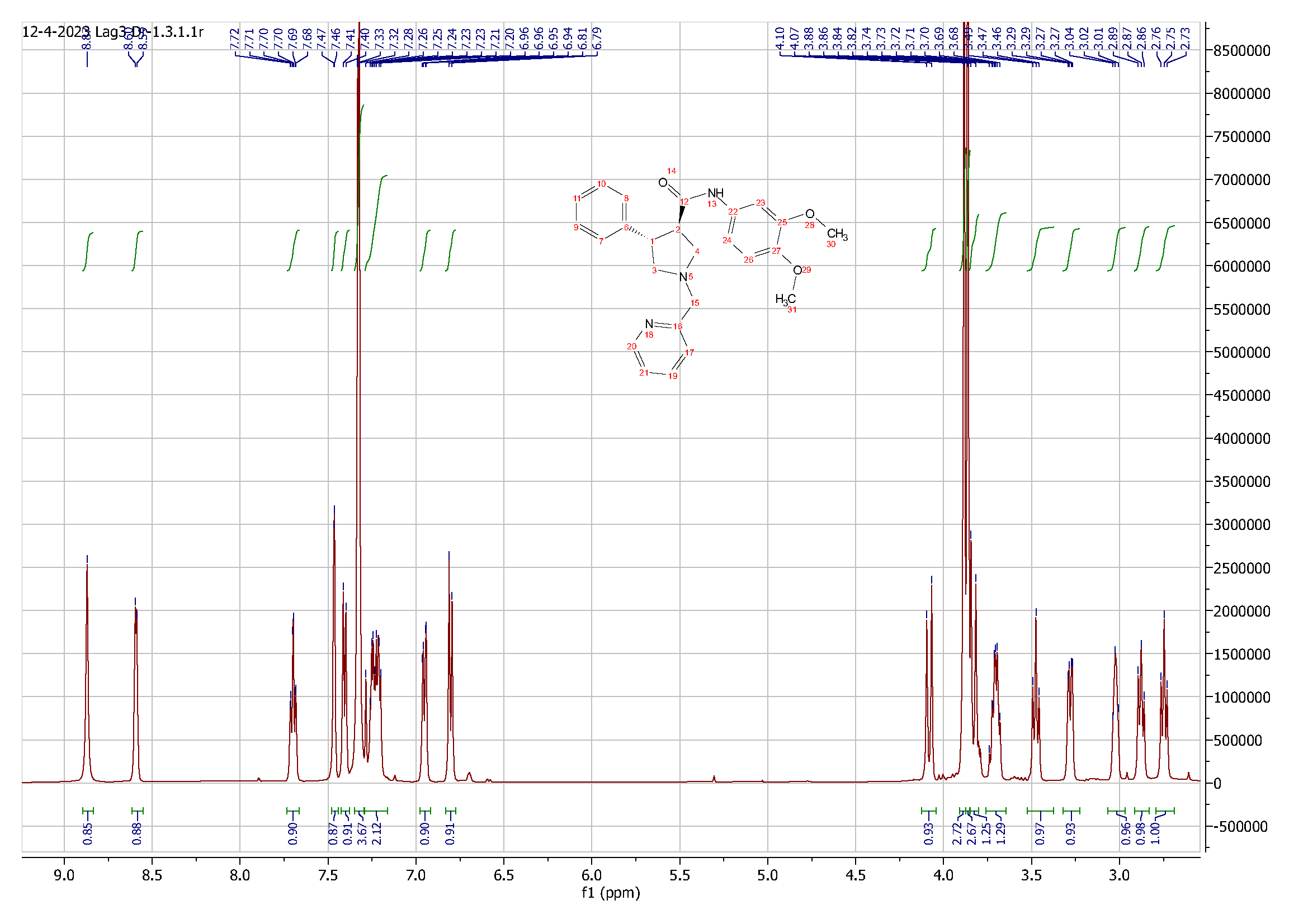
**Figure S15**. ^1^H NMR spectrum of compound **1**.

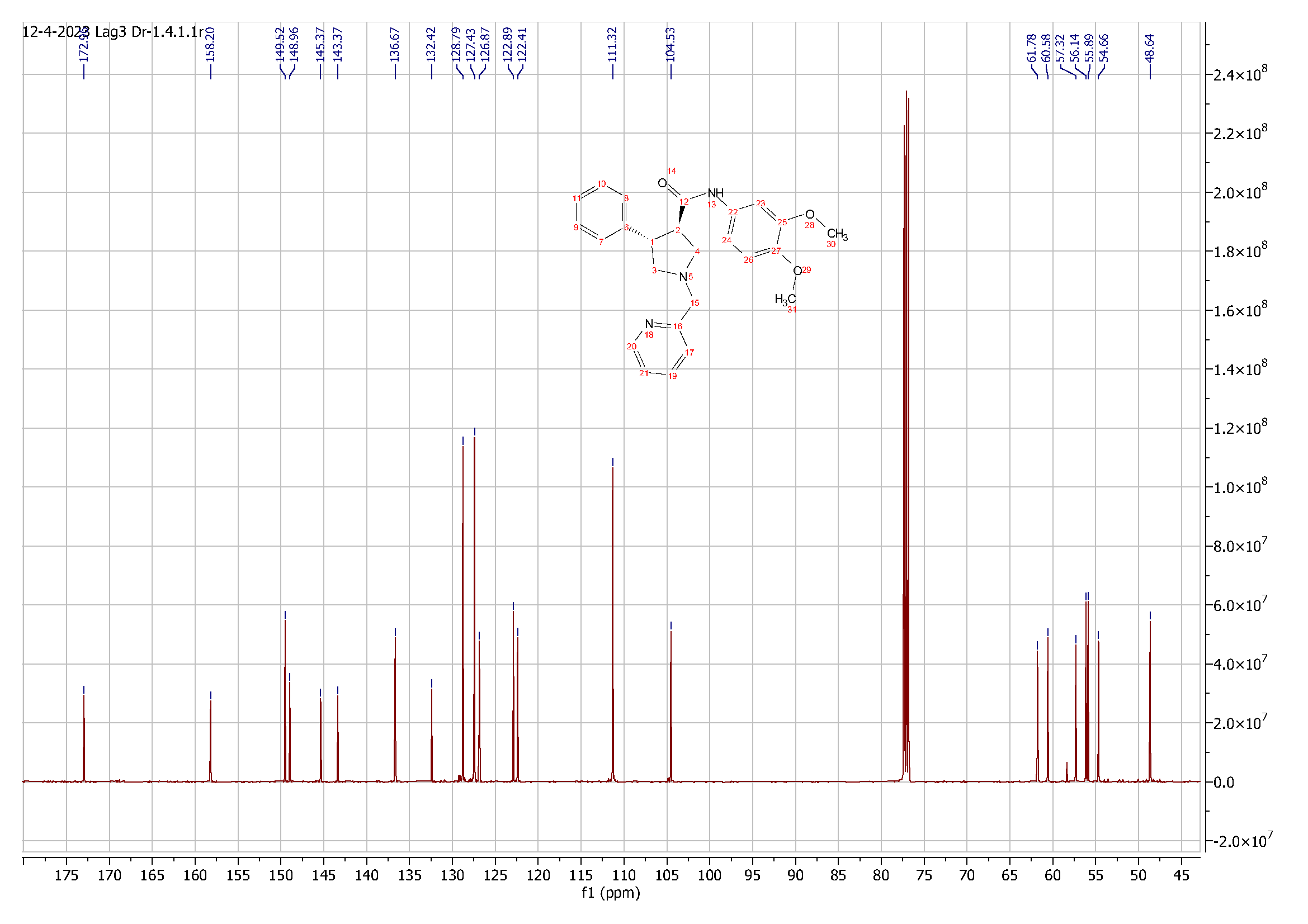

**Figure S16**. ^13^C NMR spectrum of compound **1**.

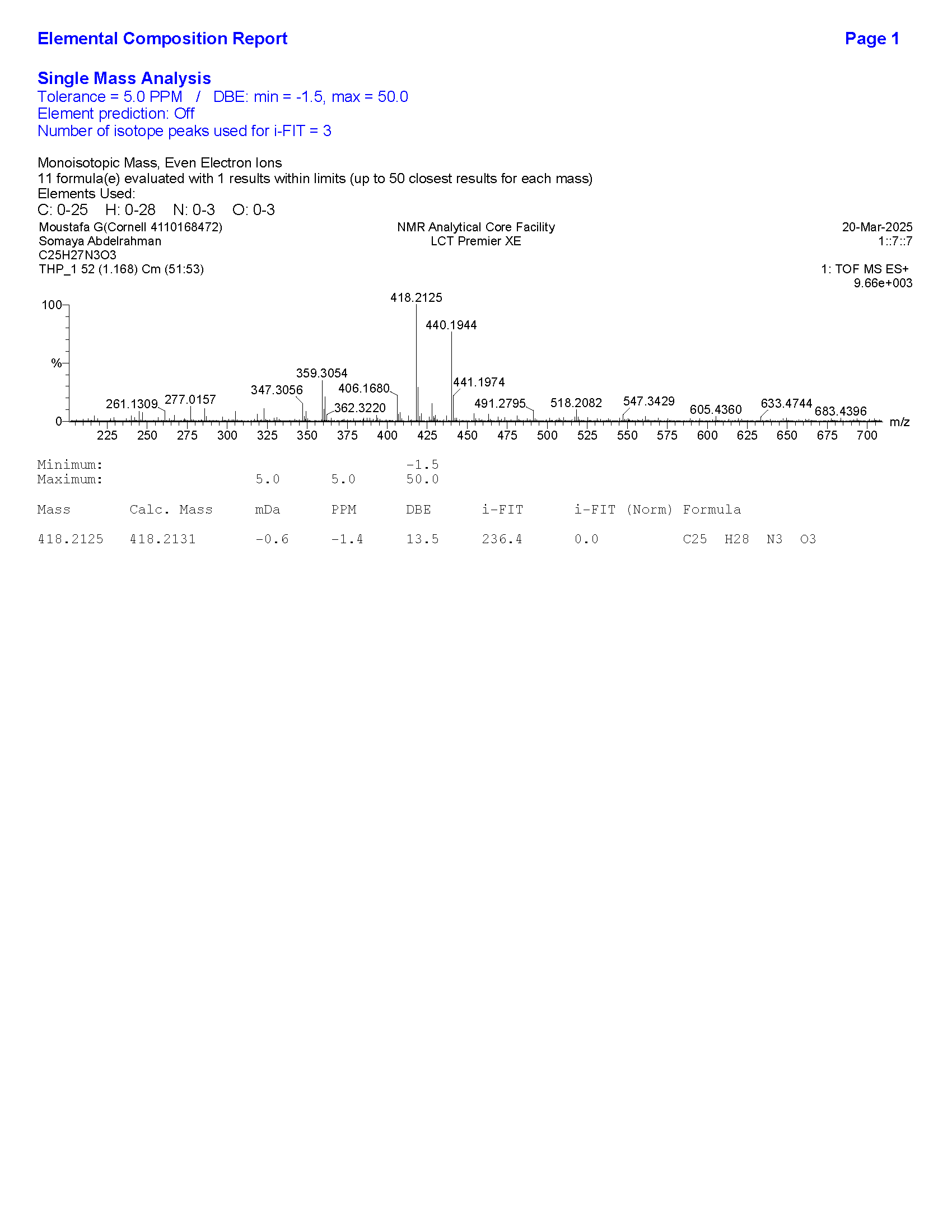

**Figure S17**. HRMS data for compound **1**.

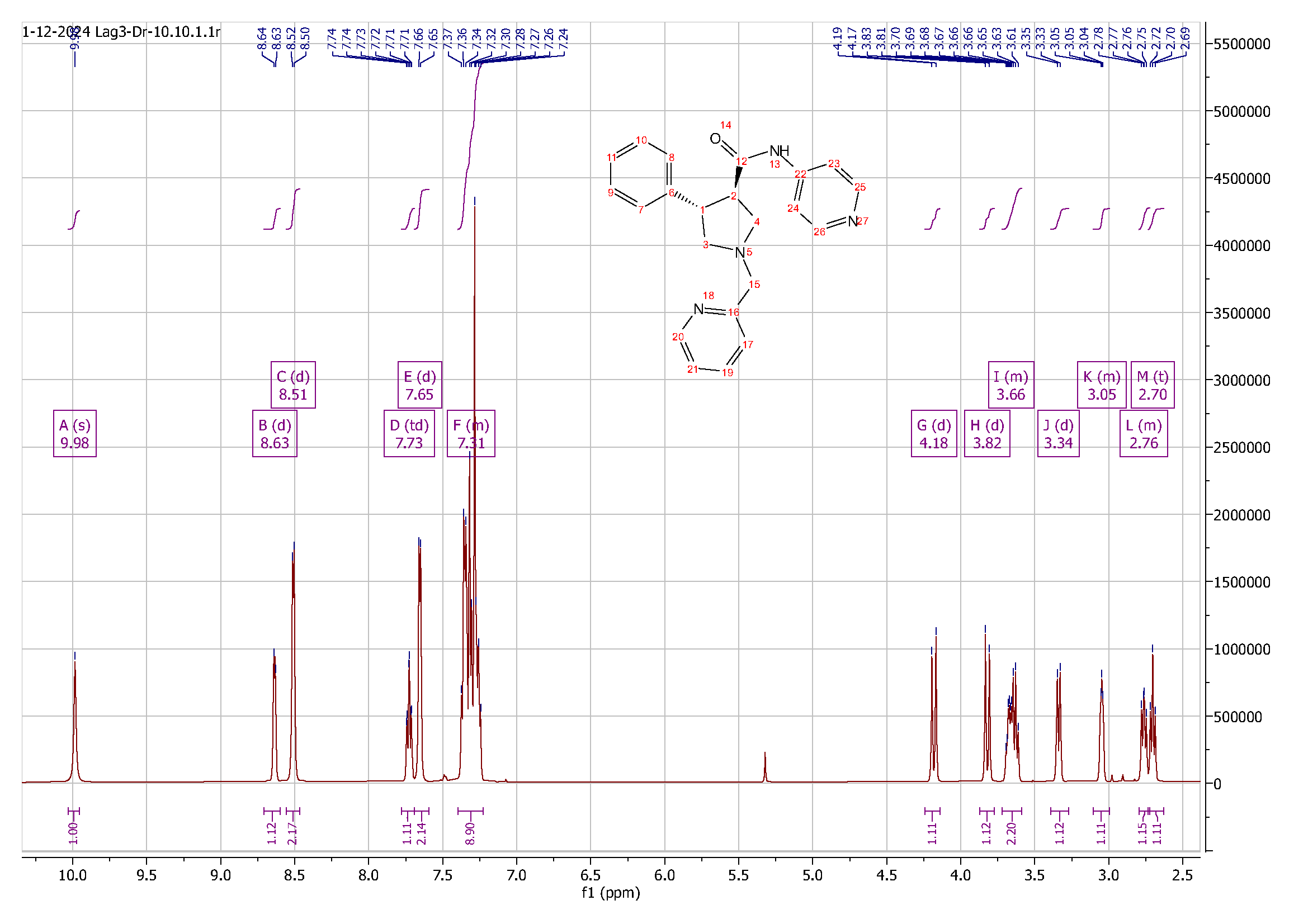
**Figure S18**. ^1^H NMR spectrum of compound **2**.

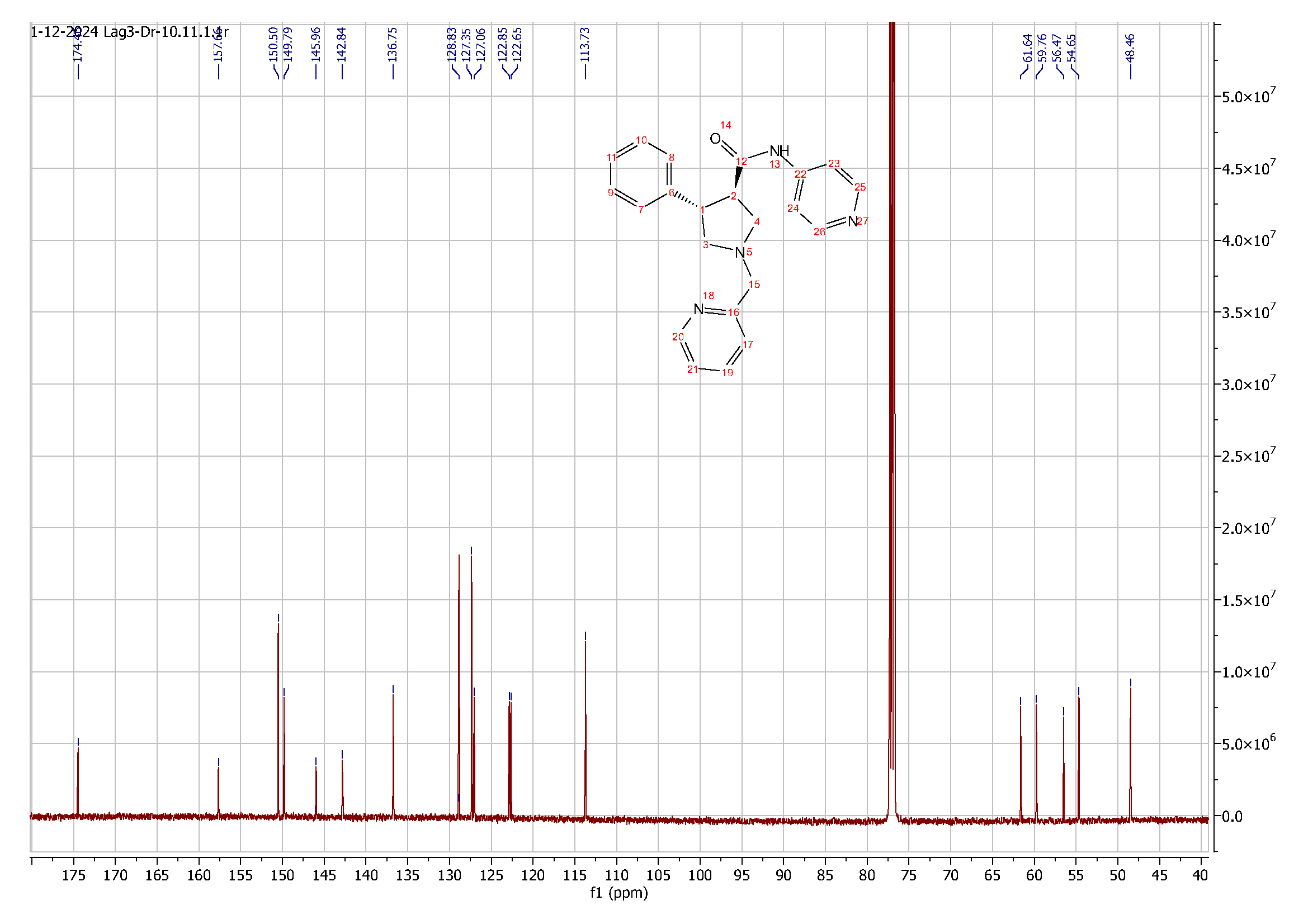

**Figure S19**. ^13^C NMR spectrum of compound **2**.

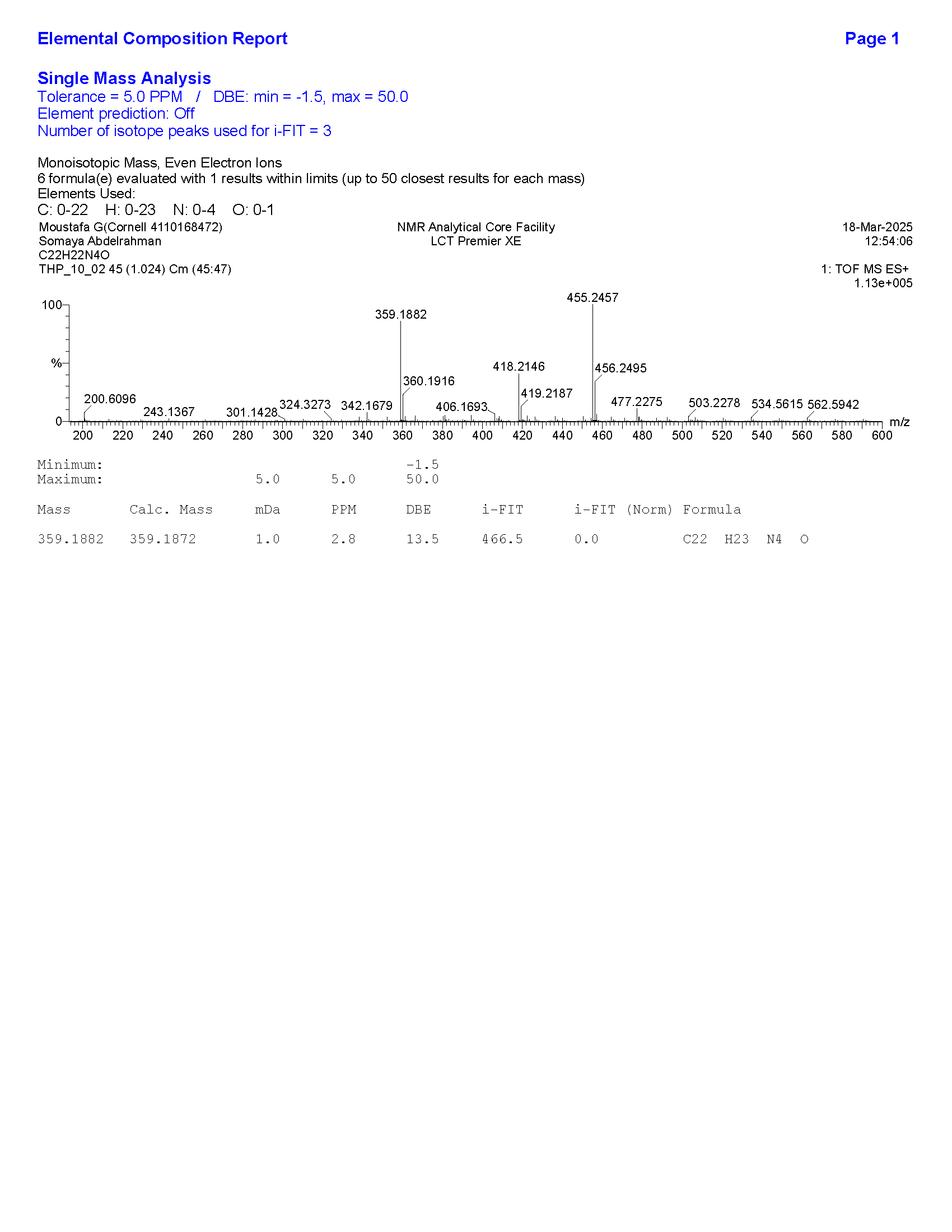

**Figure S20**. HRMS data for compound **2**.

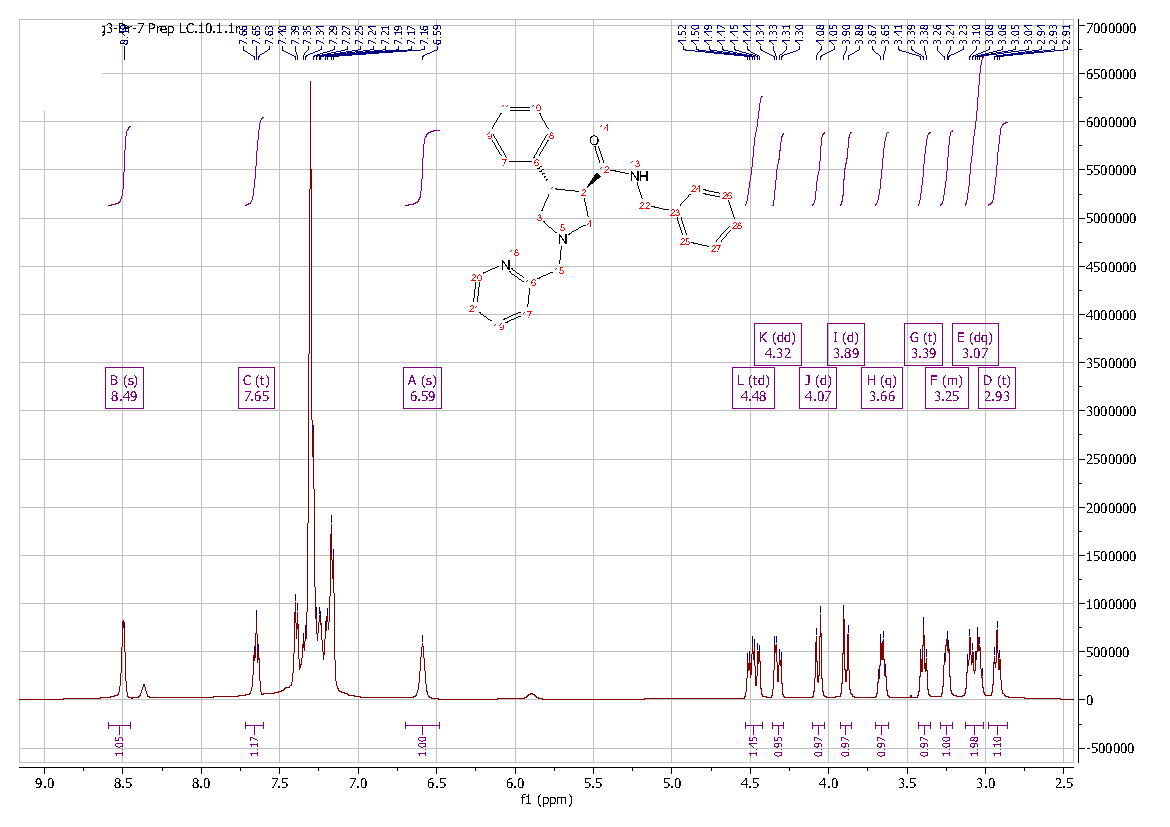
**Figure S21**. ^1^H NMR spectrum of compound **3**.

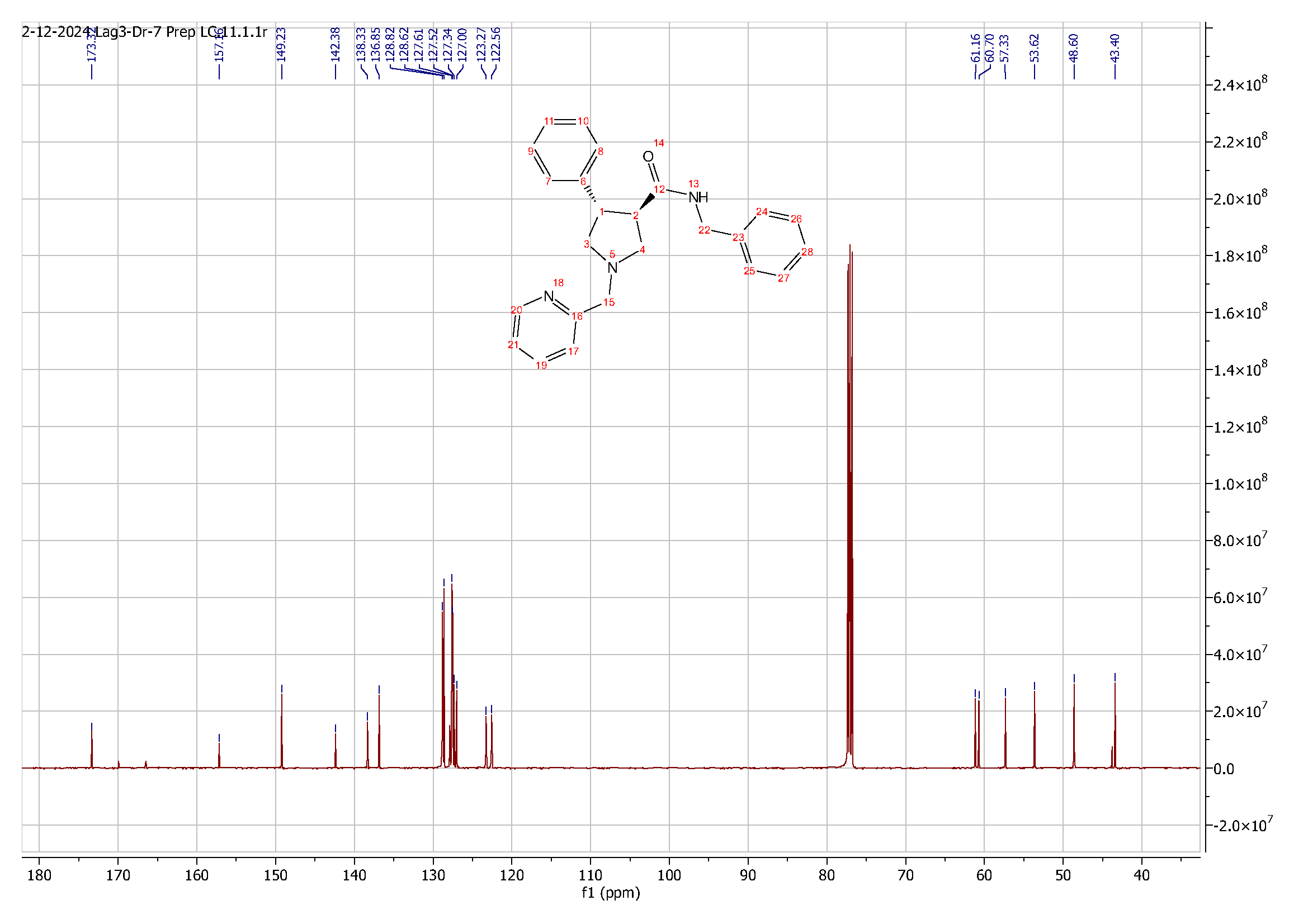

**Figure S22**. ^13^C NMR spectrum of compound **3**.

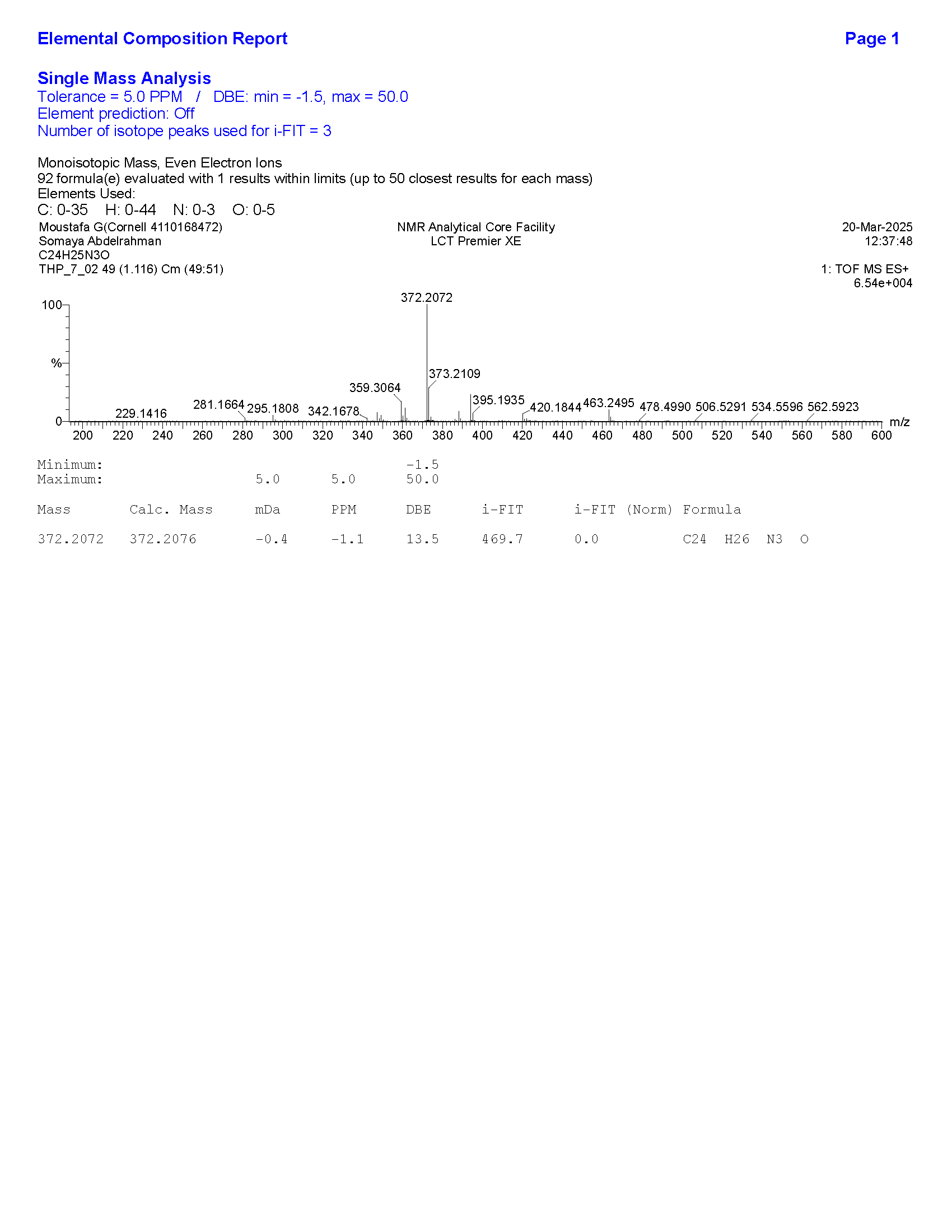

**Figure S23**. HRMS data for compound **3**.

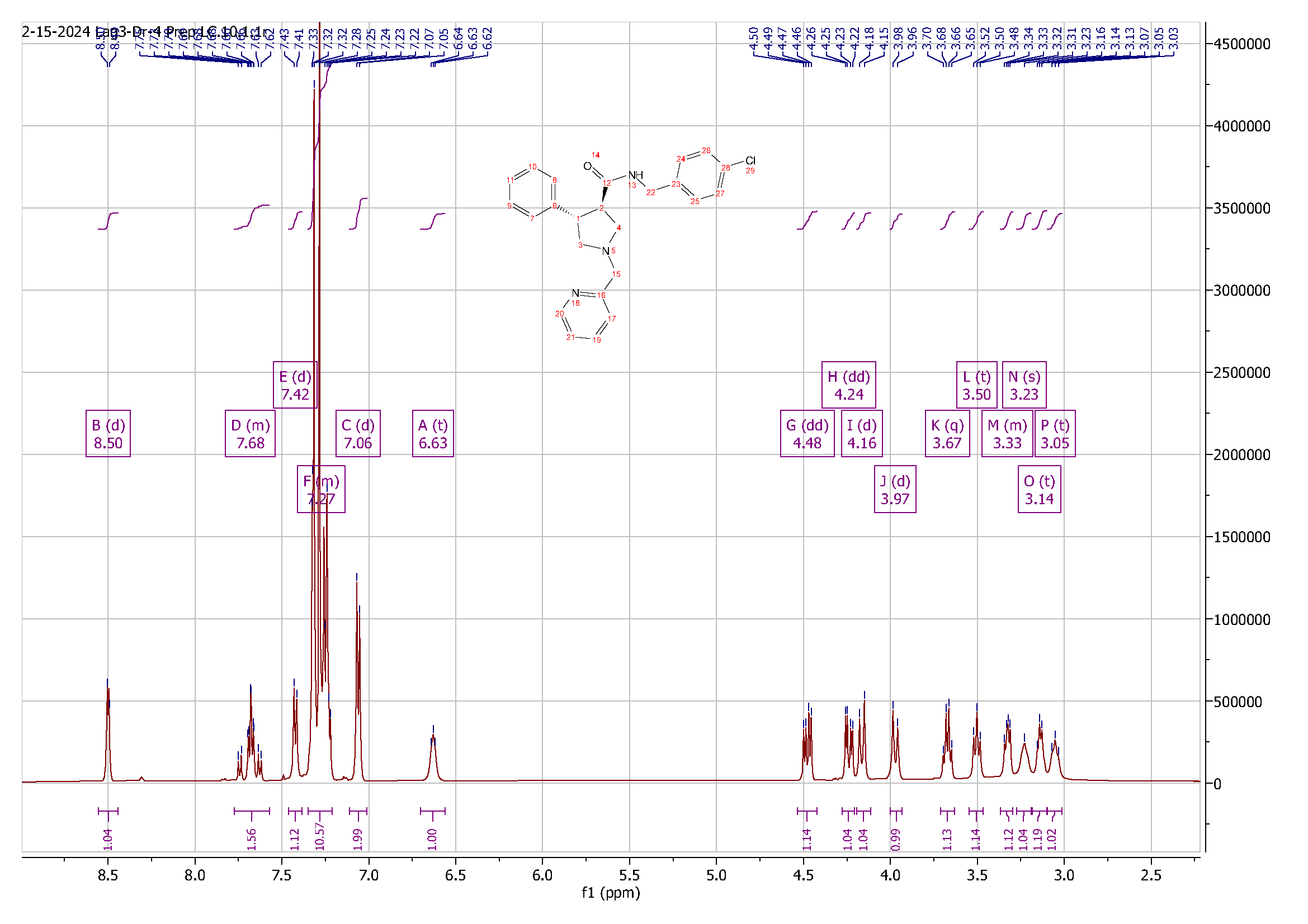
**Figure S24**. ^1^H NMR spectrum of compound **4**.

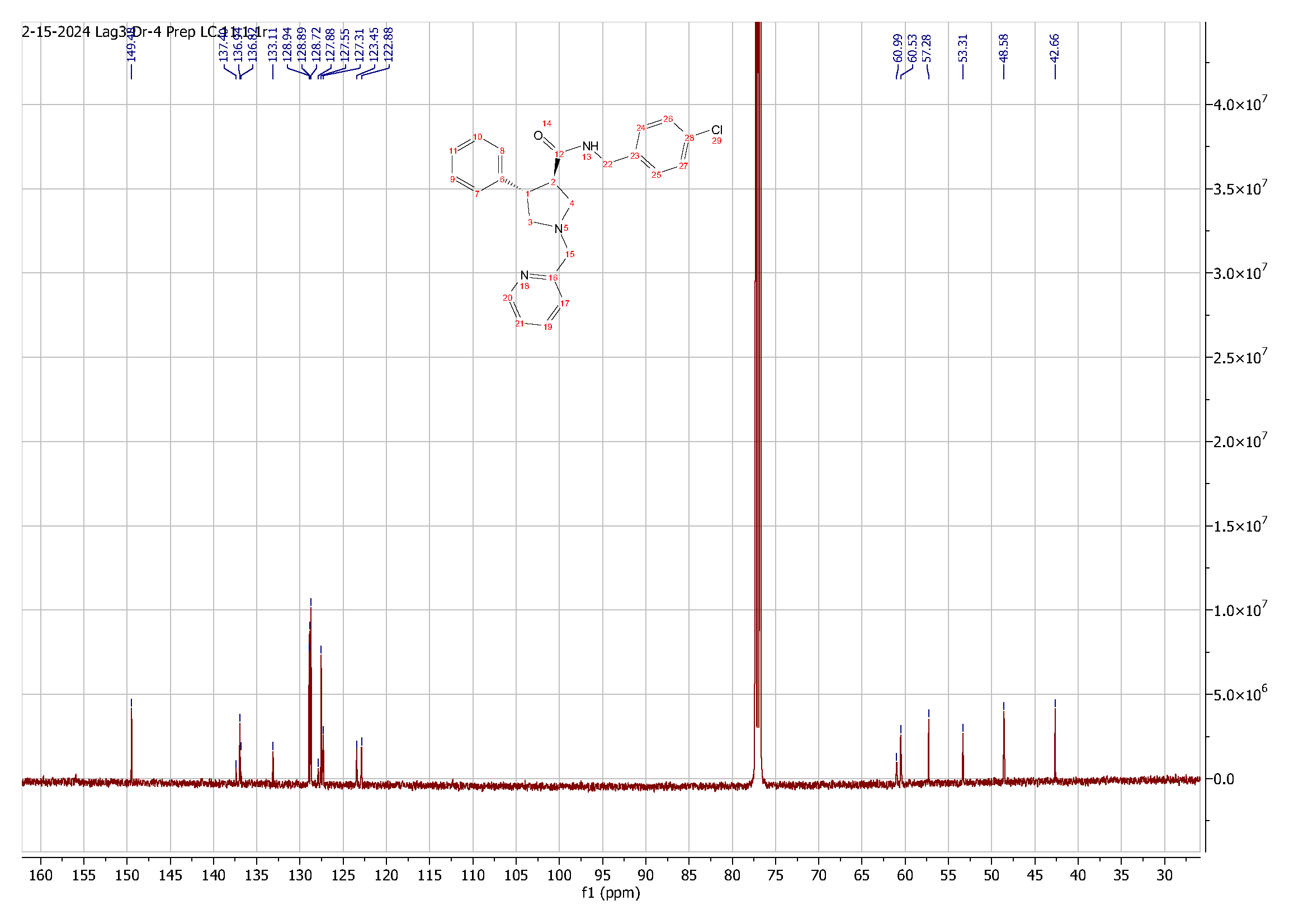

**Figure S25**. ^13^C NMR spectrum of compound **4**.

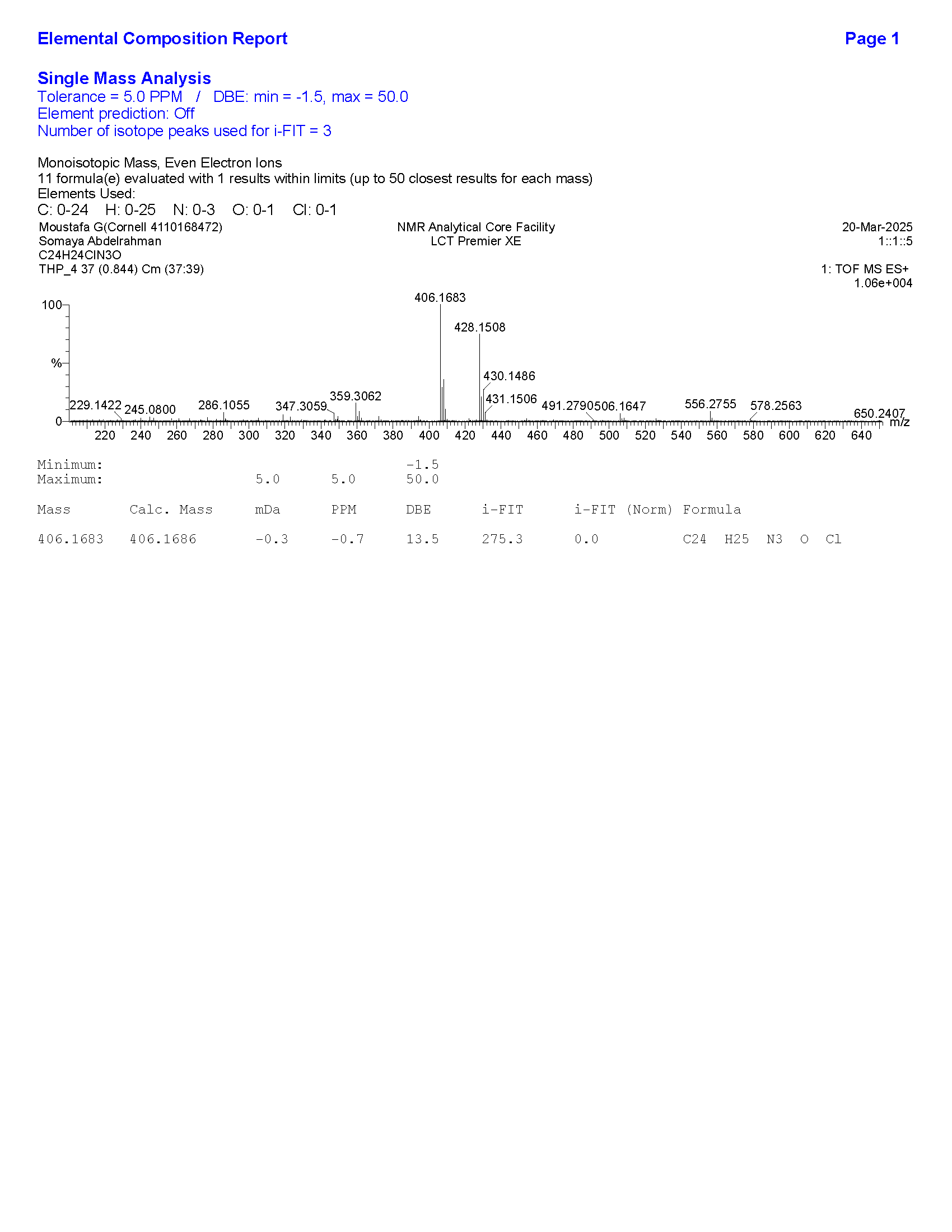

**Figure S26**. HRMS data for compound **4**.

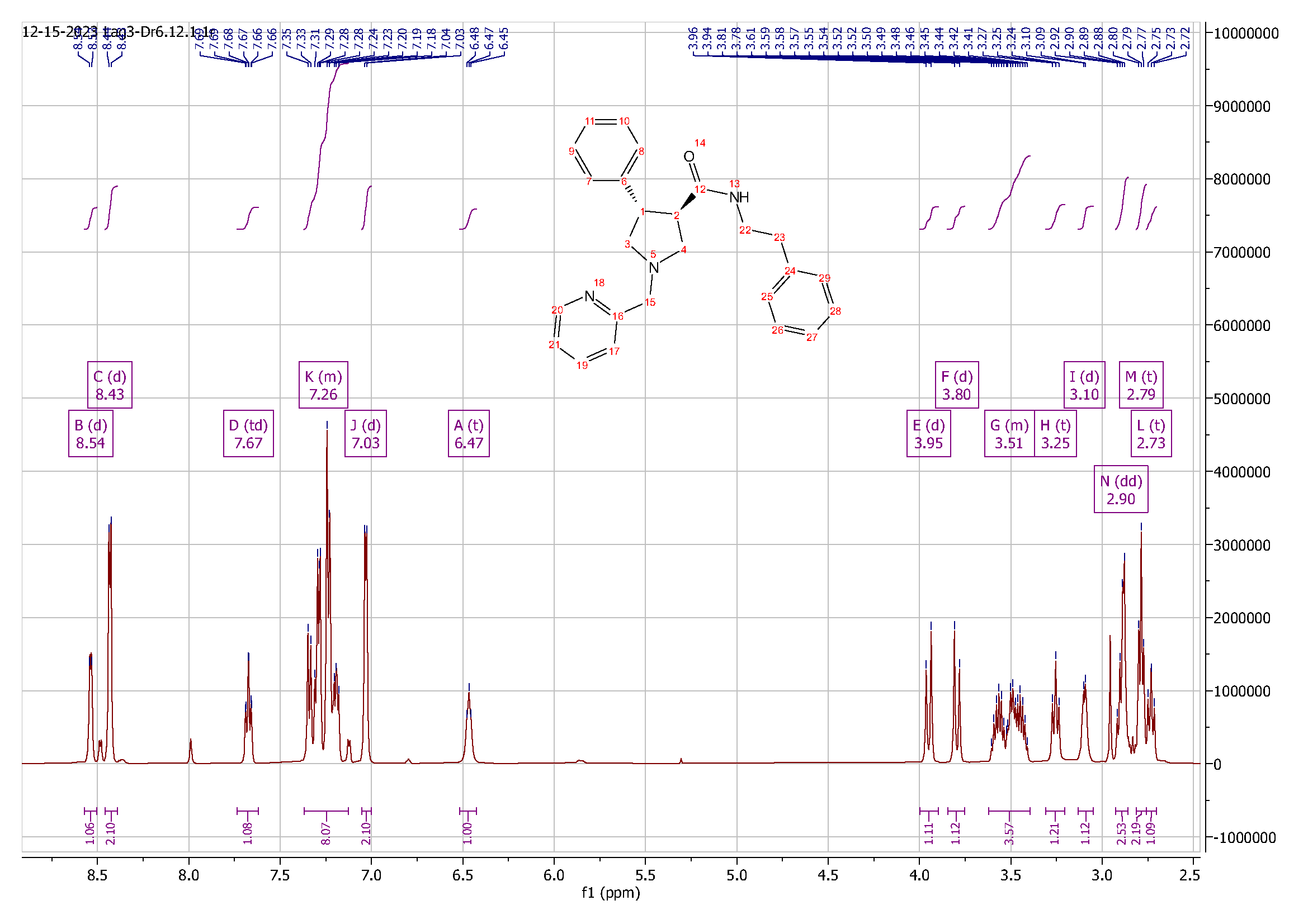
 **Figure S27**. ^1^H NMR spectrum of compound **5**.
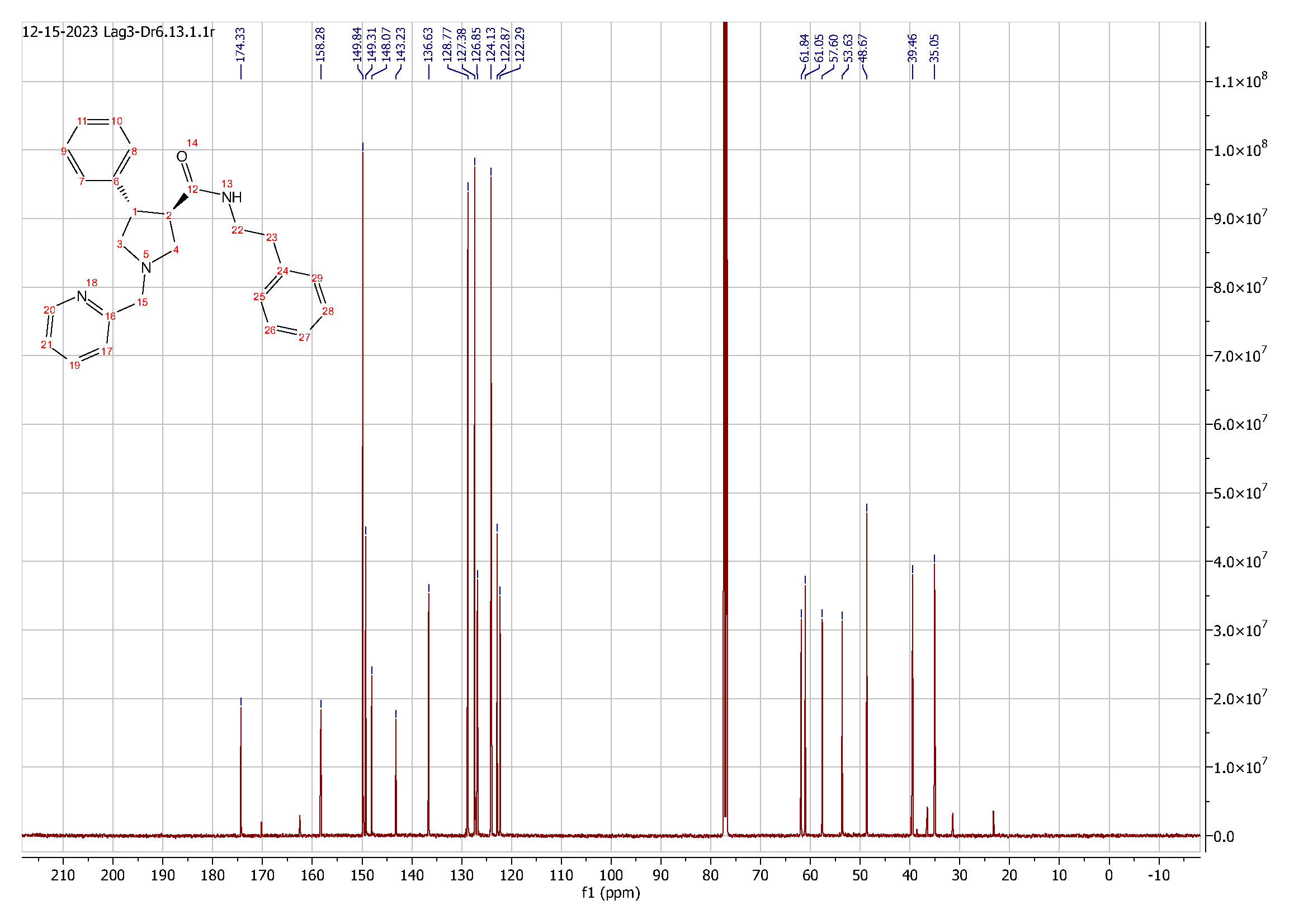
 **Figure S28**. ^13^C NMR spectrum of compound **5**.

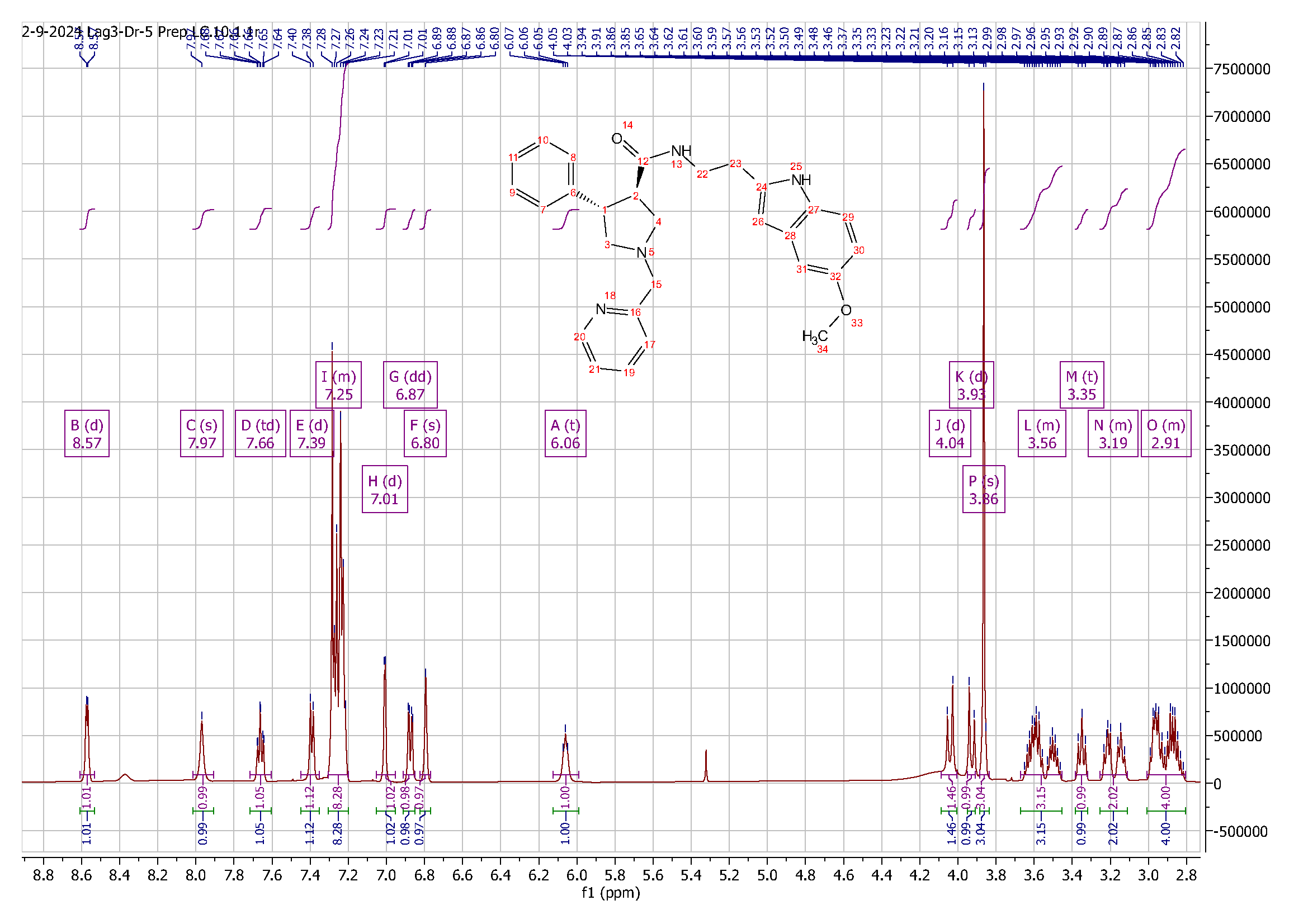

**Figure S29**. ^1^H NMR spectrum of compound **6**.

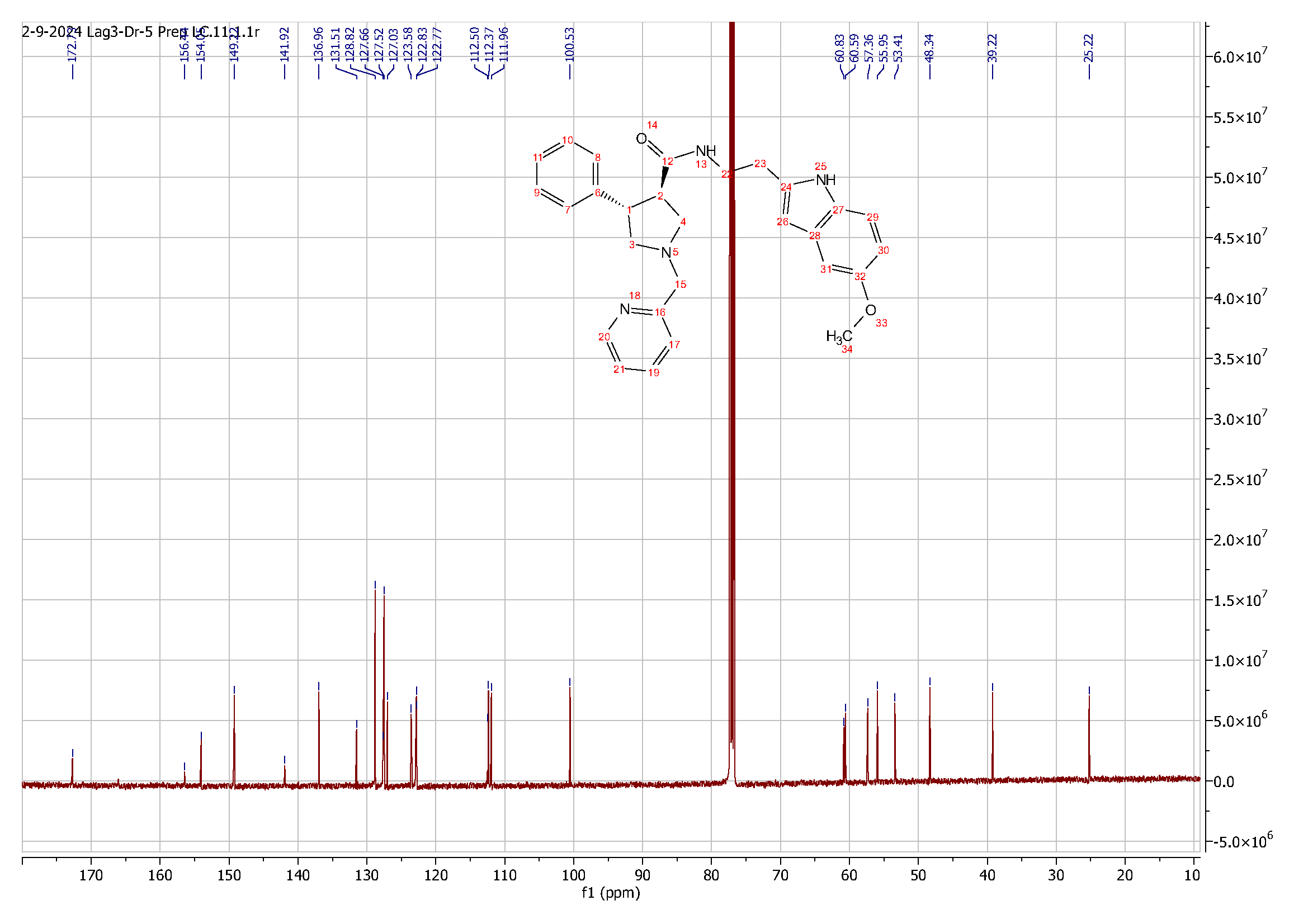

**Figure S30**. ^13^C NMR spectrum of compound **6**.

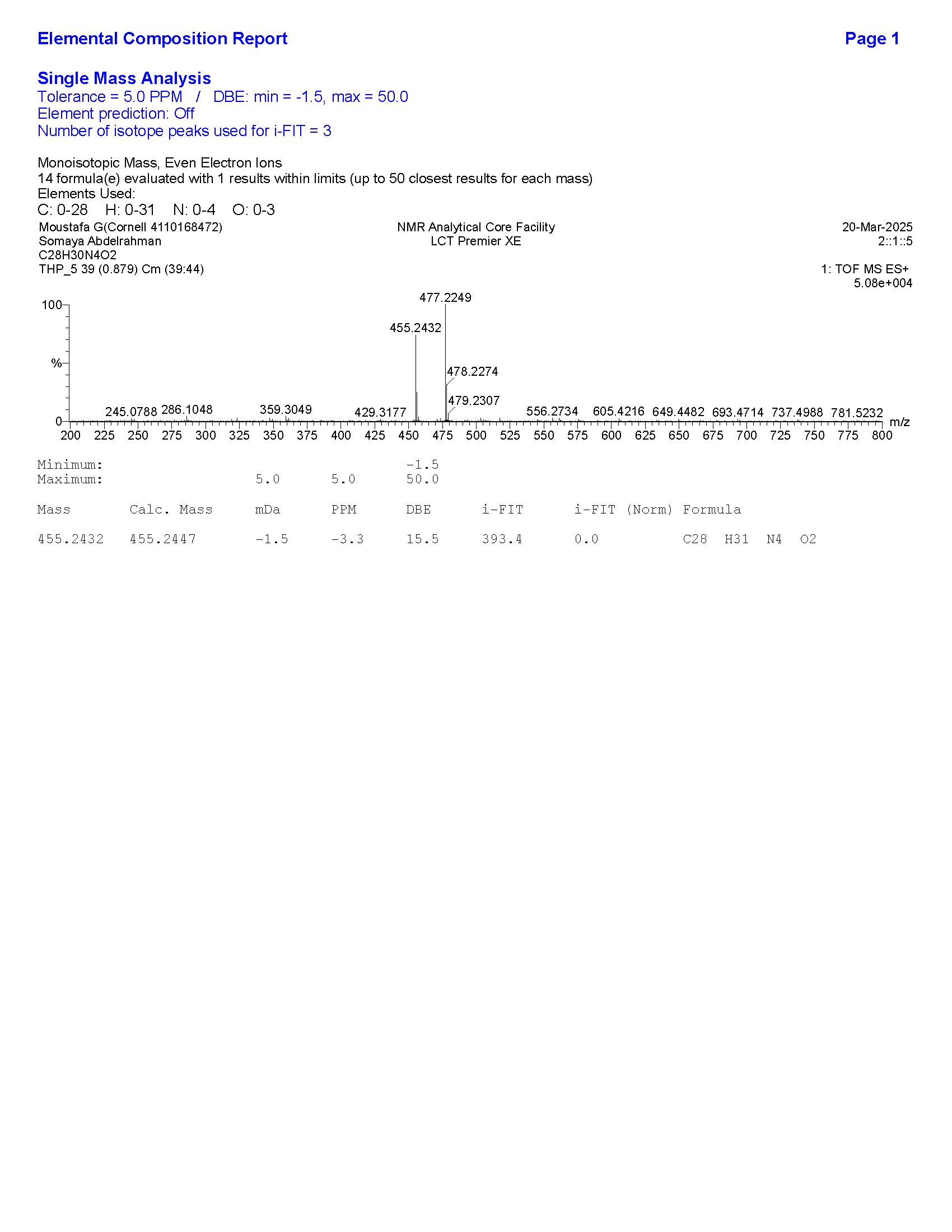

**Figure S31**. HRMS data for compound **6**.

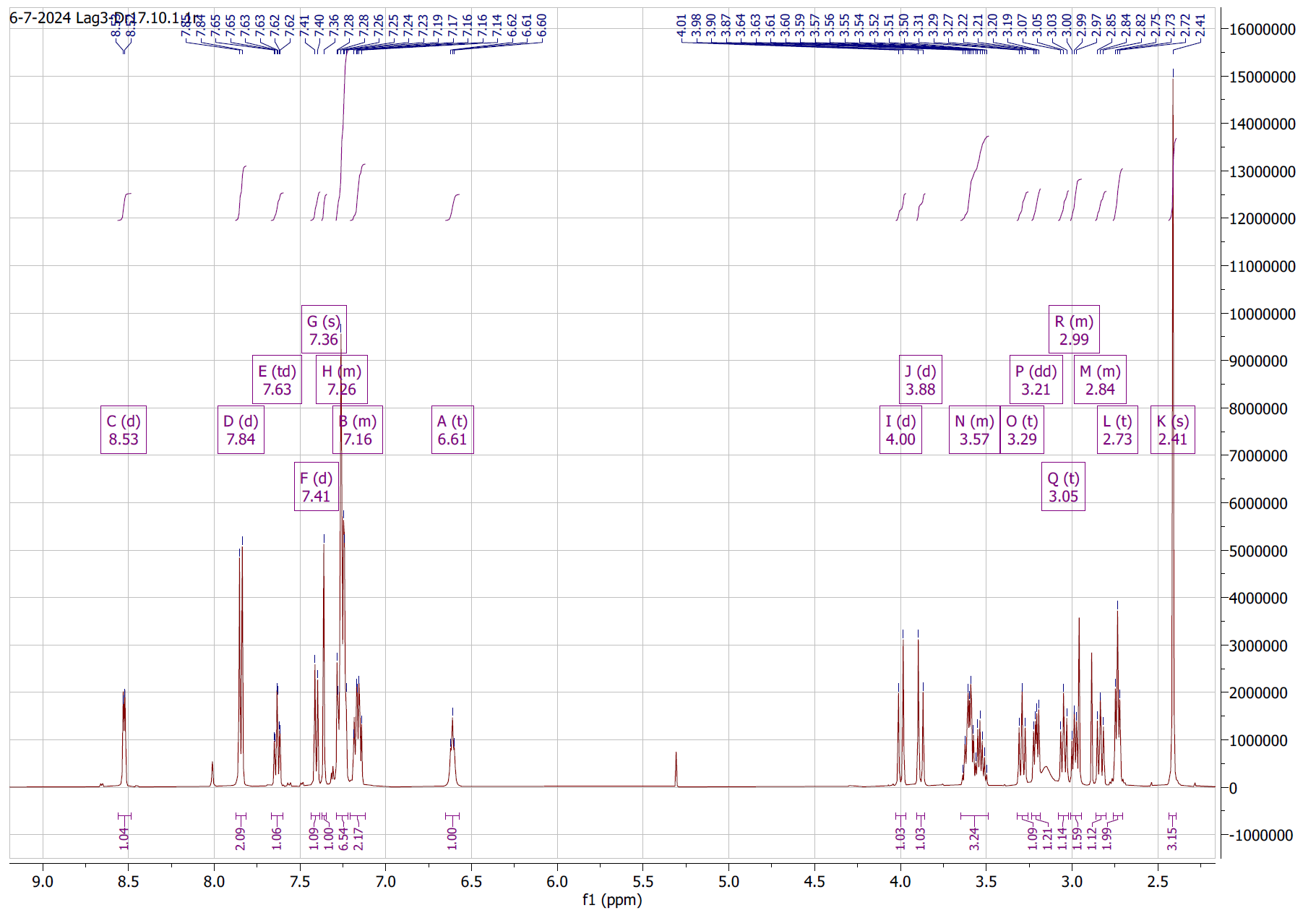

**Figure S32**. ^1^H NMR spectrum of compound **7**.

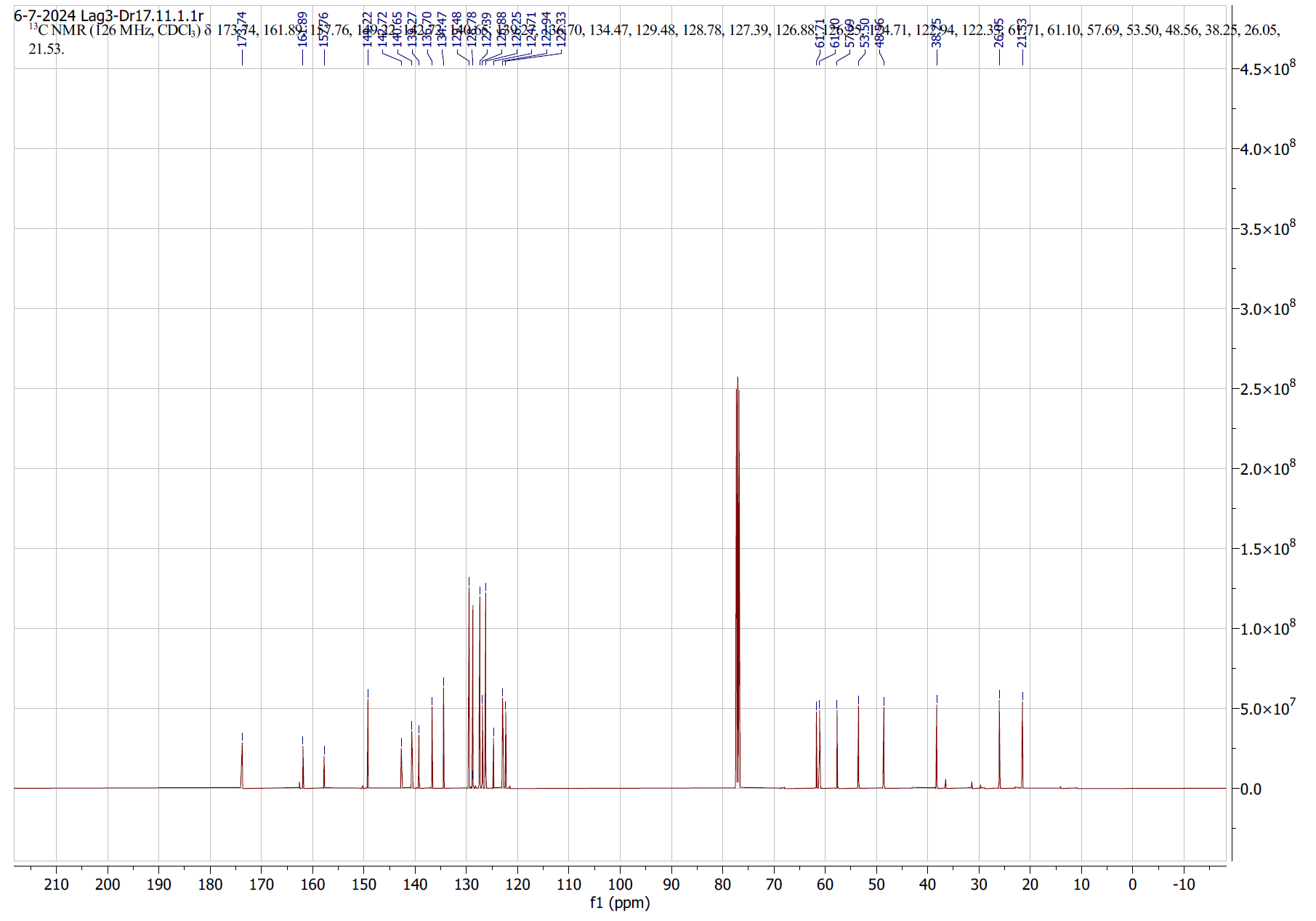

**Figure S33**. ^13^C NMR spectrum of compound **7**.

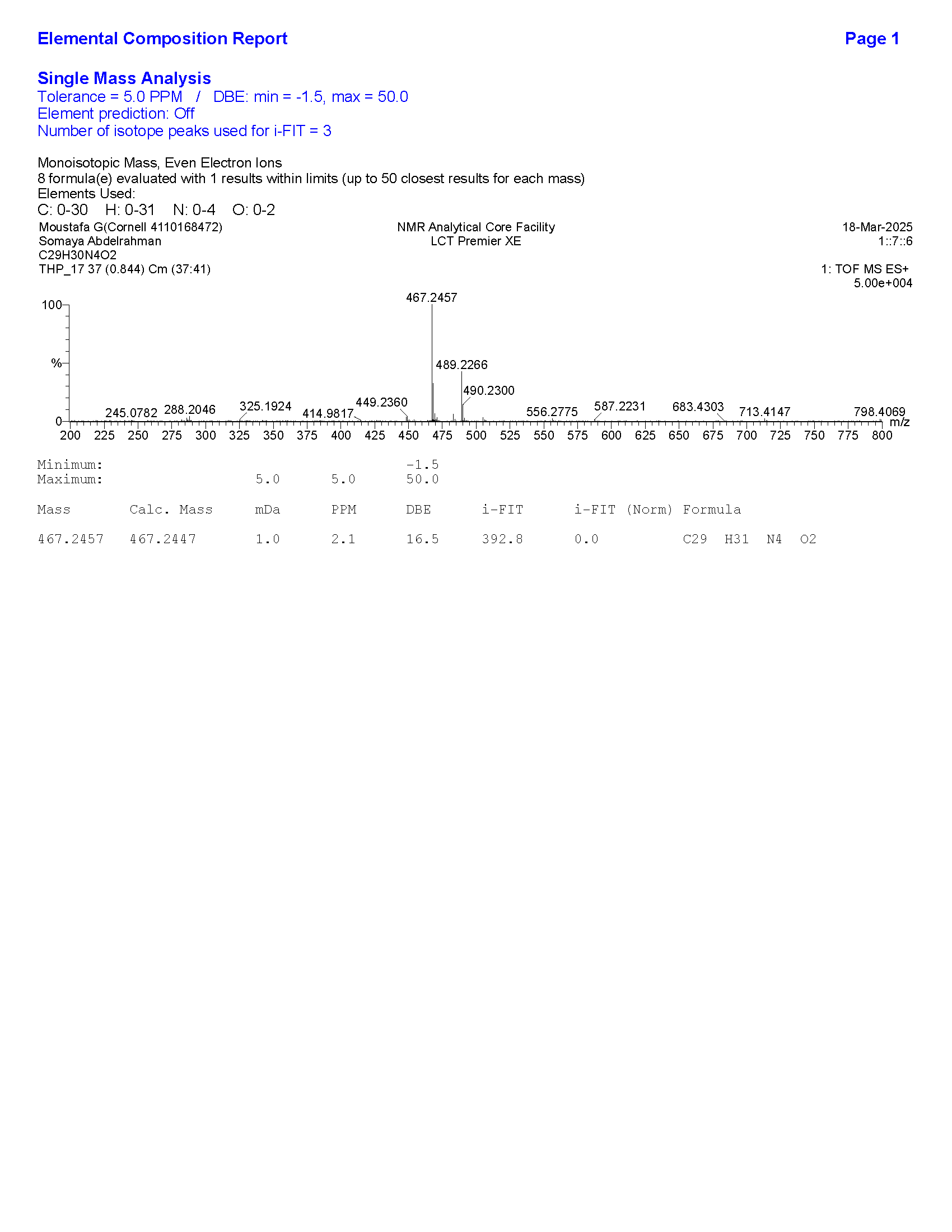

**Figure S34**. HRMS data for compound **7**.

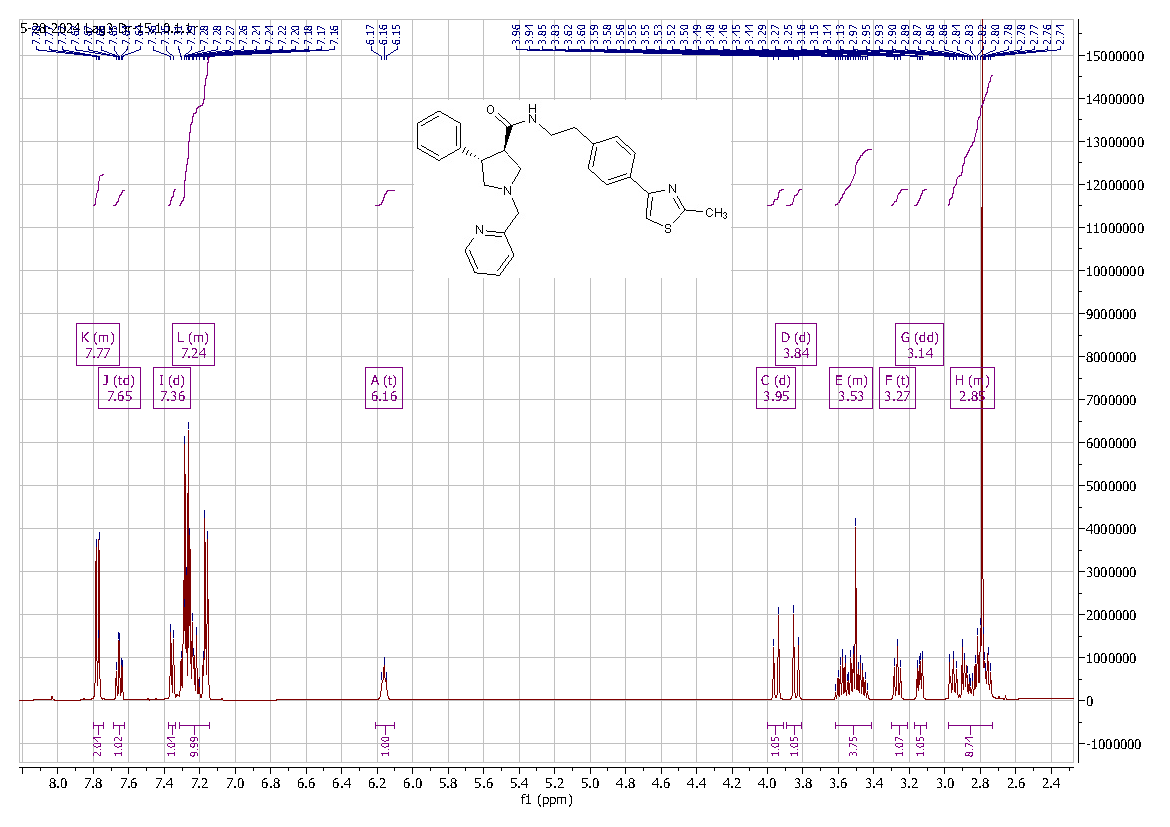

**Figure S35**. ^1^H NMR spectrum of compound **8**.

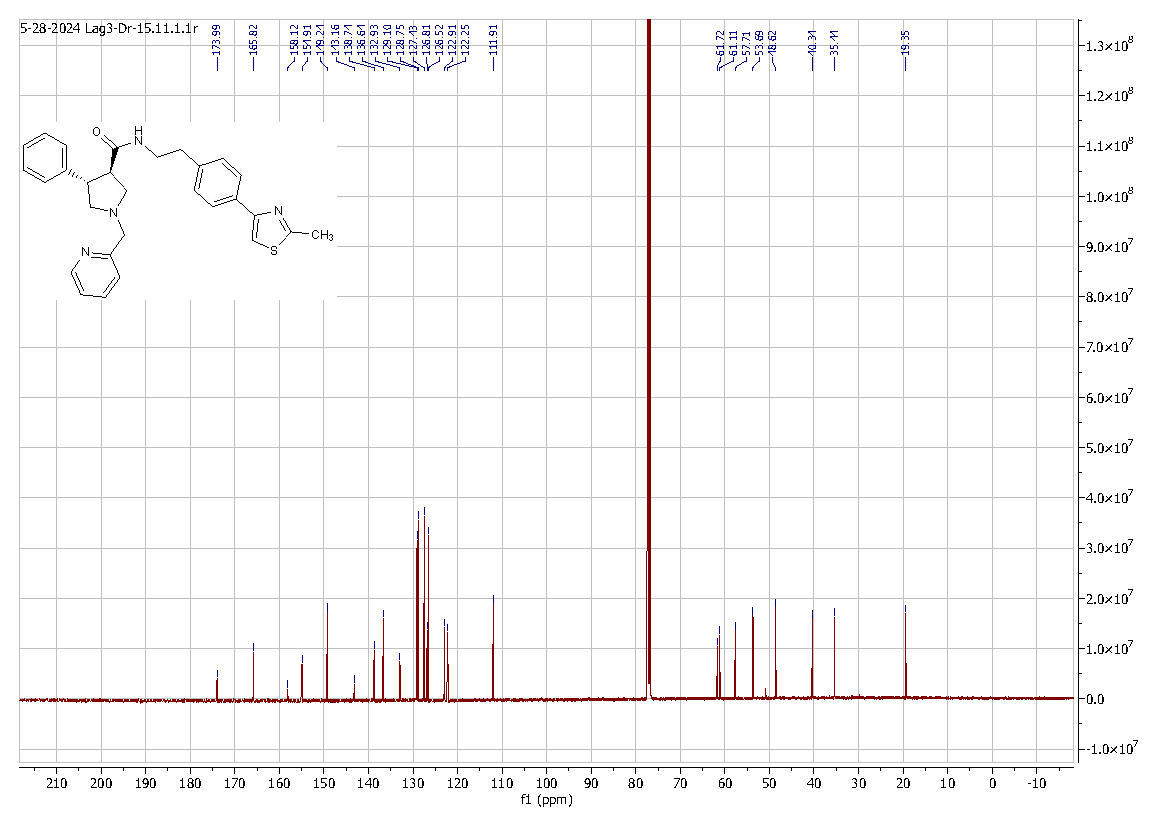

**Figure S36**. ^13^C NMR spectrum of compound **8**.

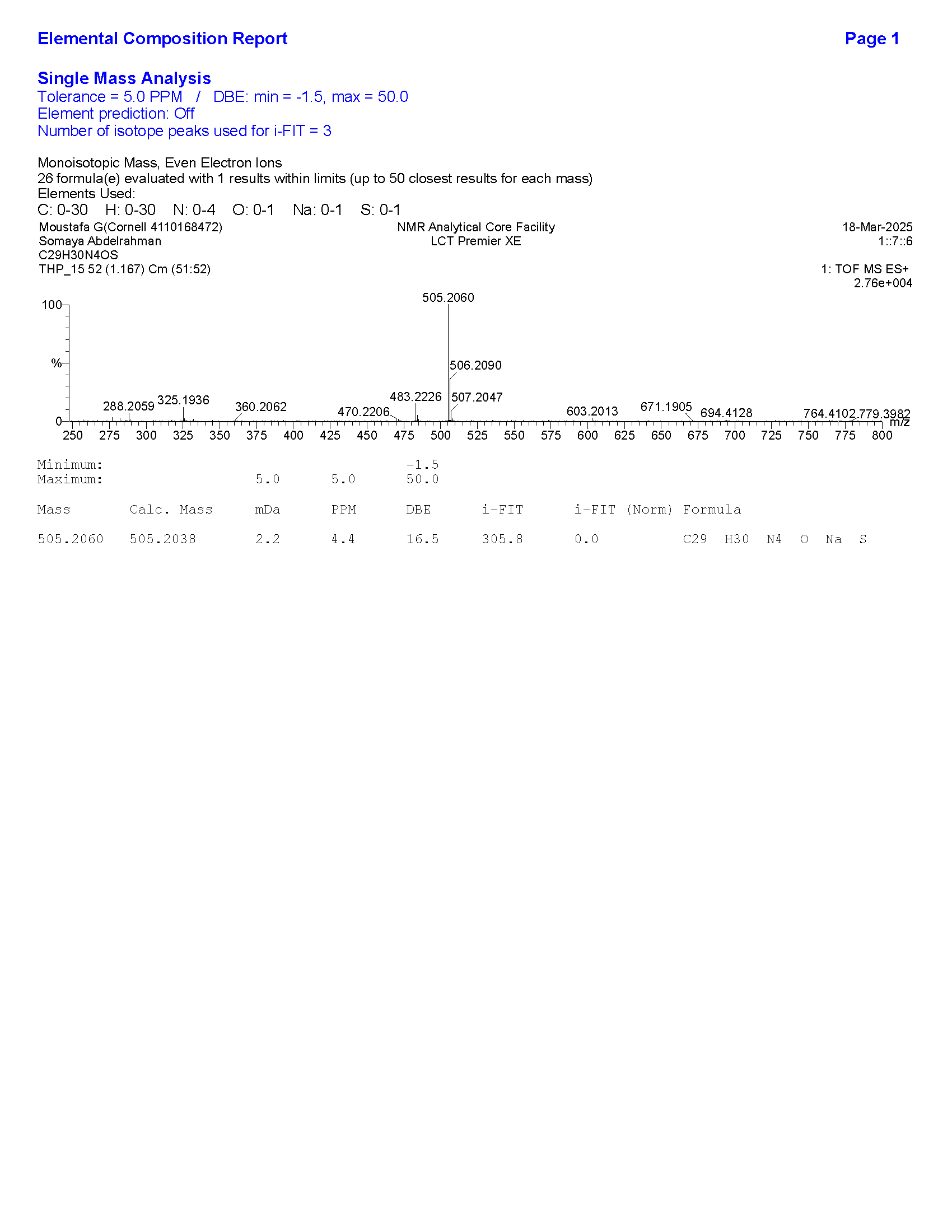

**Figure S37**. HRMS data for compound **8**.

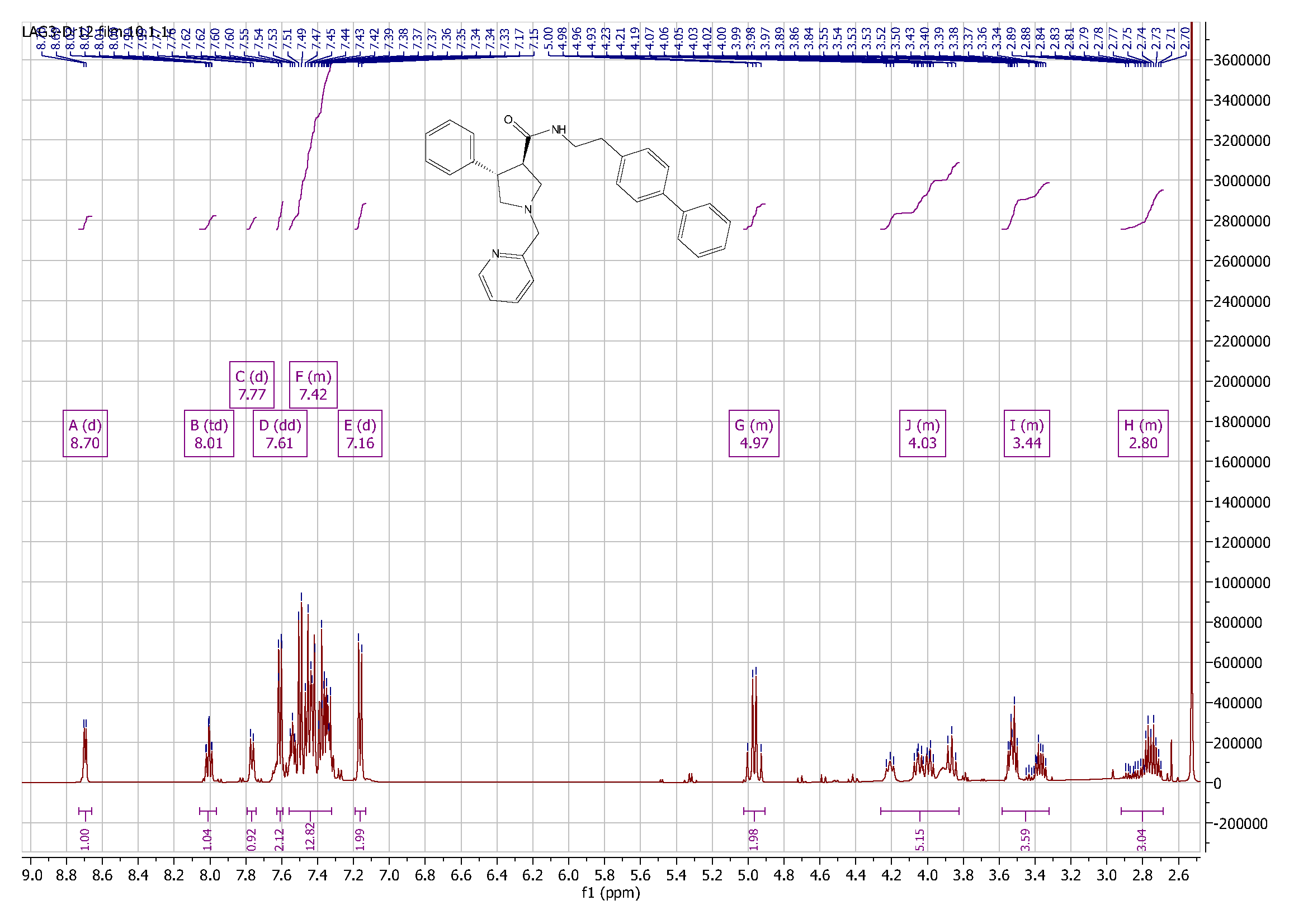

**Figure S38**. ^1^H NMR spectrum of compound **9**.

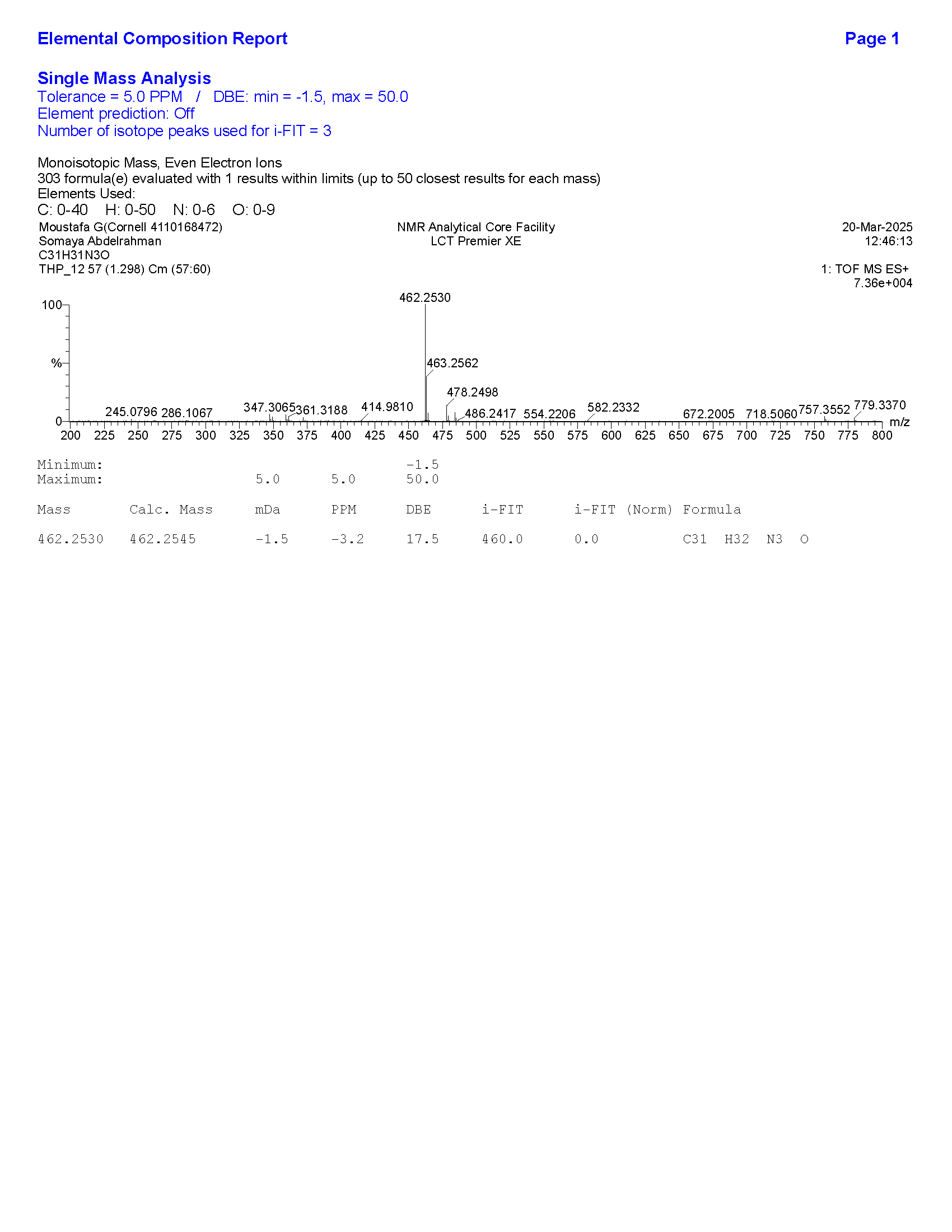

**Figure S39**. HRMS data for compound **9**.

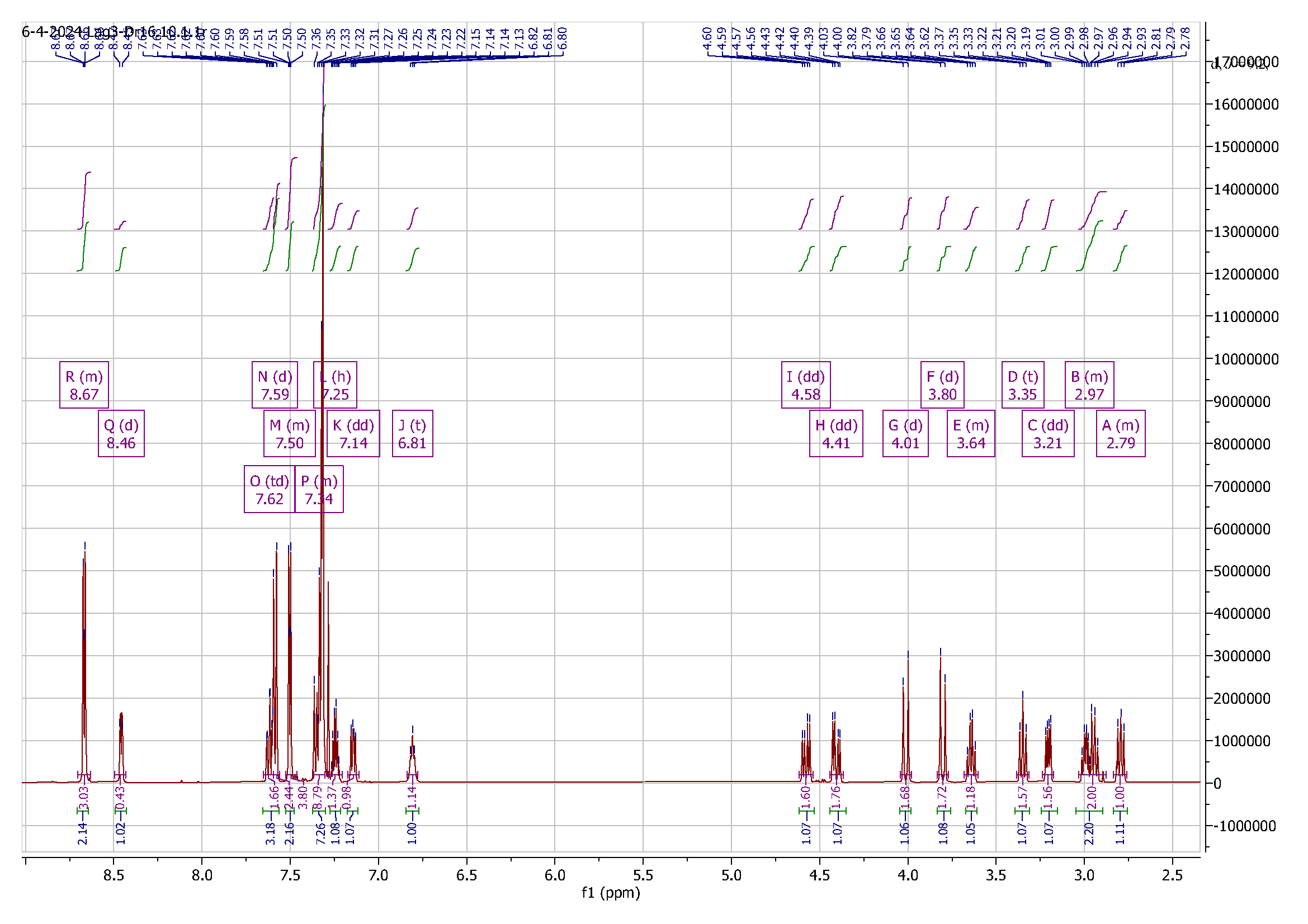
 **Figure S40**. ^1^H NMR spectrum of compound **10**.

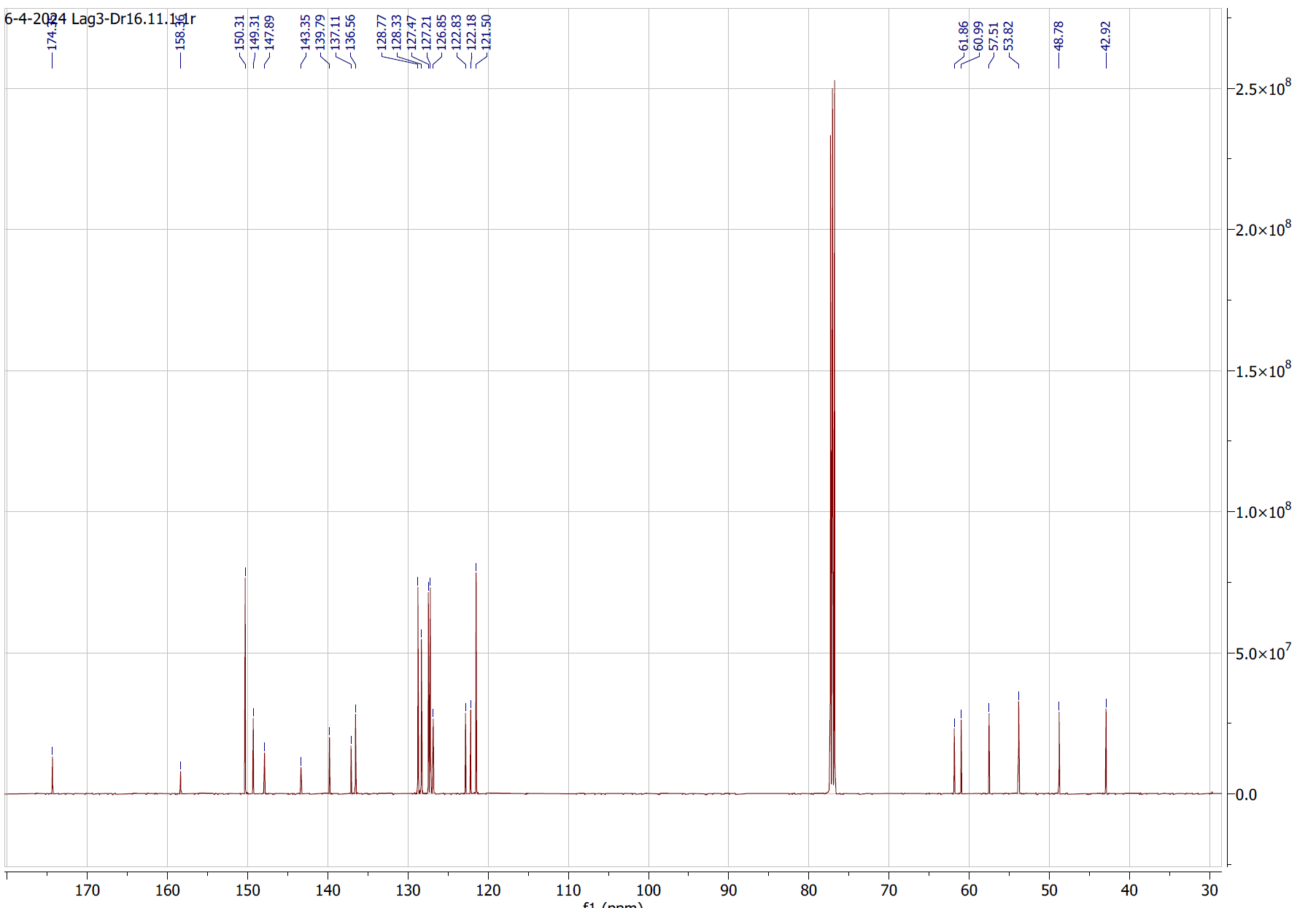

**Figure S41**. ^13^C NMR spectrum of compound **10**.

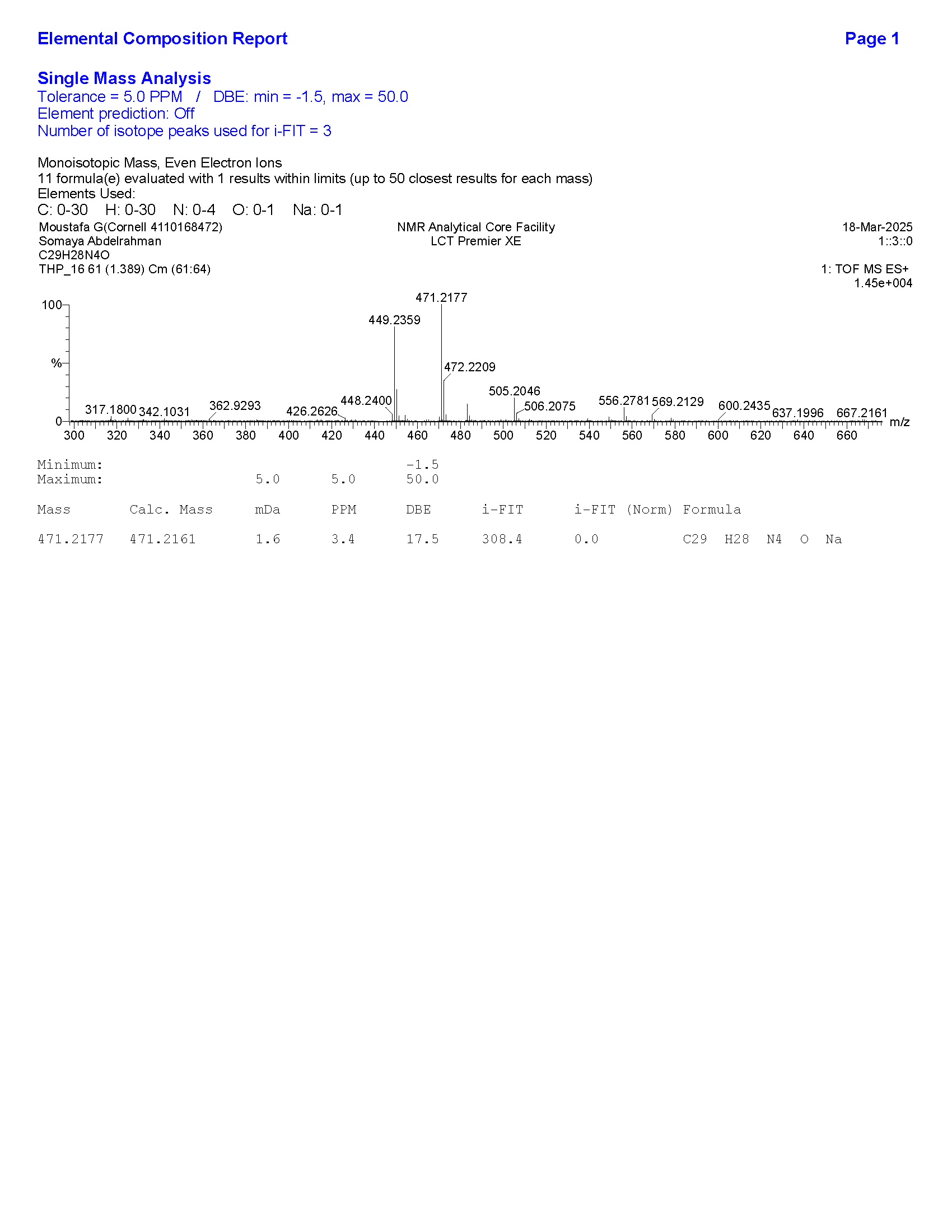

**Figure S42**. HRMS data for compound **10**.

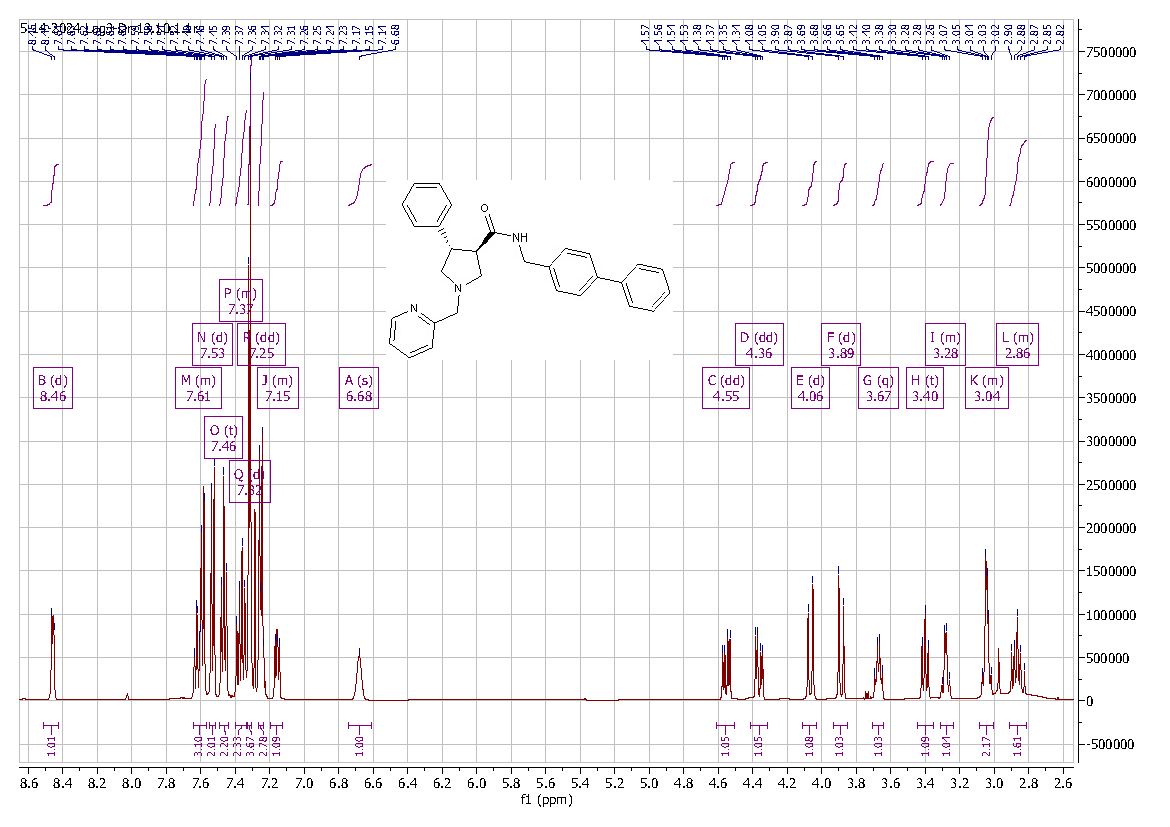

**Figure S43**. ^1^H NMR spectrum of compound **11**.

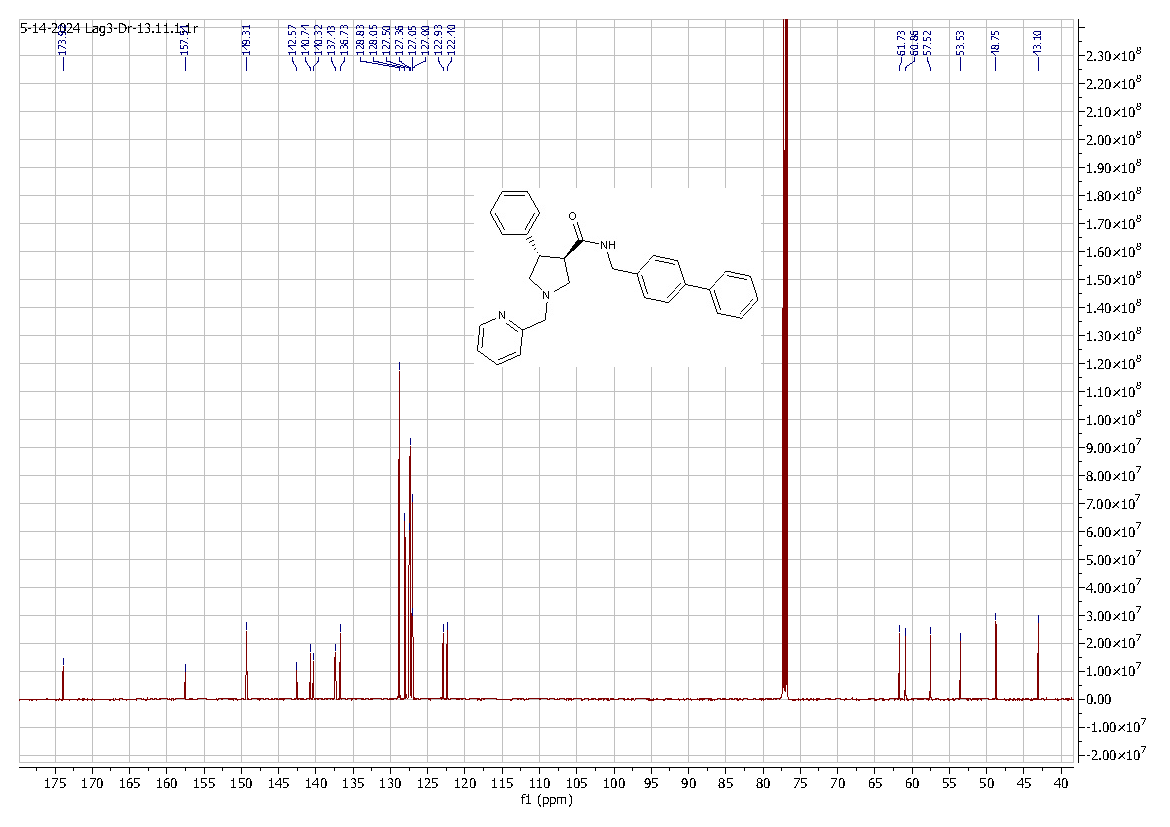

**Figure S44**. ^13^C NMR spectrum of compound **11**.

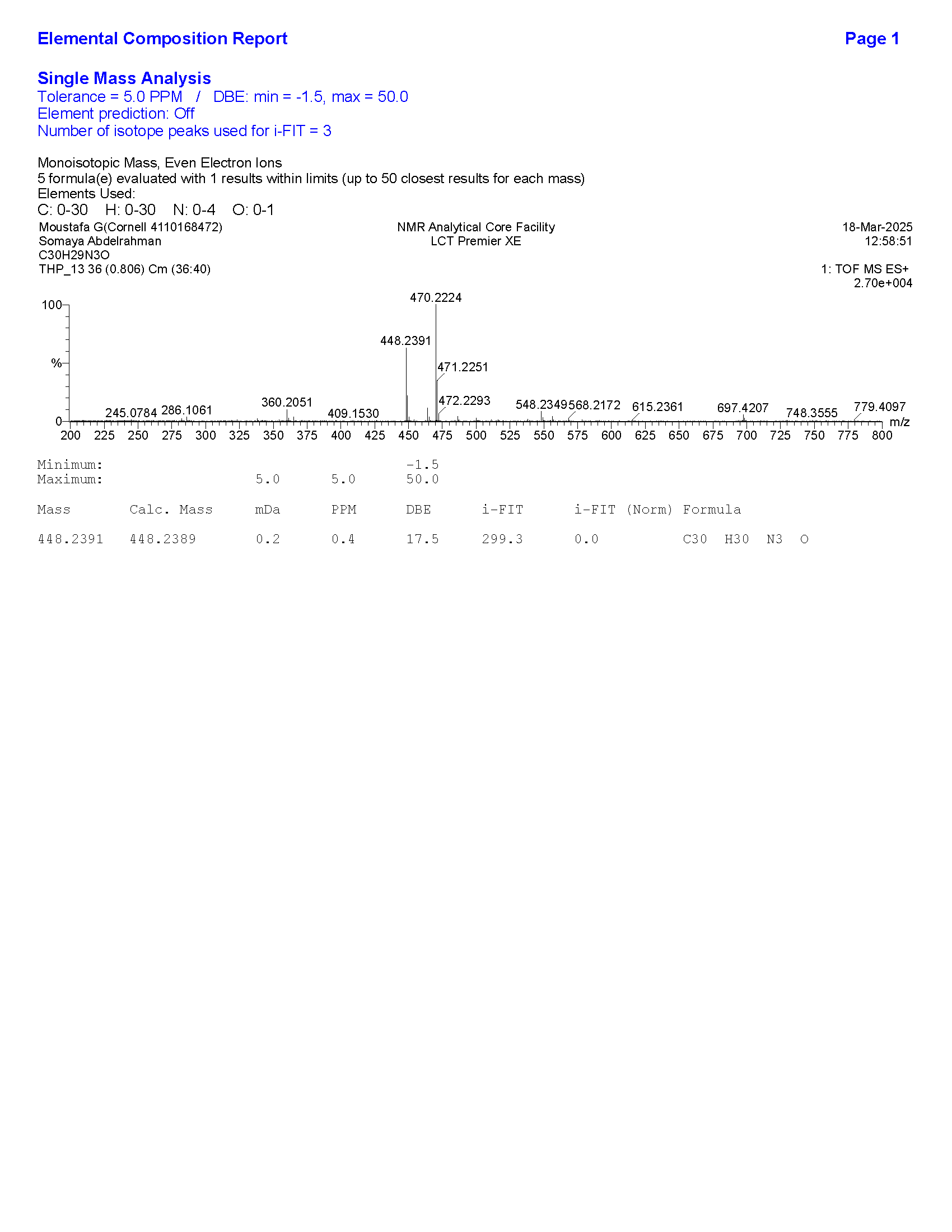

**Figure S45**. HRMS data for compound **11**.

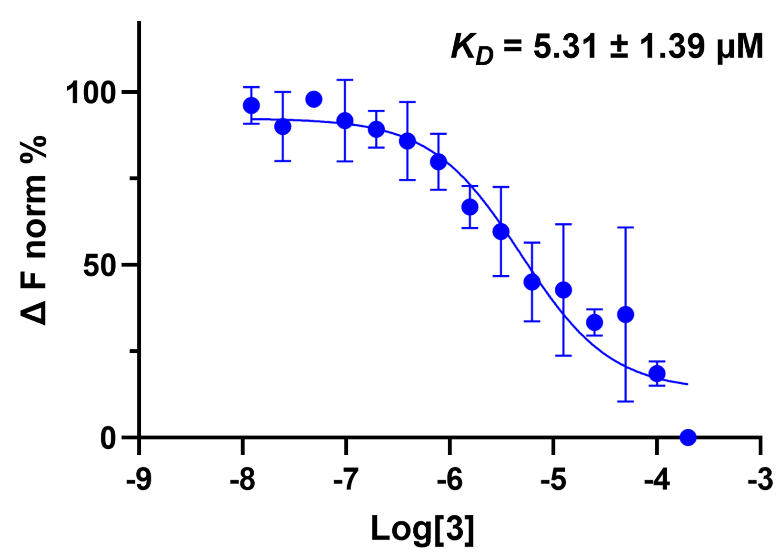

**Figure S46**. MST binding of compound **3** (increasing concentrations, n=3) to LAG-3.

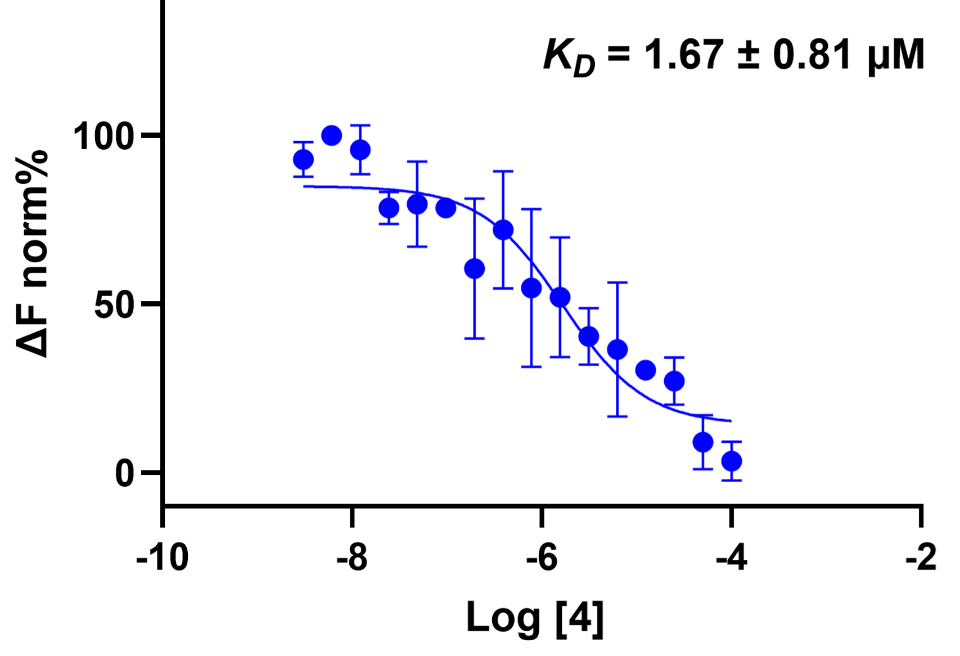

**Figure S47**. MST binding of compound **4** (increasing concentrations, n=3) to LAG-3.

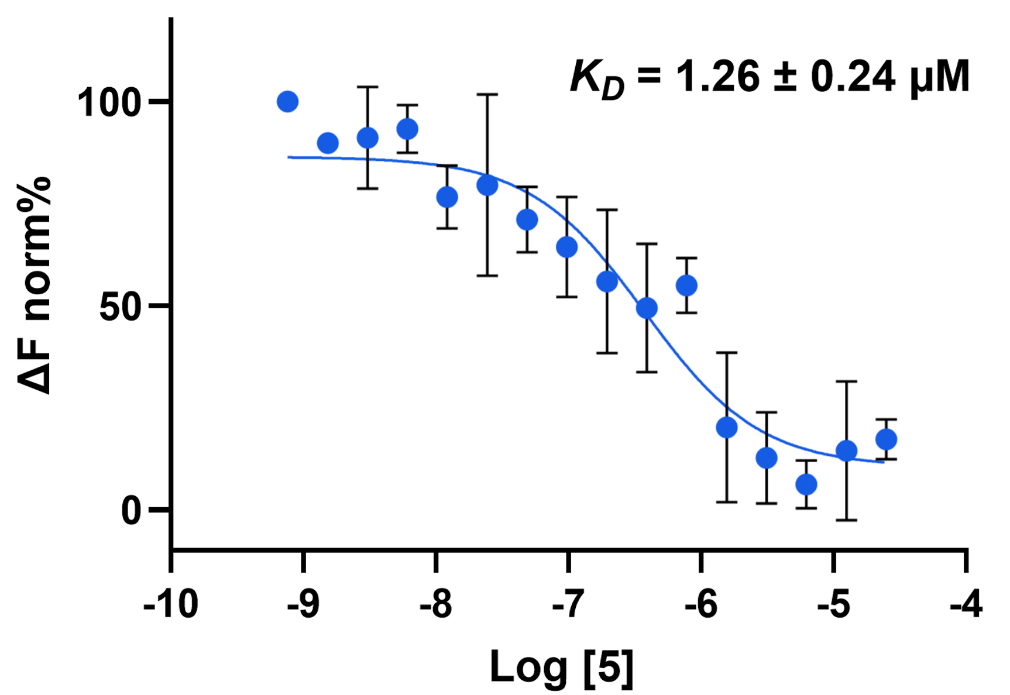

**Figure S48**. MST binding of compound **5** (increasing concentrations, n=3) to LAG-3.

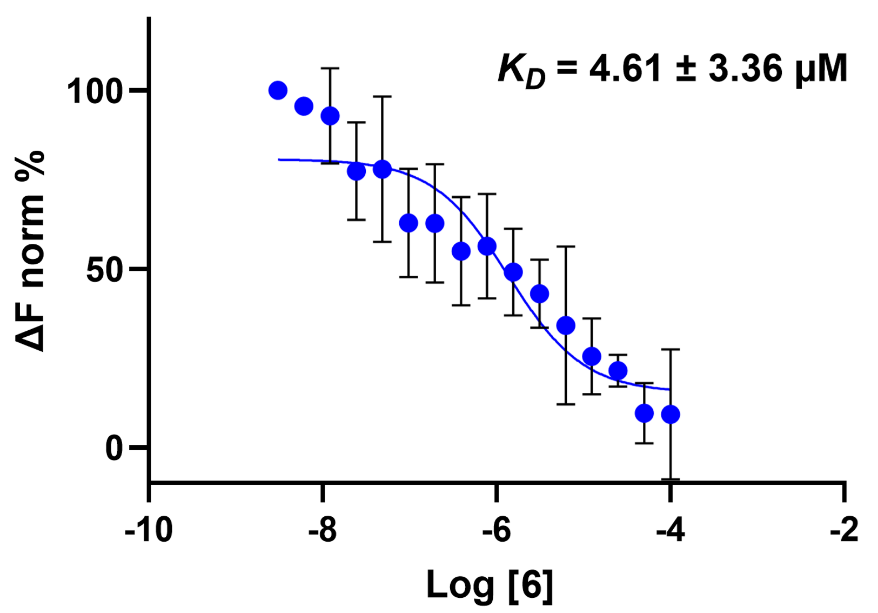

**Figure S49**. MST binding of compound **6** (increasing concentrations, n=3) to LAG-3.

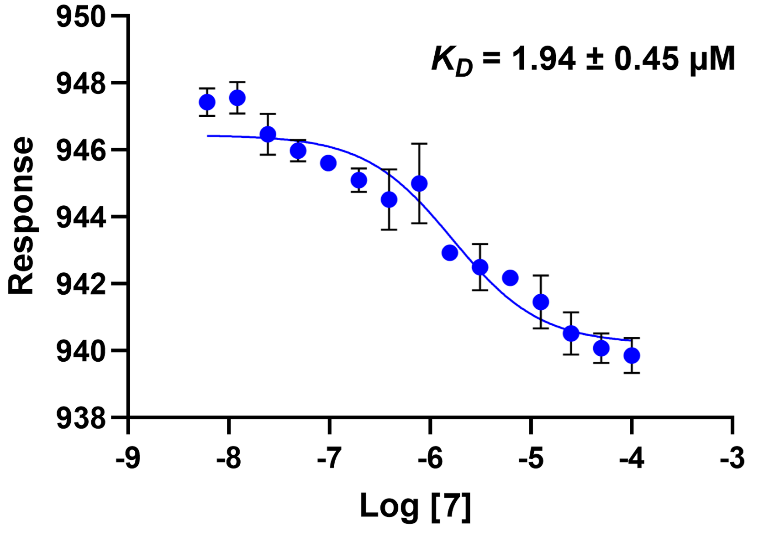

**Figure S50**. MST binding of compound **7** (increasing concentrations, n=3) to LAG-3.

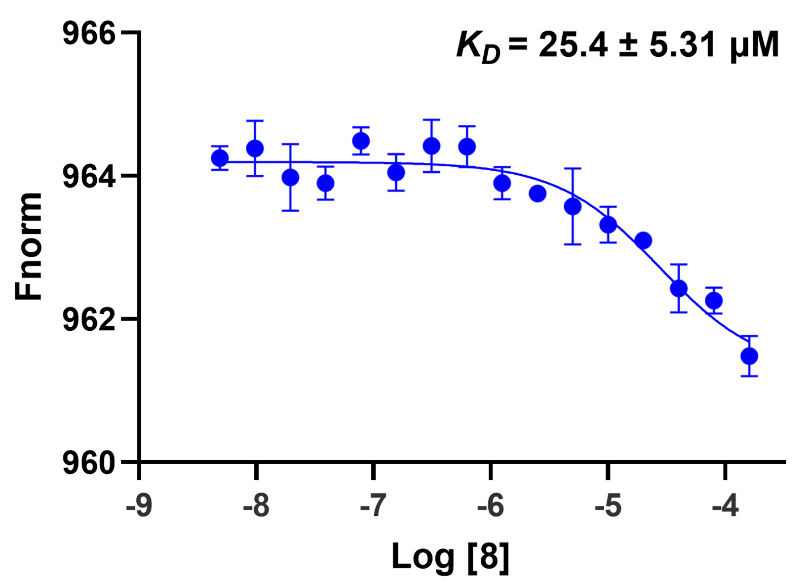

**Figure S51**. MST binding of compound **8** (increasing concentrations, n=3) to LAG-3.

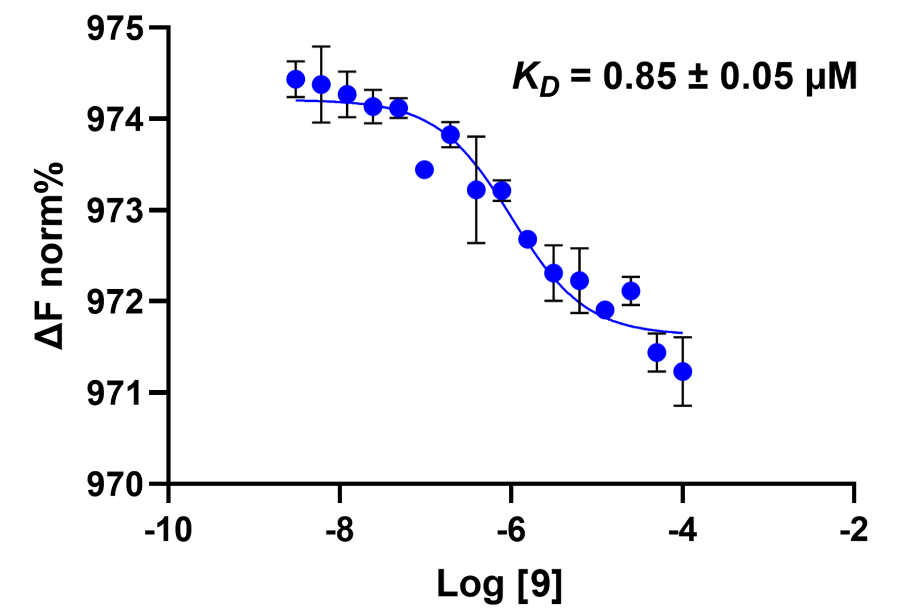

**Figure S52**. MST binding of compound **9** (increasing concentrations, n=3) to LAG-3.

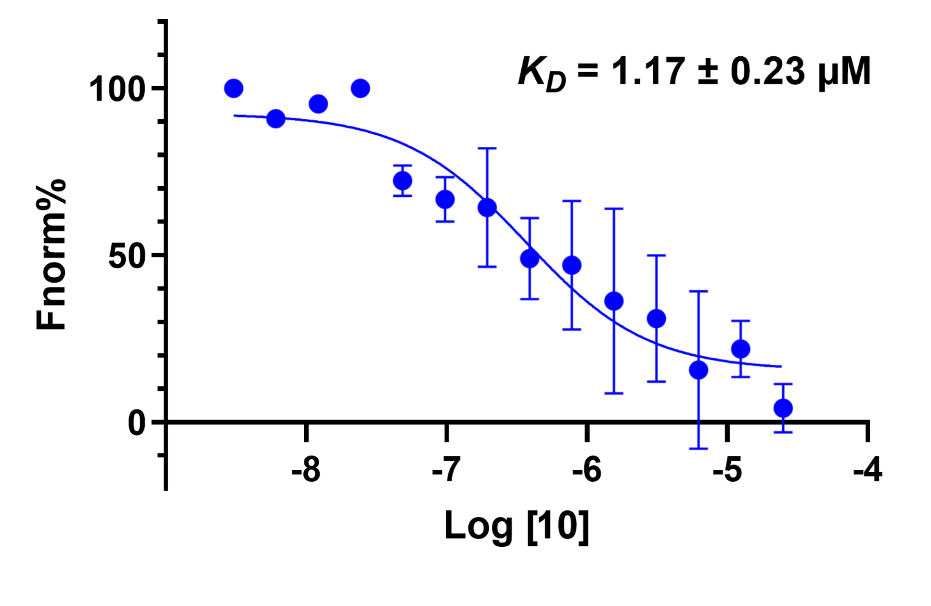

**Figure S53**. MST binding of compound **10** (increasing concentrations, n=3) to LAG-3.

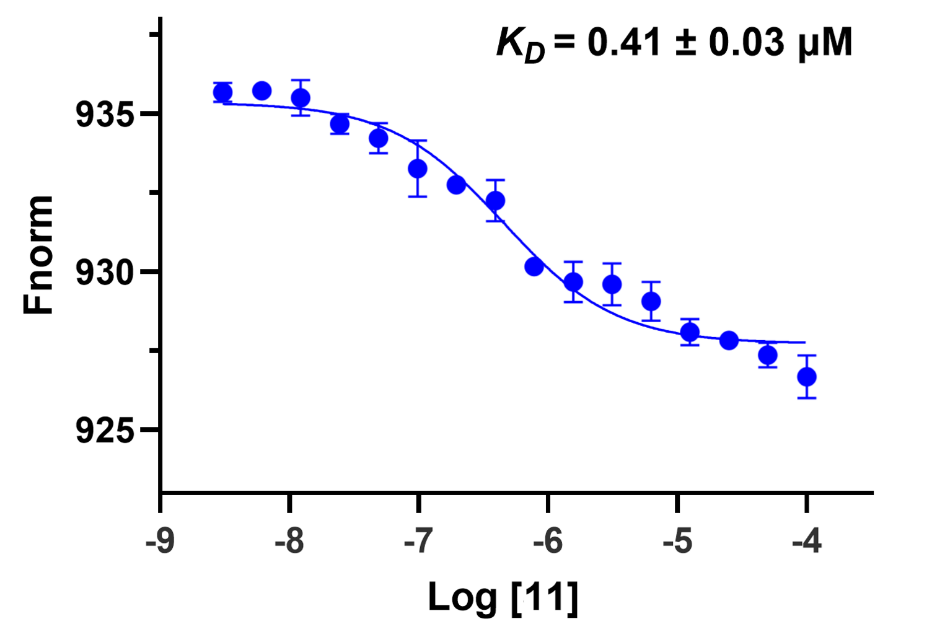

**Figure S54**. MST binding of compound **11** (increasing concentrations, n=3) to LAG-3.

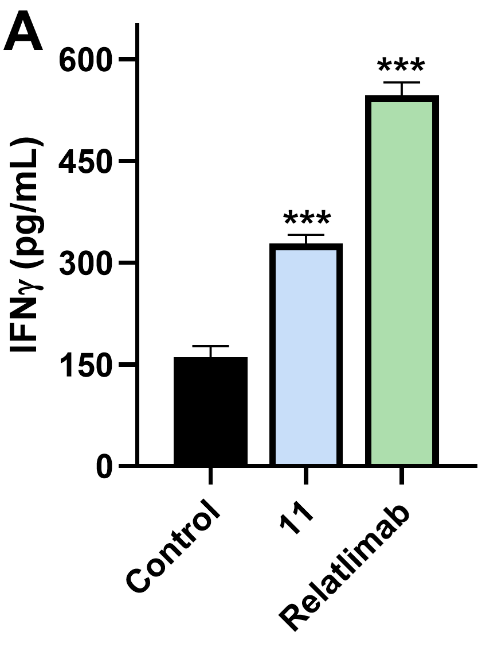

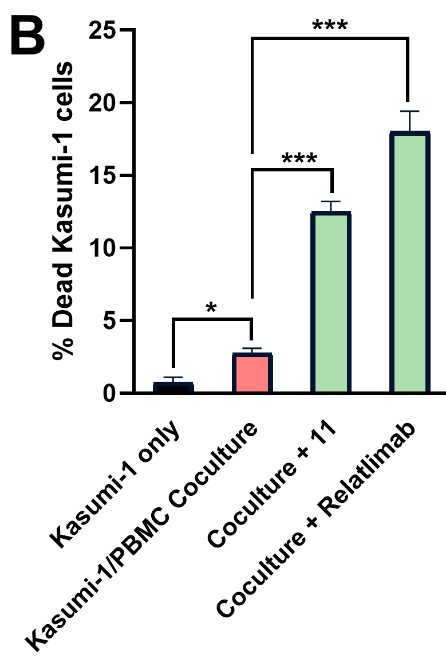

**Figure S55. A.** Production of IFNγ from PBMCs upon co-culturing with Kasumi-1 cells in the absence and presence of relatilmab (100 µg/ml) and compound **11** (10 μM), *** *p* < 0.001 in comparison to control. **B.** The % of dead Kasumi-1 cells as assessed by 7-AAD/CFSE assay in the co-culture assay of PBMCs and Kasumi-1 in the absence and presence of relatilmab (100 µg/ml) and compound **11** (10 μM), * *p* < 0.05, and *** *p* < 0.001. Error bars represent standard deviation (n = 3).

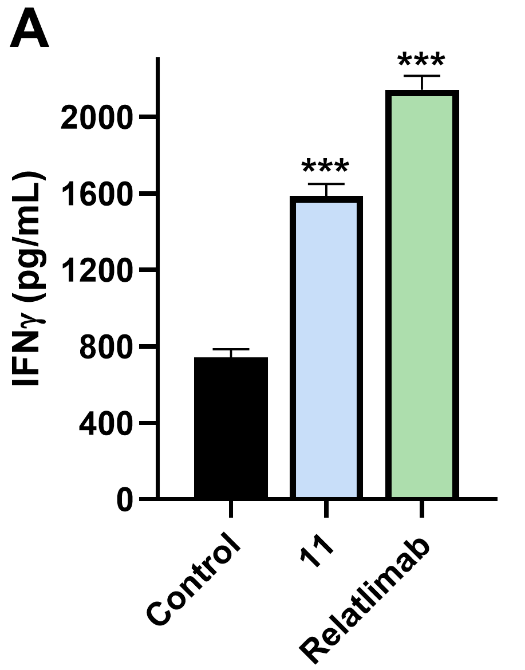

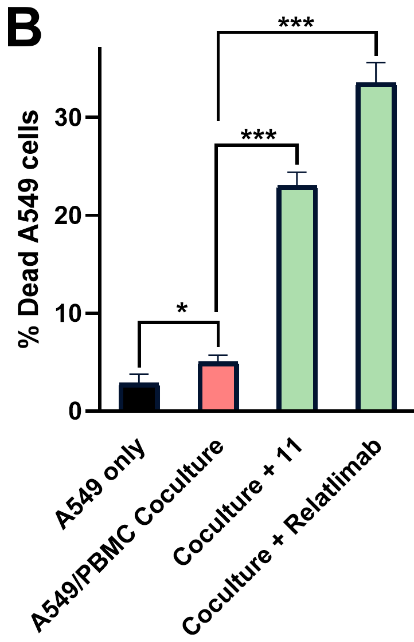

**Figure S56. A.** Production of IFNγ from PBMCs upon co-culturing with A549 cells in the absence and presence of relatilmab (100 µg/ml) and compound **11** (10 μM), *** *p* < 0.001 in comparison to control. **B.** The % of dead A549 cells as assessed by 7-AAD/CFSE assay in the co-culture assay of PBMCs and A549 in the absence and presence of relatilmab (100 µg/ml) and compound **11** (10 μM), * *p* < 0.05, and *** *p* < 0.001. Error bars represent standard deviation (n = 3).

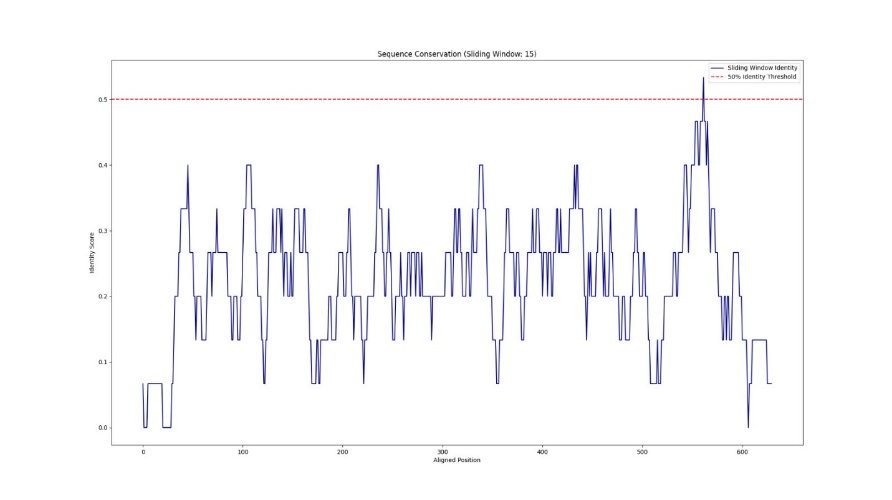

**Figure S57.** Sequence conservation diagram comparing mouse and human LAG-3.

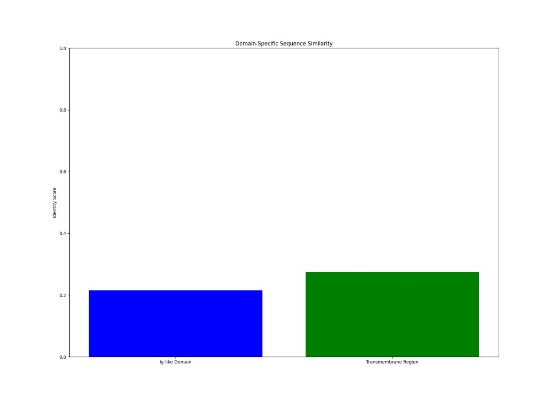

**Figure S58.** Sequence similarity diagram comparing mouse and human LAG-3.

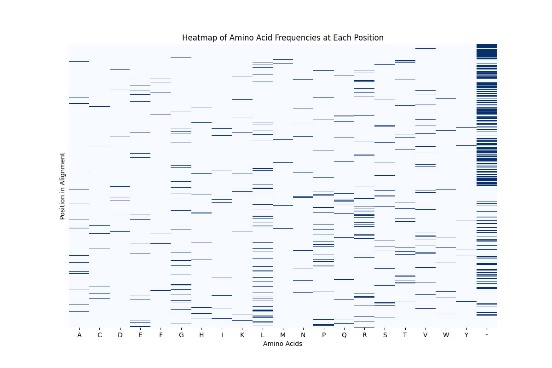

**Figure S59.** Heatmap of amino acids at each position obtained by aligning mouse and human LAG-3 sequences.

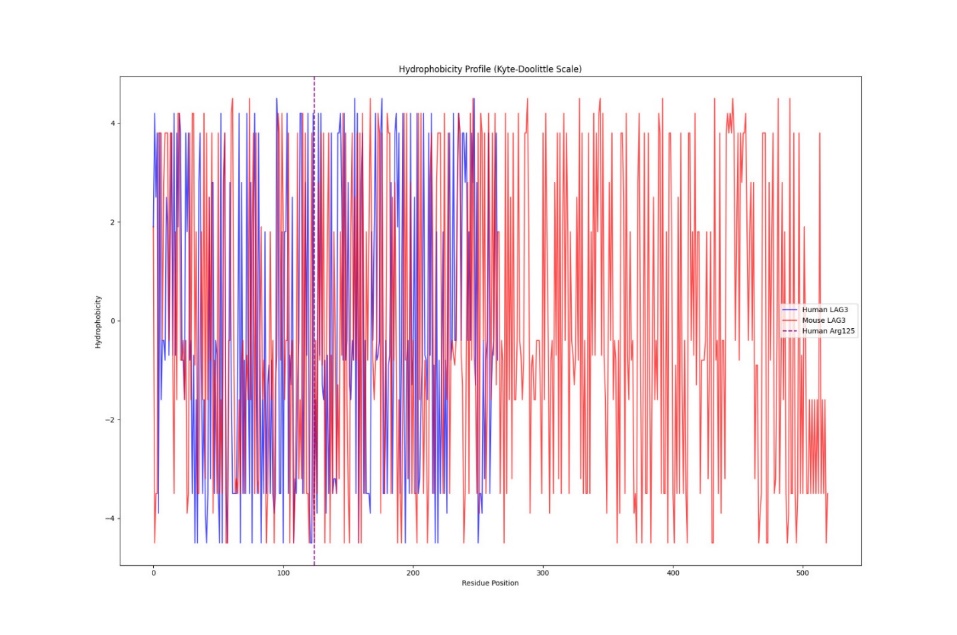

**Figure S60.** Hydrophobicity profile comparing mouse and human LAG-3 sequences.

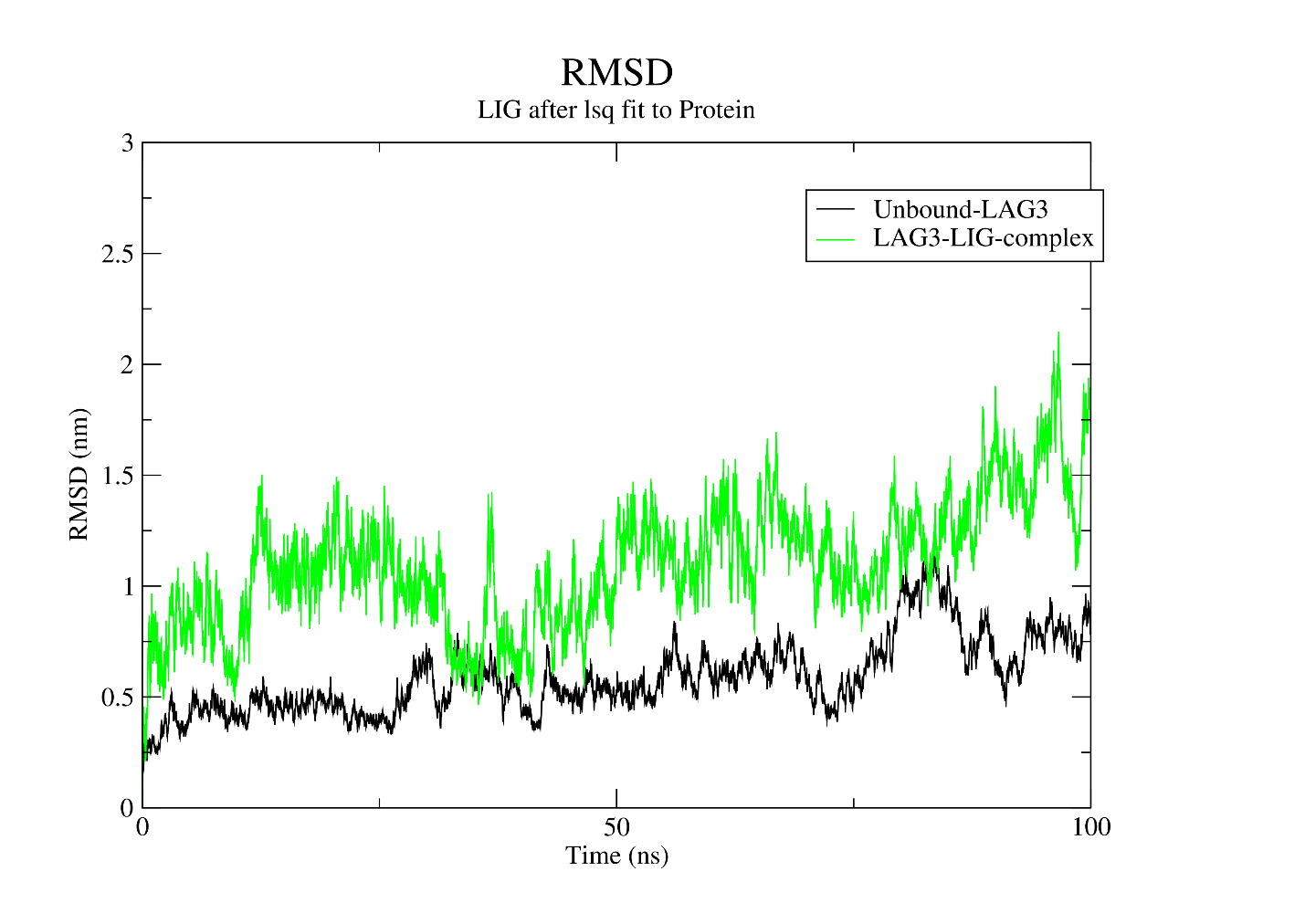

**Figure S61.** RMSD calculations for the unbound LAG-3 and LAG-3/compound **11** complex.

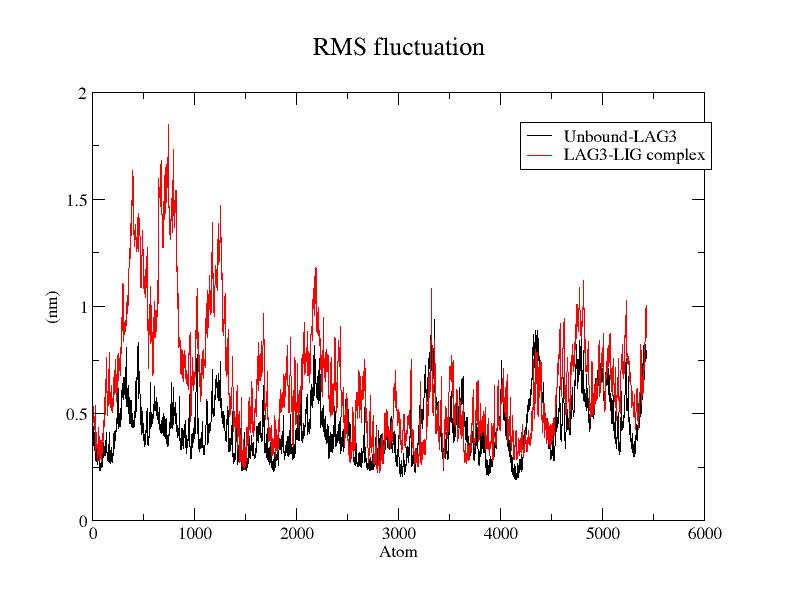

**Figure S62.** RMSF calculations for the unbound LAG-3 and LAG-3/ compound **11** complex.

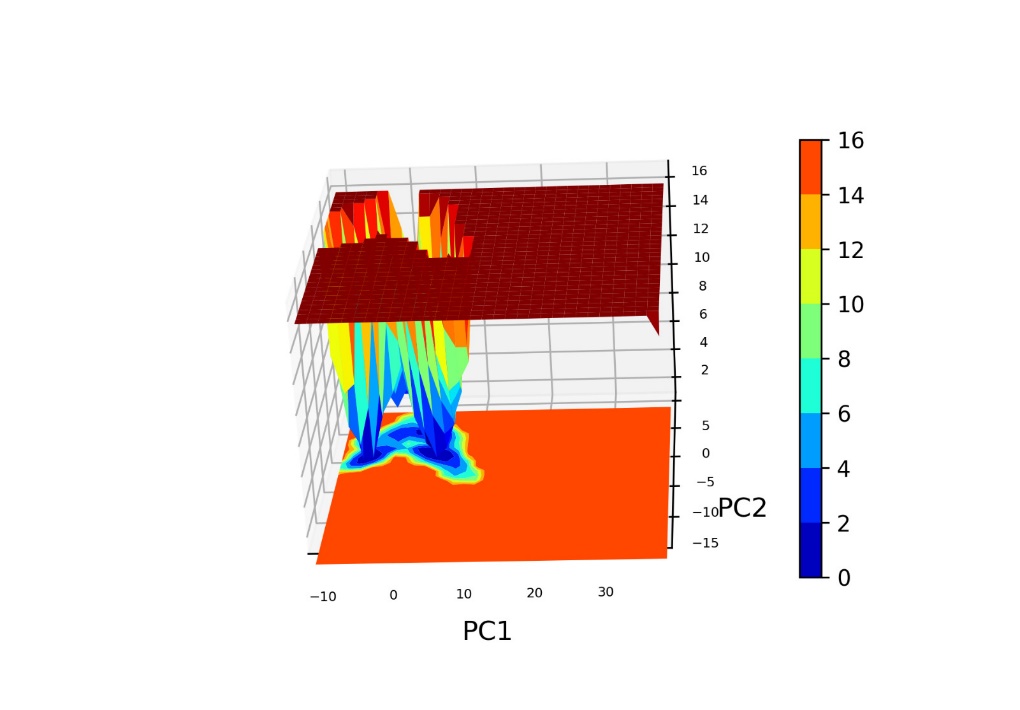

**Figure S63.** Gibbs free energy diagram of LAG-3/ compound **11** complex.

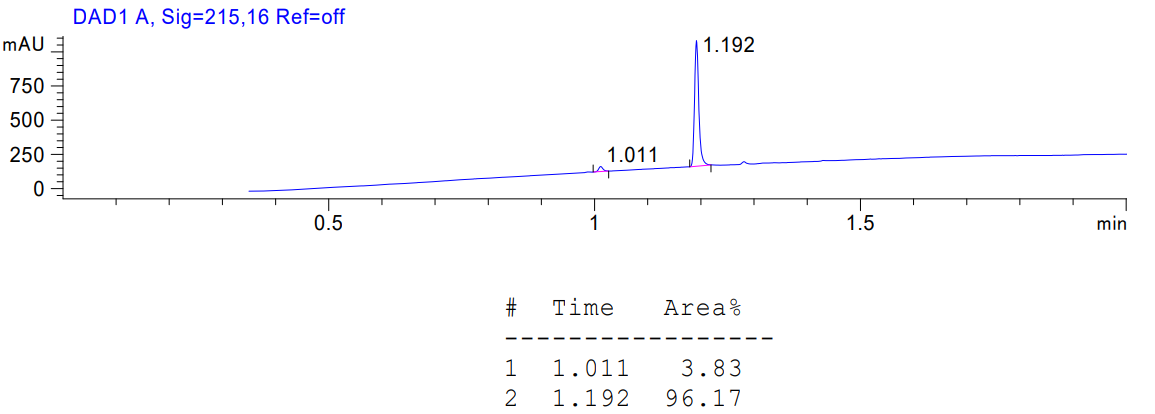

**Figure S64**. HPLC trace of compound **11**.
